# Synthesis of Dendritic Cell-Targeted Polymeric Nanoparticles for Selective Delivery of mRNA Vaccines to Elicit Enhanced Immune Responses

**DOI:** 10.1101/2023.11.13.566827

**Authors:** Chen-Yo Fan, Szu-Wen Wang, Cinya Chung, Chia-Yen Chen, Chia-Yen Chang, Yu-Chen Chen, Tsui-Ling Hsu, Ting-Jen R. Cheng, Chi-Huey Wong

## Abstract

Recent development of SARS-CoV-2 spike mRNA vaccines to control the pandemic is a breakthrough in the field of vaccine development. mRNA vaccines are generally formulated with lipid nanoparticles (LNPs) which are composed of several lipids with specific ratios; however, they generally lack selective delivery. To develop a simpler method selective delivery of mRNA, we reported here the synthesis of biodegradable copolymers decorated with guanidine and zwitterionic groups and an aryl-trimannoside ligand as polymeric nanoparticles (PNPs) for encapsulation and selective delivery of an mRNA to dendritic cells (DCs). A representative DC-targeted SARS-CoV-2 spike mRNA-PNP vaccine was shown to elicit a stronger protective immune response in mice as compared to the mRNA-LNP and mRNA-PNP vaccines without the selective delivery design. It is anticipated that this technology will be generally applicable to development of DC-targeted mRNA vaccines with enhanced immune response.

**TOC:** 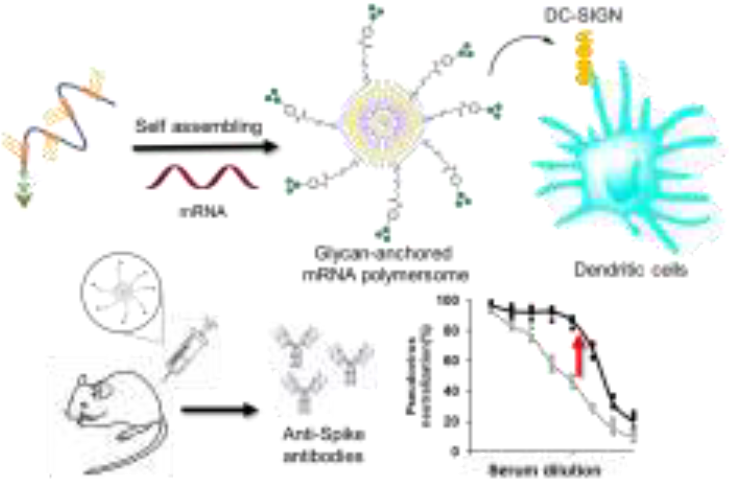

Dendritic cell-targeted mRNA-PNP vaccines

---

The global pandemic caused by the outbreak of severe acute respiratory syndrome coronavirus-2 (SARS-CoV-2) in 2019 has infected more than 750 million people and caused more than 7 million deaths. Since then, a great deal of global efforts has been made to develop safe, effective, and scalable vaccines to contain the pandemic. There have been more than 400 vaccines investigated to target SARS-CoV-2,^1^ of which the development of spike mRNA vaccines represents a major breakthrough in the field.^2-3^ After immunization, the mRNA of spike protein is translated to the spike protein *in vivo* to elicit immune responses, and there is no need to manufacture the spike protein *in vitro* for administration. This breakthrough in mRNA vaccine development has stimulated new research activities to further improve the technology. The drawbacks of using mRNA to replace protein include: it is difficult to conduct the post-translational modification of protein at the mRNA level; mRNA is generally unstable and requires low temperatures (−70 °C) for storage; mRNA must be modified and encapsulated in lipid nanoparticles (LNPs) for administration;^4-7^ the lipids used in the mRNA formulation are often mixed in specific ratios to form LNPs which are generally lack of selective delivery design and may require microfluidic device^8^; and the fate of mRNA after delivery is not well-understood and may cause undesirable side effects.^9^ One of our interests in the field is to develop a simpler method for encapsulation of mRNA vaccines and selective delivery to antigen presenting cells to elicit enhanced broadly protective immune responses.

Recently, polymers with disulfide linkages have been used as functional materials for a wide range of applications.^10-12^ Due to their dynamic and reversible properties, these polymers have been used for intracellular delivery of various cargoes.^13-14^ Disulfide exchange with exposed thiols followed by release of cargoes after uptake is achieved by reduction in the cytosol.^15-16^ The polymers with disulfide linkages are not only biodegradable in the intracellular environment, but also capable of thiol-mediated uptake, making possible efficient protein, siRNA, and mRNA delivery to the cytosol.^17^ In previous studies, a guanidine-containing polymer was demonstrated to promote the delivery of RNA-based therapeutics.^18^ However, the disadvantages of these polymer-drug complexes are relatively low efficacy *in vivo* due to the problems of serum instability and covalent cargo conjugation.^12, 19-20^

Despite advances in polymer synthesis,^21-23^ there are still lack of efficient nanocarriers for selective delivery of mRNA vaccine to antigen presenting cells,^24-25^ though an siRNA-liposome conjugated with a construct of trimeric N-acetyl-galactosamine (GalNAc) for targeting the asialoglycoprotein receptor on hepatocytes has been developed and approved by the FDA for the treatment of a liver disease. ^26-28^

In this study, we intended to identify a single polymer as mRNA carrier and we started with a thiol-initiator to react with different propagators to generate a series of different polymers for mRNA encapsulation to generate an mRNA-polymer nanoparticle (mRNA-PNP) complex for evaluation of immune responses and antibody neutralization activity. We finally identify a new type of polymers containing guanidine and zwitterionic groups with incorporation of a sugar ligand designed to target the DC-SIGN receptor for encapsulation and selective delivery of SARS-CoV-2 spike mRNA vaccine to dendritic cells to elicit enhanced immune responses.

We first designed a series of polymers and explored their ability to deliver the mRNA of green fluorescent protein (GFP) for transfection study and spike mRNA for vaccine development. As shown in figure 1, the propagator P1 containing a guanidine group and the propagator P2 containing three guanidine groups were designed to facilitate the encapsulation of mRNA by forming strong salt bridges between the guanidinium groups of the polymer and the phosphate groups of mRNA.^24^ The zwitterions were designed to enhance cell membrane fusion to facilitate uptake. We envisioned that once the polymer-encapsulated mRNA reaches the cytoplasm, the disulfide linkage will be degraded by intracellular glutathione to release mRNA to avoid the accumulation of high molecular weight polymers inside the cell to cause cytotoxicity.

**Figure 1.**
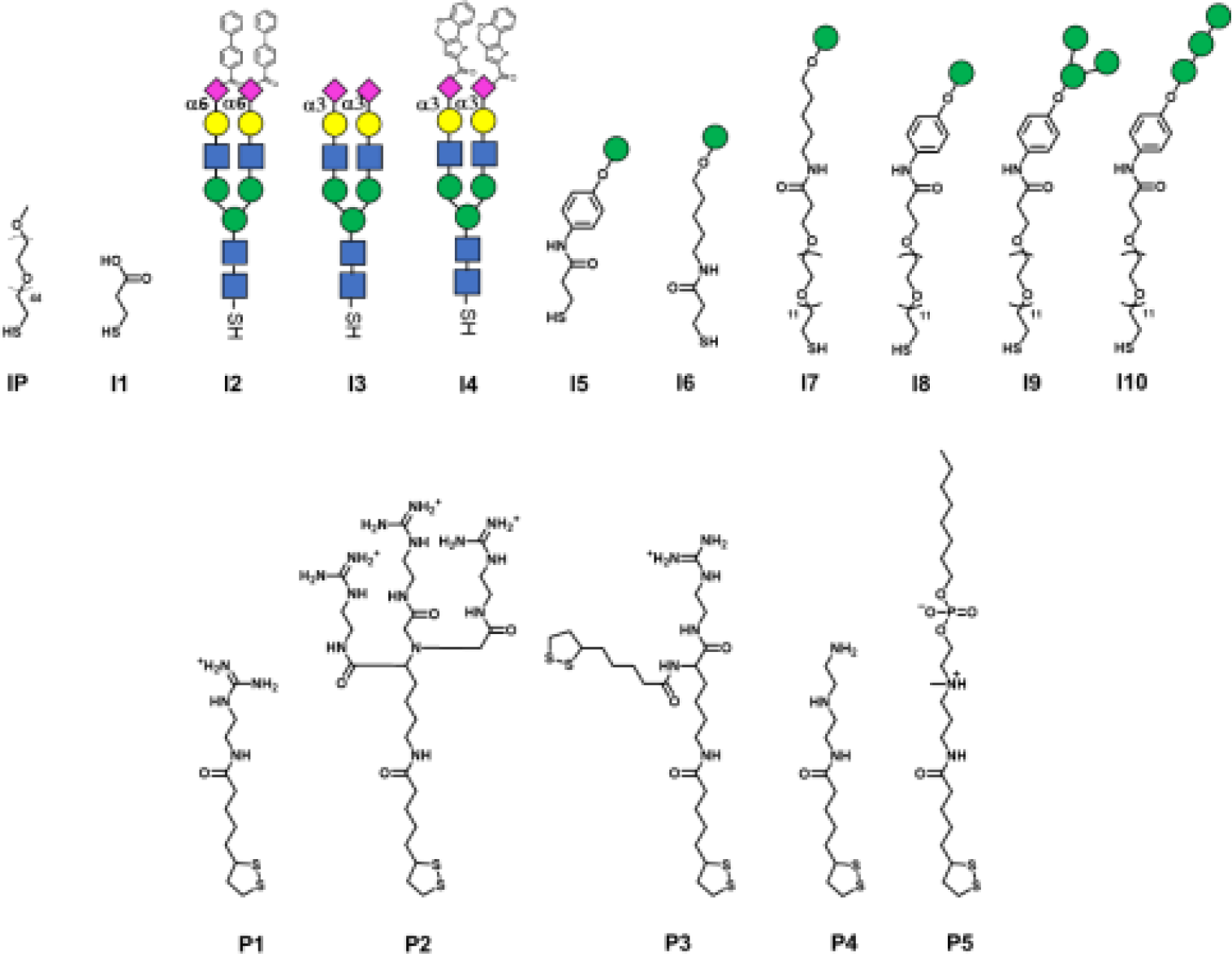
Designed structures of initiators (IP, I1, I2, I3, I4, I5, I6, I7, I8, I9 and I10) and propagators (P1, P2, P3, P4 and P5) for polymerization. Glycans that are recognized by antigen presenting cells are shown with specified color and shape symbols: blue square, GlcNAc; green circle, Man; yellow circle, Gal; pink diamond, Neu5Ac.

The mono-guanidine disulfide (P1) was synthesized according to the procedure reported^12^ and the tri-valent guanidine disulfide monomer was synthesized from nitrilotriacetic acid to generate the tri-valent guanidine monomer (see more synthetic details in SI). Propagator P3 containing two strained disulfides nearby the guanidine group was designed to form a branched structure of the polymer, whereas propagator P4 containing the diethylene-triamine moiety was designed to facilitate the escape of entrapped molecule from degradation and its terminal amine residue was used for functionalization. Propagator P5 containing a zwitterion group was intended to reduce serum protein adsorption and enhance membrane fusion, and enabled the delivery of mRNA from the nanocarrier to immune cells.^22^ The polymerization of P1, P2, P3, P4 or P5 was conducted in degassed water solution at room temperature. A mixture of initiator and propagators in 1 M pH 7 TEOA buffer was vigorously mixed for 30 min, and the reaction was terminated by adding 0.5 M iodoacetamide.

To identify the optimal polymer for efficient intracellular delivery of GFP-mRNA as a model, homo-polymers and hetero-copolymers were synthesized by co-polymerization of different propagators and their encapsulation ability and transfection efficiency in HEK293T cells were evaluated. As shown in figure S1, all the copolymers containing the guanidine group were able to encapsulate GFP-mRNA. Particularly, the P2/P3, P2/P4 and P2/P5 copolymers with trivalent guanidine moieties exhibited higher capability to encapsulate the mRNA and were comparable with the result with the traditional transfection agent polyethyleneimine (PEI). Next, we evaluated the transfection efficiency of GFP-mRNA in HEK293T cells by using different copolymers. First, we found that the mRNA encapsulated with a hetero polymer is important to achieve lysosomal escape and subsequent translation of the mRNA to proteins. The result indicated that P1 polymer was not favorable for intracellular delivery, while the hetero-polymer P1/P4, P2/P4, P1/P5, or P2/P5 was able to transfect HEK293T cells and release the mRNA for translation to GFP. Particularly, the polymer containing the zwitterionic group (Figure 2B and S2) was highly effective probably due to membrane fusion as described above. However, we found that the branched polymers P1/P3 and P2/P3 did not show a satisfactory release of mRNA cargoes, although a complete complexation with mRNA was achieved with P2/P3. These results indicated that the propagator is an important component in transfection and the co-polymers generated from P1/P5 and P2/P5 exhibited the highest transfection efficiency, and they did not exhibit any apparent cytotoxicity at the ratio of mRNA-Polymer = 1:3 (Figure S3).

**Figure 2.**
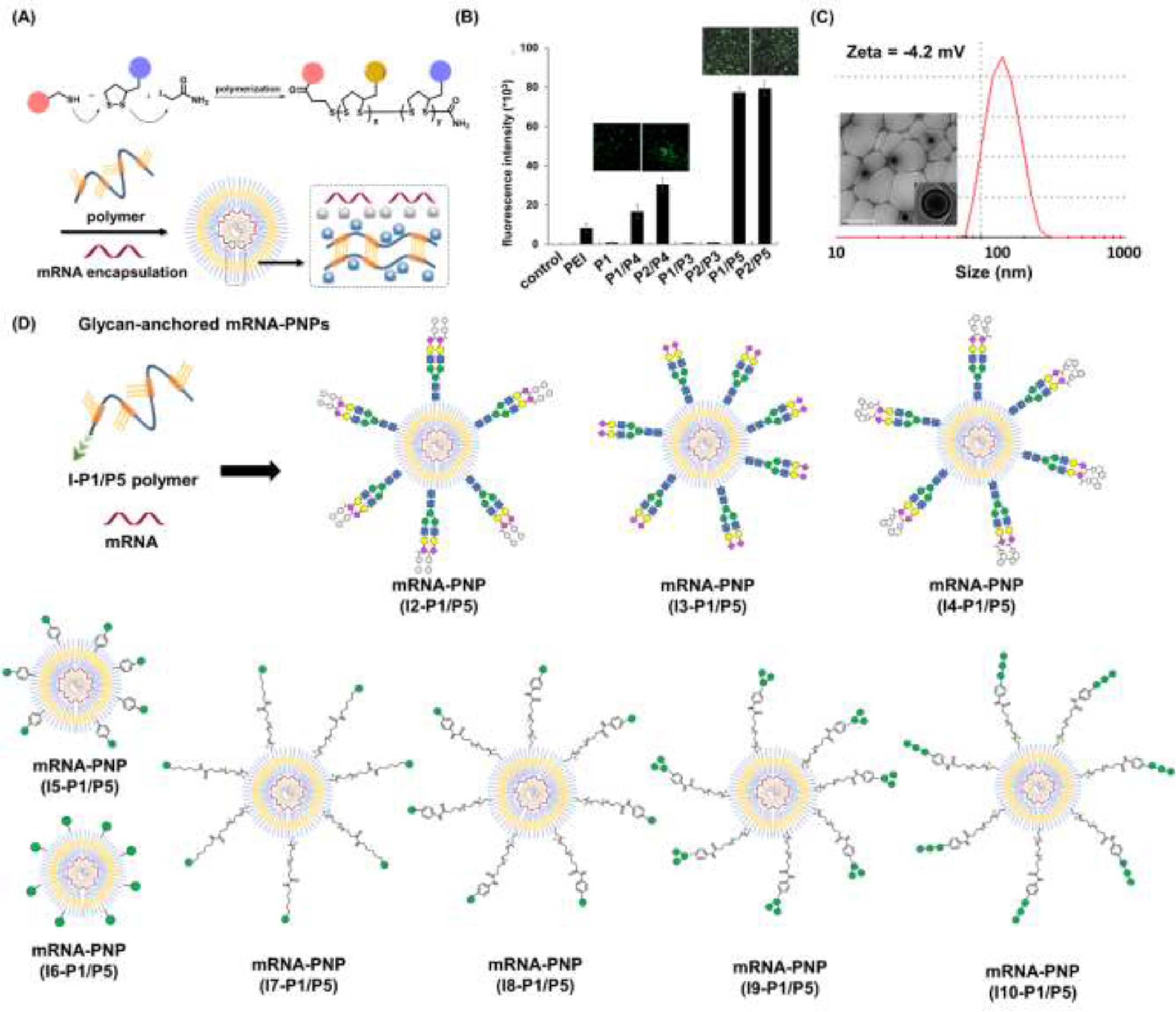
Encapsulation of mRNA with a designed polymer to form an mRNA-polymer nanoparticle (PNP) for delivery. (A) Synthesis of a polymer for mRNA encapsulation to form mRNA-PNP. (B) Mean fluorescence intensity of HEK293 cells after transfection with GFP mRNA-PNP (PNP: PEI, P1, P1/P4, P2/P4, P1/P3, P2/P3, P1/P5 and P2-P5). (C) Particle size and ζ potential of mRNA-PNP (P1/P5) with CryoEM image and insets. Scale bar represents 500 nm. (D) mRNA-PNP generated from P1/P5 and glycan initiator I2-I10 for evaluation of selective delivery to dendritic cells.

To examine the morphology of mRNA-polymer complex, we first mixed the P1 polymer with mRNA and it was found to form a nearly spherical electron-dense nanoparticle (Figure S4B), which is in general less efficacious because of serum instability.^29^. On the other hand, the complex of P1/P5 co-polymer with mRNA was found to be liposome-like (Figure 2C).^29^ The polymers are likely to form a layer with positive charge, which can encapsulate the mRNA, and facilitate subsequent cellular uptake through the engagement of the zwitterion (Figure 2A).

The molecular weight and polymerization index were characterized by gel permeation chromatography (GPC). As shown in Figure S5, the polymers displayed a unimodal but rather broad molecular weight distribution in the GPC chromatogram, and was eluted at the relatively low elution time, indicating its polymeric state. The peak molecular weight was 10.2 kDa (PDI = 1.33) for mRNA-PNP (P1/P5). Based on the result of various copolymers used for GFP-mRNA encapsulation, the P1/P5 co-polymer was selected for further studies, including its ability to complex with mRNA, the physiochemical properties of the resulting nanocomplexes, and the efficiency and selectivity of transfection.

Wild type spike mRNA was prepared and encapsulated by P1/P5 co-polymer at different N/P ratios (Figure S6). The result showed that the P1/P5 co-polymer exhibited a good ability to capture spike mRNA at the N/P ratio of 1, and the average particle size of the resulting mRNA-P1/P5 co-polymer complex was ca. 127 nm, as revealed by CryoEM, TEM and dynamic light scattering (DLS) analysis (Figure 2C and S4). To ensure the fully encapsulation of mRNA, we use a N/P ratio of 3 for study. The release and translation of mRNA may be attributed to GSH-mediated polymer degradation, as it takes place in the presence of 10 mM GSH in a time-dependent manner (Figure S7).

We then transfected the spike mRNA-P1/P5 complex (3 µg) in HEK293T cells and performed western blotting. After 48 h post transfection, cells were analyzed for spike-protein translation by western blotting with spike-specific antibody. A significant band at ∼250 kDa corresponding to SARS-CoV-2 spike protein was observed compared to the spike mRNA as negative control (Figure S8). This study confirmed that P1/P5 copolymer is an effective nanocarrier for mRNA transfection *in vitro*.

Zwitterionic lipids have been reported to improve serum resistance and cell membrane fusion.^25^ To visualize the location of mRNA-PNP in cells, an FITC-labelled polymer was synthesized from P1/P4/P5, in which FITC was conjugated to the polymer through the amine group on P4. The confocal fluorescence imaging confirmed that mRNA-PNP (I1-P1/P4/P5) exhibited effective mRNA-PNP escape from lysosomes within 2 h; however, the mRNA-PNP (I1-P1/P4) did not exit from lysosomes even in 4 h after internalization (Figure S9). As shown in Figure 3, the colocalization of lysosome with mRNA-PNP (I1-P1/P4/P5) was lower than the mRNA-PNP (I1-P1/P4), indicating that the alkylated zwitterion residue may significantly improve membrane fusion and lysosomal escape.

**Figure 3.**
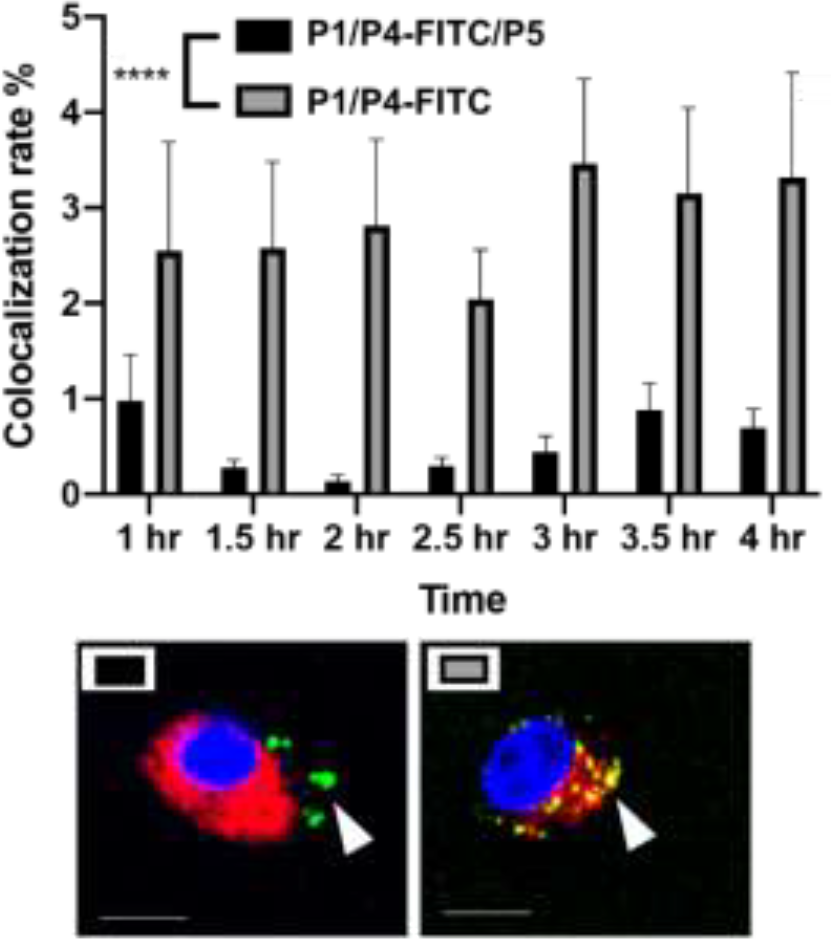
Colocalization rate of lysosome and mRNA-PNP by confocal fluorescence imaging with P1/P4/P5 or P1/P4 conjugated to FITC. The blue channel is DAPI, the red channel is lysotracker, and the green channel is mRNA-PNP-FITC.

In order to selectively deliver the mRNA vaccine to antigen-presenting cells (APCs) especially dendritic cells, we designed the initiator with different glycan heads recognized by the lectin receptors such as Siglec-1, Siglec-2, Siglec-5/E and DC-SIGN which are expressed predominately on DCs and macrophages.^30^ For Siglec-1 (sialoadhesin), 9-N-(4H-thieno[3,2-c]chromene-2-carbamoy-l)-Neu5Ac-α2,3-Gal-GlcNAc was found to be the most reactive ligand.^31^ Besides, 9-biphenyl Neu5Ac-α2,6-Gal derivatives showed an increased affinity for Siglec-2,^32^ and Siglec-5/E prefers the sialosides with the Neu5Ac-α2,3-Gal-GlcNAc structure.^33^ All ligands were synthesized and used as initiators for incorporation into the polymer which was evaluated for selective delivery of mRNA to dendritic cells.

I2-I5 initiated polymers were generated and I1 initiated polymer bearing no glycan ligand was served as control. Assuming the polymerization efficiencies of all the initiators are similar, polymers with fixed density of glycan ligands allowed us to examine how these functionalized polymers influence the cellular uptake of mRNA. To assess mRNA-PNP uptake *via* Siglecs, we compared the binding and internalization of mRNA-PNP into T cells, B cells, and bone marrow-derived dendritic cells (BMDCs). As evaluated by flow cytometry, FITC-conjugated mRNA-PNP were incubated with each cell line for 1 h. The I2 mRNA-PNP with 9^BPC^Neu5Ac conjugated N-glycan intended to target Siglec-2 showed a higher cellular uptake by all APCs compared to I1 mRNA-PNP without glycan modification (Figure S10). However, I2 mRNA-PNP did not show a selective uptake by dendritic cells. Similar results were obtained by using I3 and I4 mRNA-PNP to target Siglec-5/E and Siglec-1, respectively, in which all glycan-decorated mRNA-PNPs exhibited a better uptake by all APCs than the glycan-free mRNA-PNP but lack of selectivity toward DCs. With these discouraging results, we decided to evaluate the other C-type lectin DC-SIGN as targeting receptor.

DC-SIGN recognizes high mannose glycans and can route antigens for antigen processing to elicit robust immune responses, especially the CD4^+^ and CD8^+^ T cell responses,^34-36^ making it an attractive receptor for selective mRNA vaccine delivery. It has been used as a target for selective delivery of vaccines and mannose has been used as a ligand for such a delivery system.^37^ The initiator I5 with aryl mannoside was synthesized for mRNA-PNP preparation as it was shown to bind preferentially to DC-SIGN.^38^ The DC-SIGN-mediated cellular uptake by APCs was evaluated by flow cytometry at different time points (Figure S11). As shown in Figure S11B, the I5 mRNA-PNP with the aryl mannose head (I5-P1/P4-FITC/P5) exhibited ∼34% higher cellular uptake by BMDCs compared to the mRNA-PNP (I1-P1/P4-FITC/P5) without the glycan head. On the other hand, B cells and T cells with insignificant DC-SIGN expression showed only a slight increase of fluorescence signal when treated with mRNA-PNP (I1-P1/P4-FITC/P5). These data showed a proof of principle that efficient internalization and selective uptake of mRNA-PNP (I5-P1/P4-FITC/P5) by dendritic cells can be achieved through DC-SIGN receptor targeting.

Based on this approach, I6, I7, I8, I9, and I10 with different mannoside valency and spacer were synthesized and the corresponding mRNA-PNPs were prepared for study. I9 with aryl-α-1,3-α-1,6-trimannoside was first investigated, which contains a high-valent mannose and a hydrophobic presentation. Although the polymer with incorporation of mono-aryl mannose is likely to behave as multivalent mannose, the spatial arrangement of mannose would influence the specificity of receptor binding. Our comparative binding study has demonstrated that the linear trimannose ligand is more specific for mannose receptors (MRs),^39^ while the aryl tri-antennary mannose is prone to binding to DC-SIGN. Besides, antigen internalization and signaling rely on DC-SIGN engagement on the cell surface and in the endosome, and the pH of each environment differs. We therefore evaluated DC-SIGN binding to the mRNA-PNP generated from I5-P1/P5, I6-P1/P5, I7-P1/P5, I8-P1/P5, I9-P1/P5 and I10-P1/P5 under different pH values. We found that the mRNA-PNP from I5-P1/P5, I8-P1/P5, I9-P1/P5 and I10-P1/P5 bind to the DC-SIGN extracellular domain (ECD) at pH 7.4 and 5.0 with nearly the same affinity However, the mRNA-PNP generated from I6-P1/P5 and I7-P1/P5 lost their binding affinity at lower pH (Figures 4) to DC-SIGN probably because the coordination to calcium ions decreases at low pH values.^40^ This binding results indicate that aryl trimannoside interacts with DC-SIGN in the acidic endosomal compartments. Such binding stability would enhance DC-SIGN-mediated signaling and its synergism with that of endosomal-resident Toll-like receptors such as TLR7. The strong binding of PNP with aryl-mannoside to DC-SIGN may be due to the dense display of the ligand and the aryl group may engage in the CH−π and hydrophobic interactions.^41^ In addition, for receptor-targeting delivery, ligands are commonly linked with a carrier, and the distance between the carrier and ligand is regulated by the presence of a spacer.^42^ The mRNA-PNP carrying the longer length ligand (I6-P1/P5 with Man-Ar-PEG12, and I7-P1/P5 with Man-PEG12) showed slightly higher affinity toward DC-SIGN. Overall, the mRNA-PNP with aryl-trimannoside (I9-P1/P5) exhibited the highest affinity and lowest K_D_ to DC-SIGN. The increased affinity may be due to a clustering effect and the spatial arrangement of the ligand in the polymer, and the aryl moiety may facilitate its hydrophobic interaction. The aryl-trimannoside ligand and the resulting polymer were also characterized by ^1^H NMR (Figure S12-16), IR (Figure S17) and quantitative glycan analysis (Figure S18).

**Figure 4.**
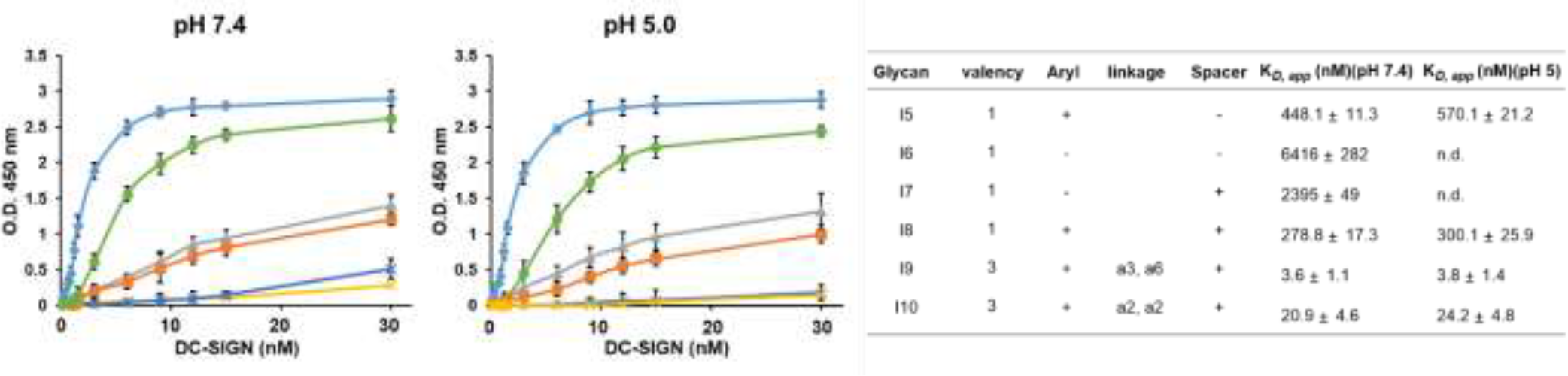
Enzyme-linked immunosorbent assay (ELISA) for measurement of dissociation constants in binding to DC-SIGN ECD oriented surface at pH 7.4 and pH 5.0: mRNA-PNP (I5-P1/P5) (orange), mRNA-PNP (I6-P1/P5) (yellow), mRNA-PNP (I7-P1/P5) (indigo), mRNA-PNP (I8-P1/P5) (grey), mRNA-PNP (I9-P1/P5) (blue), and mRNA-PNP (I10-P1/P5) (green). The error bars represent standard error of the mean from three independent experiments.

The preferential uptake of mRNA-PNP (I9-P1/P5) by BMDCs and C2C12 muscle cells was also evaluated by flow cytometry (Figure S19 and S20). As shown in Figure S19, FITC signals increased when BMDCs were incubated with mRNA-PNP (I9-P1/P5) especially with 1:1000 and 1:2000 dilutions. Figure S20 showed FITC signals at different incubation intervals and the ratios of I9-P1/P5 versus I1-P1/P5 signals indicated that the mRNA-PNP had a better uptake than the ones without glycans. On the other hand, when C2C12 muscle myoblasts cells, a cell line without mannose receptors were treated with mRNA-PNP (I9-P1/P5) or mRNA-PNP (I1-P1/P5) (1:1000 dilution) (Figure S21), both samples showed very similar FITC profiles (geometric mean ratio of I9-P1/P5 versus I1-P1/P5 = 100.9%). These data suggested that there is no selectivity towards C2C12 muscle cells.

We further tested whether the uptake of mannosylated mRNA-PNPs are dependent on DC-SIGN. A binding study using ELISA-based measurement including DC-SIGN, macrophage mannose receptor (MMR), MINCLE, Dectin-2, and langerin was conducted. As shown in Figure S22, we observed that the branched type of aryl-trimannoside **I9** showed a better preference to DC-SIGN compared to the linear type of aryl-trimannoside **I10**. In addition, **I10** was strongly bound to either DC-SIGN, MMR, Dectin-2 or langerin with no significant selectivity. However, **I9** presented a more selective binding toward DCSIGN. These data supported that the efficient uptake of mRNA-I9-P1/P5 by DCs is more likely mediated by DC-SIGN.

We next evaluated the effect of WT spike mRNA-PNP with or without the aryl mannoside head on vaccination and immune responses. BALB/c mice were immunized with the vaccines on day 0 and received a boost on day 14 (Figure 5A). As a positive control, a group treated with spike mRNA-LNP (including ALC-0315, DSPC, ALC-0159, and cholesterol commonly used in current mRNA vaccine formulation) was included. We observed a ten thousand level of anti-spike antibodies in the serum of the mRNA-PNP (I1-P1/P5) and mRNA-LNP groups (Figure 5B). However, the titer of spike-specific antibody response in mice after second immunizations with the aryl-trimannoside containing mRNA-PNP (I9-P1/P5) was 3.1-fold higher than the two control groups on days 28. The serum was then tested for the ability to neutralize pseudovirus-mediated entry to the ACE2-expressing cells. While the level of antisera from mRNA-PNP (I1-P1/P5) was similar to the mRNA-LNP group, a significantly higher level of neutralizing antibodies in the mRNA-PNP (I9-P1/P5) group was observed (Figure 6C), and the spike-specific antibody titer and pseudovirus neutralization activity were well correlated.

**Figure 5.**
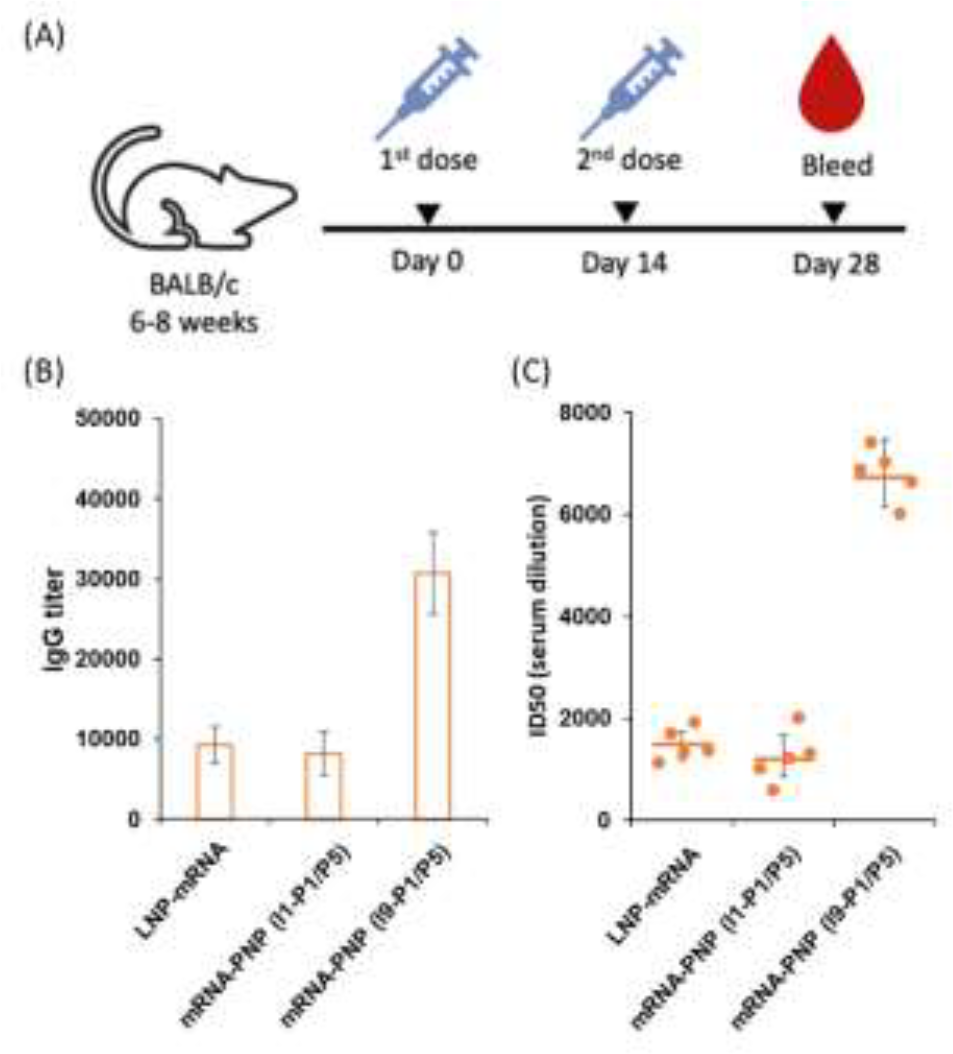
(A) BALB/c mice were immunized with 15 μg of WT spike, mRNA-LNP, mRNA-PNP (I1-P1/P5), and mRNA-PNP (I9-P1/P5), and boosted on day 14 after priming (n = 5) (B) Serum levels of spike-specific IgGs from spike mRNA-PNP were monitored over 28 days postpriming by ELISA. (C) Sera obtained from mice after the last immunization were evaluated for their neutralization activities against SARS-CoV-2 by a pseudovirus-based neutralization assay. Neutralization titers (ID50) were calculated as the reciprocal of the serum dilution that resulted in a 50% reduction in RLUs compared to virus control wells after subtraction of background RLU. The ID50 values are labeled on the plots with standard error of mean.

In conclusion, we have developed and investigated a series of polymers with different glycan ligands for mRNA vaccine delivery and demonstrated that the polymer containing guanidine and zwitterionic groups exhibited great efficiency for mRNA vaccine delivery both *in vitro* and *in vivo*. In addition, the polymer generated by initiator I9 with an aryl trimannoside head and propagators P1/P5 is an effective carrier for selective delivery of mRNA vaccines to dendritic cells for translation, processing, presentation, and efficient immune responses against the spike antigen. More detailed studies of CD4^+^ and CD8^+^ T-cell responses and further evaluation of the mRNA-PNP in animal challenge study are ongoing. We anticipate that this method is generally applicable to other mRNAs for vaccine design and therapeutical development.

## ASSOCIATED CONTENT

### Supporting Information

The Supporting Information is available free of charge at .

## Notes

The authors declare no competing financial interest.

## ACKNOWLEDGMENT

We thank the National RNAi Core Facility at Academia Sinica for pseudotyped lentivirus neutralization assay and related services. This research was supported by Academia Sinica.

