## Supplementary Information for "Synthesis of Dendritic Cell-Targeted Polymeric Nanoparticles for Selective Delivery of mRNA Vaccines to Elicit Enhanced Immune Responses"

Chen-Yo Fan,[a] Szu-Wen Wang,[a] Cinya Chung,[a] Chia-Yen Chen,[a] Chia-Yen Chang,[a] Yu-Chen Chen,[a] Tsui-Ling Hsu,[a] Ting-Jen R. Cheng,[a] and Chi-Huey Wong[a,b,*]
[a] Genomics Research Center, Academia Sinica, Taipei 115, Taiwan
[b] Department of Chemistry, The Scripps Research Institute, La Jolla, CA 92037

Corresponding Author: Chi-Huey Wong


### Table of contents

Materials and Methods
Experimetal Section
Scheme S1-S6
Synthesis of compound **2-54**
Figures S1-S22
$^{1}$H and $^{13}$C NMR of compound **2-54**

### Chemical Materials and Methods

For chemical synthesis, all starting materials and commercially obtained reagents were purchased from Sigma-Aldrich and used as received unless otherwise noted. All reactions were performed in oven-dried glassware under nitrogen atmosphere using dry solvents. $^{1}H$ and $^{13}C$ NMR spectra were recorded on Brucker AV-600 spectrometer, and were referenced to the solvent used ($CDCl_3$ at δ 7.24 and 77.23, $CD_3OD$ at δ 3.31 and 49.2, and $D_2O$ at δ 4.80, and DMSO-$d_6$ at δ 2.5 and 39.51 for $^{1}H$ and $^{13}C$, respectively). Chemical shifts (δ) are reported in ppm using the following convention: chemical shift, multiplicity (s = singlet, d = doublet, t = triplet, q = quartet, m = multiplet), integration, and coupling constants (*J*), with *J* reported in Hz. High-resolution mass spectra were recorded under ESI-TOF mass spectroscopy conditions. Silica gel (E, Merck) was used for flash chromatography. IMPACT™ system (Intein Mediated Purification with Affinity Chitinbinding Tag) was purchased from New England Biolabs. His-tag purification resin was purchased from Roche. HiTrap IMAC column (5 mL) was purchased from GE Healthcare Life Sciences. Gel permeation chromatography (GPC) equipped with Ultimate 3000 liquid chromatography associated with a 101 refractive index detector and Shodex columns was used to analyze the polymeric products using THF as the eluent at 30 °C with 1 mL $min^{-1}$ flow rate. The calibration was based on narrow linear poly(styrene) Shodex standard (SM-105). The Mw and dispersity of the polymeric products were calculated by DIONEX chromeleon software.Transmission electron microscopy (TEM) images were obtained by a FEI Tecnai G2 F20 S-Twin.

### Experimental Section

#### Cell Cultures

Human Embryonic Kidney Cells 293 (HEK293T cell) was cultured in Dulbecco's modified Eagle's medium (DMEM) (invitrogen, Carlsbad, CA, USA) supplemented with 10% fetal bovine serum (FBS) (Invitrogen) and 1% Gibco Antibiotic-Antimycotic (anti-anti) (Invitrogen).

#### Animals

Balb/c mice (8 weeks) were purchased from National Laboratory Animal Center, Taiwan. All the mice were maintained in a specific pathogen-free environment. Eight-week-old Balb/c mice were immunized i.m. twice with 2-week interval. Each vaccination contains in PBS (100 μl). Sera collected from immunized mice were subjected to ELISA analysis 10 days after the last immunization. The experimental protocol was approved by Academia Sinica's Institutional Animal Care and Utilization Committee (approval no. 22-08-1901).

**Antibodies and Proteins**

The SARS-CoV-2 full-length spike protein was purchased from ACROBiosystems. HRP conjugated anti-mouse secondary antibody and horseradish peroxidase substrate were purchased from Thermo Scientific. Mouse monoclonal anti–β-actin were purchased from Millipore. All commercial antibodies were validated for specificity by companies via Western blot.

**WT Spike DNA Construction**

pMRNA$^{XP}$ mRNA Synthesis Vector was obtained from System Biosciences. WT (Wuhan/WH01/2019 strain) spike DNA sequence with K986P and K987P mutations (2P) was codon-optimized for Homo sapiens.[1] pMRNA$^{XP}$ vector was digested with EcoRI and BamHI at 37 °C for 1 hour. DNA sequence of the spike protein was amplified by KOD One™ PCR master mix (TOYOBO Bio-Technology). the linearized pMRNA$^{XP}$ vector and the PCR fragment of spike protein DNA were clean up by Wizard SV Gel and PCR Clean-Up System (Promega). The PCR fragment of spike protein DNA were cloned into linearized pMRNAXP vector using In-Fusion HD Cloning Kit (Clontech Laboratories, Inc.). The cloning mixture was transformed to One Shot™ TOP10 Chemically Competent *E. coli* (Invitrogen™) and incubated at 37 °C overnight. Quick Taq HS DyeMix (TOYOBO Bio-Technology) was used for screening for successful construct. Insert-specific primer and backbone-specific primer were designed for colony PCR. Single clone was selected by pipet tip to conduct PCR. The PCR products were analyzed by agarose gel electrophoresis. the possible candidates were selected and analyzed by DNA sequencing.

**Synthesis of Polymers**

The polymerization process was done according to published procedures[2] with modifications where applicable. Briefly, stock solutions of the monomers (2 M in DMF), initiators (50 mM in DMF, fresh prepared), terminator (iodoacetamide, 0.5 M in $H_2O$, fresh prepared), and triethanolamine (TEOA) buffer (1 M, pH = 7.0) were prepared. The initiator was added to 80 μL of mixture buffer (DMF/TEOA = 1/1) and 10 μL of the monomer stock solution (a v/v ratio of 1:1 was used for hetero-polymers P1/P3, P2/P3, P1/P4, P2/P4, P1/P5, and P2/P5). After 30 min of agitation at room temperature, the polymerization reaction was quenched by addition of 1.9 mL of the terminator stock solution. The resulting polymer was dialyzed against $H_2O$ in the same day. The solution was lyophilized and the polymers were kept at -20 °C. For the in vitro and in vivo experiment, the initiator was mixed with 5% IP followed by the same protocol mentioned above.

**Formulation of WT spike mRNA to form mRNA-PNP**

To obtain the WT spike mRNA, the linear DNA that contained the T7 promoter, 50 untranslated region, 30 untranslated region, S-2P, and poly(A) tail signal sequence was amplified by using TOOLS Ultra High Fidelity DNA Polymerase (BIOTOOLS Co., Ltd.) with 1 μL of the DNA template in an mMESSAGEmMACHINE Kit (Thermo Scientific) at 37 °C for 1 h according to the manufacturer's protocol. The mRNA was purified by RNA cleanup kit (BioLabs) according to the manufacturer's protocol and stored at -80 °C until further use. For the formulation of mRNA-PNP, the mRNA was encapsulated in a corresponding polymer using a self-assembly process, that is, the polymer in ethanol phase (10 mg/mL) was mixed with an aqueous solution of mRNA (1 mg/mL) at pH 4.0 in a 3 to 1 N/P ratio. The mRNA-PNP was dialyzed against PBS buffer (pH 7.4) using Micro Float-A-Lyzer (10 kDa MWCO, spectrum lab) overnight at 4 ºC and stored at -40 °C until further use.

**Quantification of encapsulated mRNA**

Encapsulation efficiency was determined by Quant-iT™ RiboGreen™ RNA Reagent and Kit (Thermo Scientific™). The prepared mRNA polymersome was treated with 10 mM GSH overnight, and then the solutionwas diluted 250-fold with 1X TE buffer and further diluted 2-fold with TE buffer or TE buffer containing 2% Triton X-100. mRNAs were prepared as 100, 50, 25, 12.5 and 0 ng/ml in TE or TE buffer containing 1% Triton X-100 to establish the standard curve. After incubation at 37 ºC for 10 minutes, Quant-iT™ RiboGreen™ RNA Reagent was added into well. The fluorescence intensity was measured by CLARIOstar® Plus (BMG Labtech).

**HEK293T cell transfection**

HEK293T cells were plated at $5*10^5$ cells per well in a 6-well plate in 2.5 mL DMEM media. 1 μg of the GFP mRNA or 3 μg of WT spike mRNA was formulated with the corresponding polymer by the procedure mentioned above and then added to the cells. 18 hours post-transfection, GFP and spike expression were monitored by fluorescence microscopy and western blotting, respectively. For the WB, cells containing spike protein were lysed with 200 μL RIPA lysis buffer including protease inhibitor and incubated for 10 minutes. Cells were then vortexed, centrifuged and analyzed by western blot with polyclonal anti-SARS-CoV-2 S protein antibodies (1:5000 with 1% BSA) followed by HRP conjugated anti-rabbit antibody (1:10000). The spike protein was detected by chemiluminescent HRP substrate and visualized by a trans-illuminator (FUJIFILM LAS3000).

**Cell viability assay**

The HEK293T cells were transfected with mRNA-PNP and the cell viability was evaluated by MTT assay kit. Briefly, HEK293T cells were seeded in 96-well plates overnight. Then, different types of mRNA-PNP were transfected in 100 μl DMEM with 10% FBS. After incubation for 48 h, the media was replaced by 100 μl PBS buffer containing 10 μl of MTT solution in each well for 1 h. The absorption of each well at 450 nm was tested by a microplate reader to calculate the OD values. The cell viability was determined by the formula: $(OD_{experiment} - OD_{blank}) / (OD_{control} - OD_{blank}) \times 100\%$. $OD_{control}$ is the absorbance of the cells without any treatment.

**Glycan-PNP and DC-SIGN binding assay via ELISA**

To assess binding of DC-SIGN to mannoside modified PNP, ELISA plates were coated with mRNA-PNP(I5-P1/P5), mRNA-PNP (I6-P1/P5), mRNA-PNP (I7-P1/P5), mRNA-PNP (I8-P1/P5), mRNA-PNP(I9-P1/P5), or mRNA-PNP(I10-P1/P5) (10 mg/mL) in PBS at 4 °C overnight, respectively. The plates were incubated with diluted DC-SIGN ECD (15 to 0.075 nM in HEPES buffer containing 20 mM HEPES, 150 mM NaCl, 10 mM $CaCl_2$, 0.1% BSA) at pH 7.4, 6.0, and 5.0 for 1 h at rt. The bound DC-SIGN ECD was detected using HRP-conjugated anti-DC-SIGN (B2) IgG antibody (Santa Cruz Biotechnology). After 1 h of incubation at rt, the plates were treated with tetramethlybenzidine (TMB) for 10 min. The optical density was measured at 450 nm after addition of 0.5 M sulfuric acid to the plates using a microplate reader. The apparent Kd was calculated by a nonlinear regression curve fit for total binding using GraphPad Prism.

**Glycan-PNP binding assay to DC-SIGN-Fc, MMR-Fc, MINCLE-Fc, Dectin-2-Fc, and Langerin-Fc via ELISA**

To assess binding of receptor proteins to mannoside modified PNP, ELISA plates were coated with mRNA-PNP(I1-P1/P5), mRNA-PNP (I8-P1/P5), mRNA-PNP (I9-P1/P5), or mRNA-PNP (I10-P1/P5) (10 mg/mL) in PBS at 4 °C overnight, respectively. The plates were incubated with diluted DC-SIGN-Fc, MMR-Fc, MINCLE-Fc, Dectin-2-Fc, and Langerin-Fc (0.625 μg/mL in buffer) at pH 7.4 for 1 h at rt. The bound proteins was detected using HRP-conjugated anti-Fc IgG antibody. After 1 h of incubation at rt, the plates were treated with tetramethlybenzidine (TMB) for 10 min. The mean optical density was measured at 450 nm after addition of 0.5 M sulfuric acid to the plates using a microplate reader.

**Quantification of glycan content on PNP**

A 4M solution of trifluoroaceticacid (2 mL) was added to the PNP solution with stirring at 110 °C for 3 h. The acid was removed, and the resulting residue was lyophilized to dryness. A standard curve was prepared by different concentrations of free mannose (0, 1.5625, 3.125, 6.25, 12.5, 25, and 50 μM). The standards and lyophilized residues were prepared in 30 uL $ddH_2O$, and then loaded to the High-Performance Anion-Exchange Chromatography with Pulsed Amperometric Detection (HPAEC-PAD)(with column: Dionex CarboPac PA10 (2 X 250 mm), flow rate=0.25 mL/min, temp=20 °C, waveform selector: Gold, Garbo, Quad, reference electrode: AgCl, mobile phase: Eluent A= 200 mM NaOH, Eluent B= $H_2O$). The area at corresponding retention time was calculated to determine the amount of mannose.

**Splenic cells preparation and BMDCs culture**

To prepare splenic cells, mouse spleen was homogenized with the frosted end of glass slide, treated with RBC lysis buffer (Sigma) to deplete red blood cells (RBCs), followed by passing through the cell strainer (BD Biosciences). Bone-marrow derived dendritic cells (BMDCs) were prepared as described.[3] Briefly, bone marrow was isolated from mouse femurs and tibiae and treated with RBC lysis buffer (Sigma-Aldrich) to deplete RBCs. Cells were then cultured in RPMI-1640 containing 10% heat inactivated FBS (Thermo Fisher Scientific), 1% Penicillin/Streptomycin (Thermo Fisher Scientific), 50 μM 2-mercaptoethanol (Thermo Fisher Scientific), and 20 ng/ml recombinant mouse GM-CSF (eBioscience) at a density of 2 × 105 cells/ml. The cells were supplemented with an equal volume of the complete culture medium described above at day 3 and refreshed with one-half the volume of medium at day 6. At day 8, the suspended cells were then harvested.

**Treatment of PNP to splenic cells and BMDCs**

Splenic cells or BMDCs were incubated with 1:2000 mRNA-PNP(I1-P1/P4-FITC-P5) or mRNA-PNP(I5-P1/P4-FITC/P5) (diluted by a stock of 10 mg/mL) in RPMI-1640 at 37 °C for 24 hours. Cells were blocked with Fc receptor binding inhibitor (clone: 93, eBioscience) for 20 minutes. Splenocytes were stained with antibodies against CD3 (clone: 17A2, BV421-conjugated, Biolegend), CD19 (clone: 1D3, PECy7-conjugated, BD Biosciences). BMDCs were stained with antibody against CD11c (clone N418 APC-conjugated, Biolegend). Labeled cells were analyzed using FACSC and Flow Cytometer (BD Biosciences).

**C2C12 cell culture**

The mouse muscle myoblast cell line C2C12 was purchase from the Bioresource Collection and Research Center, Taiwan. C2C12 cells were cultured in DMEM with high-glucose (ATCC) supplemented with 10% FBS and 1 × antibiotic-antimycotic. Cells were incubated at 37°C with 5% $CO_2$ and humidified atmosphere control. Culture medium was changed every 2 to 3 days.

**Treatment of polymersomes to C2C12**

Cultured C2C12 myoblasts were detached from a culture dish using 0.25% Trypsin-EDTA (Gibco), and neutralized with growth medium containing 10% FBS. mRNA-PNP(I1-P1/P4-FITC-P5) or mRNA-PNP (I9-P1/P4-FITC/P5) were added to 200 μL C2C12 cells ($2 \times 10^5$ cells) in growth medium to reach a final dilution of 1:1000, 1:2000, 1:4000, or 1:8000 to the original stocks (10 mg/mL) . Three time points were measured: 5 min, 1 hr and 24 hr.

**Flow Cytometry**

After incubation with mRNA-PNP(I1-P1/P4-FITC-P5) or mRNA-PNP(I9-P1/P4-FITC/P5) mRNA, BMDC cells were washed with ice-cold FACS buffer (1% FBS in 1 × DPBS with 0.1% Sodium Azide), and incubated with purified anti-mouse CD16/32 antibody (BioLegend) in FACS buffer on ice for 20 min, followed by washing with FACS buffer. BMDCs were stained with APC anti-mouse CD11c antibody (BioLengend) at 4℃ for 30 min, and washed with FACS buffer. Finally, BMDCs were stained with propidium iodide (Sigma-Aldrich). C2C12 cells were centrifuged and washed with FACS buffer. Cells were stained with propidium iodide. Flow cytometry was performed on FACSCanto flow cytometer (BD Bioscience).

**Animal Immunizations**

BALB/c mice aged 6 to 8 wk old (n = 5) were immunized intramuscularly with 15 μg of mRNA-PNP in phosphate-buffered saline (PBS). Animals were immunized at wk 0 and boosted with a second vaccination at wk 2, and serum samples were collected from each mouse 1 wk after the second immunization.

**Measurement of serum IgG titer**

ELISA was used to determine the IgG titer of the mouse serum. The wells of a 96-well ELISA plate (Greiner Bio-One) were coated with 100 ng SARS-CoV-2 spike protein (ACROBiosystems) in 100mM sodium bicarbonate pH 8.8 at 4°C overnight. The wells were blocked with 200 ul 5% skim milk in 1X PBS at 37 °C for 1 hour and washed with 200 ul PBST (1X PBS, 0.05% Tween 20, pH 7.4) three times. Mice serum samples with 2-fold serial dilution were added into wells for an incubation at 37 °C for 2 hours and washed with 200 ul PBST six times. The wells were incubated with 100 ul HRP conjugated anti-mouse secondary antibody (1:10000, in PBS) at 37 °C for 1 hour and washed with 200 ul PBST six times. 100 μl horseradish peroxidase substrate (1-Step™ Ultra TMB-ELISA Substrate Solution) (Thermo Scientific™) was added into wells followed by 100 ul 1M $H_2SO_4$. After incubation for 30 mins, Absorbance (OD 450 nm) was measured by SpectraMax M5.

**Pseudovirus neutralization assay**

Pseudovirus was constructed by the RNAi Core Facility at Academia Sinica using a procedure similar to that described previously.[4] Briefly, the pseudotyped lentivirus carrying SARS-CoV-2 spike protein was generated by transiently transfecting HEK-293T cells with pCMV-ΔR8.91, pLAS2w.Fluc.Ppuro. HEK-293T cells were seeded one day before transfection, and indicated plasmids were delivery into cells by using TransITR-LT1 transfection reagent (Mirus). The culture medium was refreshed at 16 hr and harvested at 48 hr and 72hr post-transfection. Cell debris was removed by centrifugation at 4,000 xg for 10 min, and the supernatant was passed through 0.45-μm syringe filter (Pall Corporation). The pseudotyped lentivirus was aliquot and then stored at -80°C. To estimate the lentiviral titer by AlarmaBlue assay (Thermo Scientific), The transduction unit (TU) of SARS-CoV-2 pseudotyped lentivirus was estimated by using cell viability assay in responded to the limited dilution of lentivirus. In brief, HEK-293T cells stably expressing human ACE2 gene were plated on 96-well plate one day before lentivirus transduction. For the titering pseudotyped lentivirus, different amounts of lentivirus were added into the culture medium containing polybrene (final concentration 8 μg/ml). Spin infection was carried out at 1,100 xg in 96-well plate for 30 minutes at 37 °C. After incubating cells at 37°C for 16 hr, the culture medium containing virus and polybrene were removed and replaced with fresh complete DMEM containing 2.5 μg/ml puromycin. After treating puromycin for 48 hrs, the culture media was removed and the cell viability was detected by using 10% AlamarBlue reagents according to manufacturer's instruction. The survival rate of uninfected cells (without puromycin treatment) was set as 100%. The virus titer (transduction units) was determined by plotting the survival cells versus diluted viral dose. For neutralization assay, heat-inactivated sera or antibodies were serially diluted and incubated with 1,000 TU of SARS-CoV-2 pseudotyped lentivirus in DMEM for 1 h at 37°C. The mixture was then inoculated with 10,000 HEK-293T cells stably expressing human ACE2 gene in a 96-well plate. The culture medium was replaced with fresh complete DMEM (supplemented with 10% FBS and 100 U/mL penicillin/streptomycin) at 16 h postinfection and continuously cultured for another 48 h. The expression level of luciferase gene was determined by using Bright-Glo Luciferase Assay System (Promega). The relative light unit (RLU) was detected by Tecan i-control (Infinite 500). The percentage of inhibition was calculated as the ratio of RLU reduction in the presence of diluted serum to the RLU value of no serum control using the formula $(RLU^{control} - RLU^{Serum})/RLU$ control.

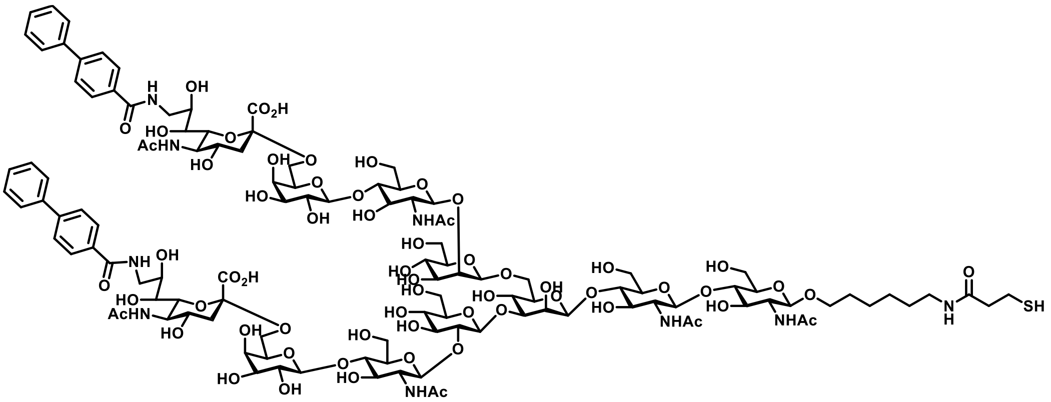

**I2**

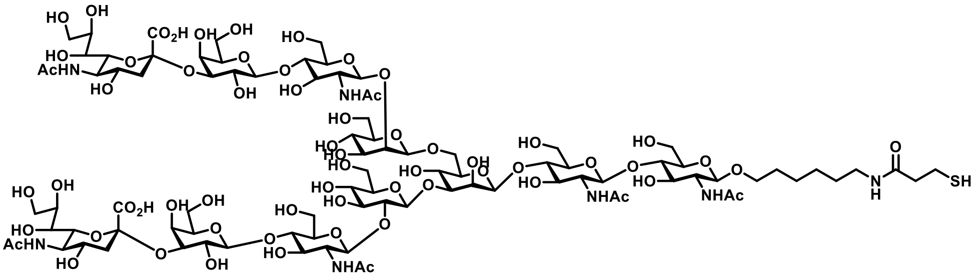

**I3**

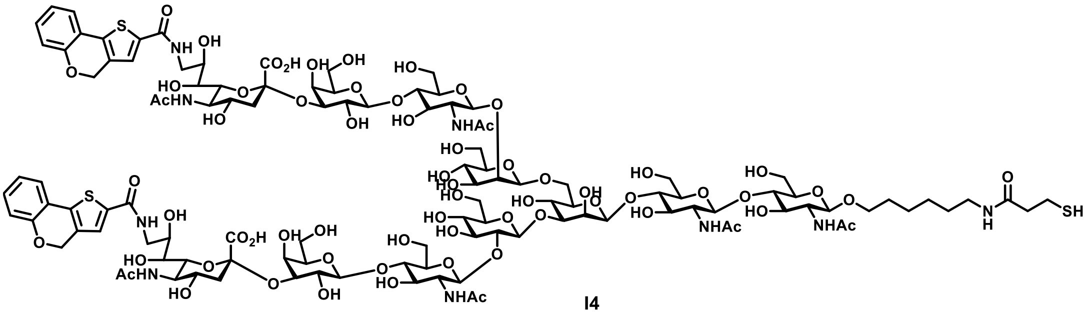

**I4**

**Scheme S1.** Structure of initiator **I2**, **I3**, and **I4**.

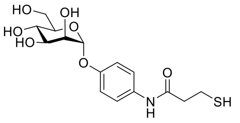

**I5**

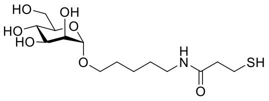

**I6**

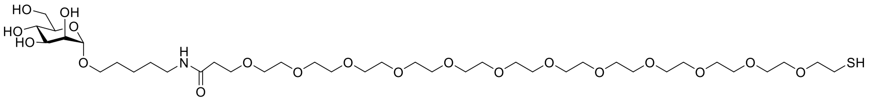

**I7**

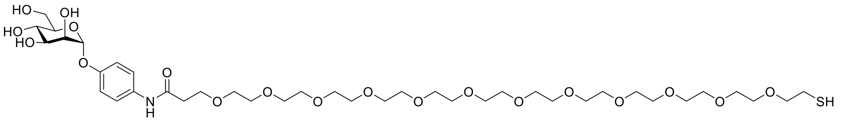

**I8**

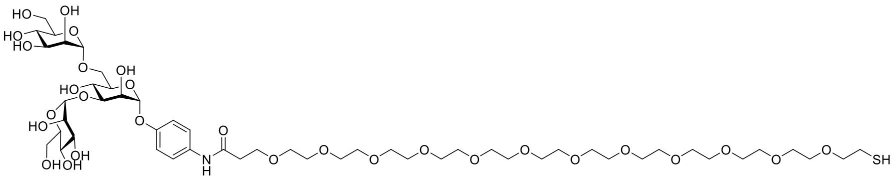

**I9**

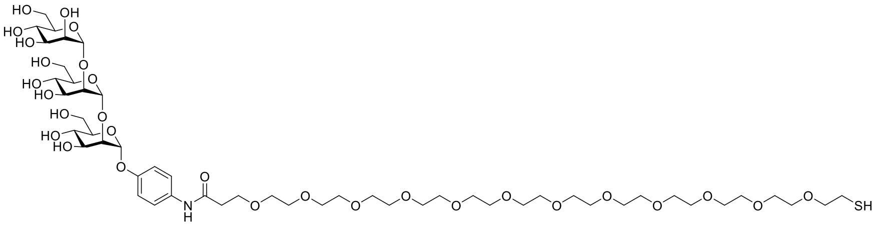

**I10**

**Scheme S2.** Structure of initiator **I5**, **I6**, **I7**, **I8, I9** and **I10**.

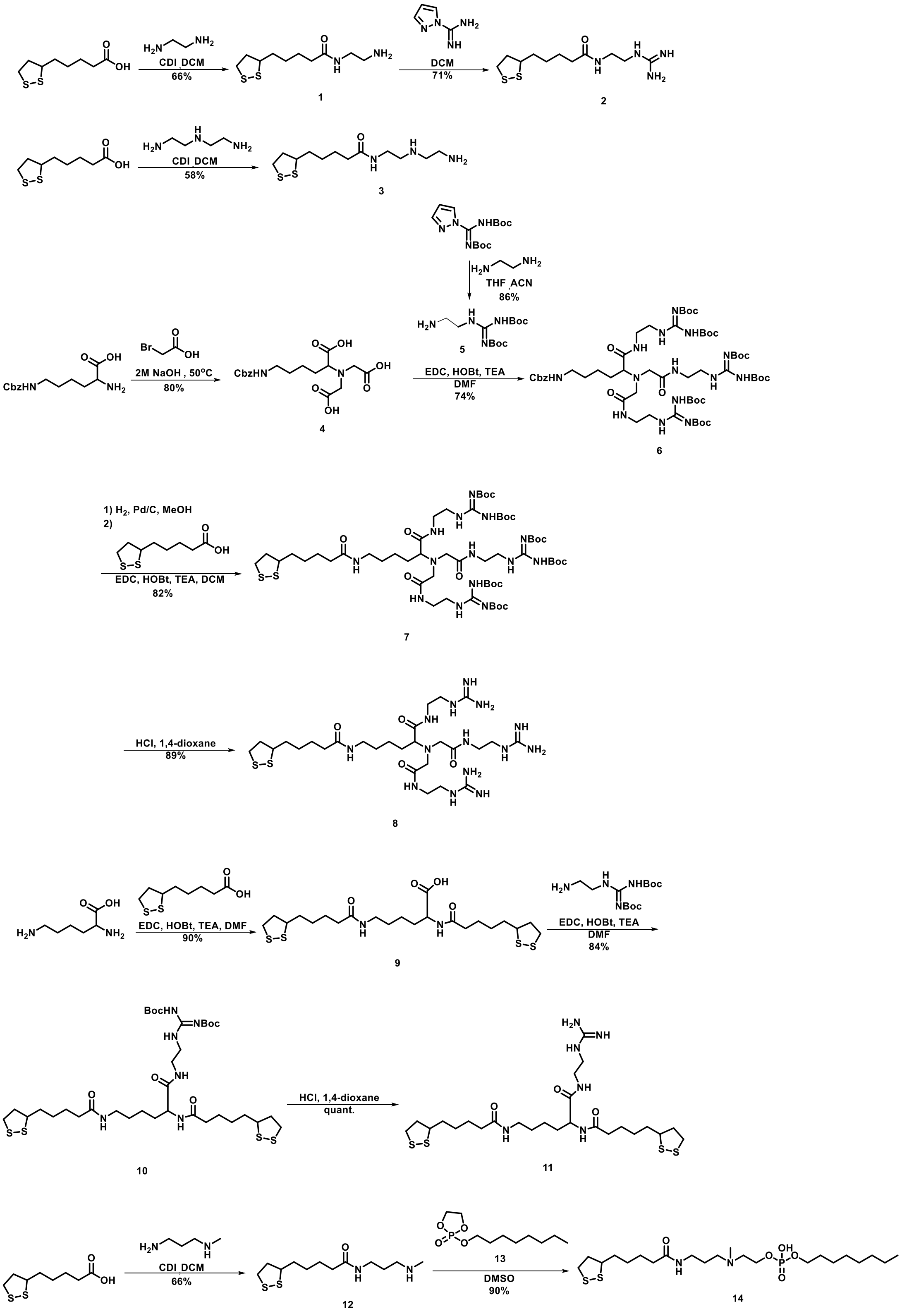


**Scheme S3.** Synthetic procedure of **2**, **3**, **8**, **11, 14**.

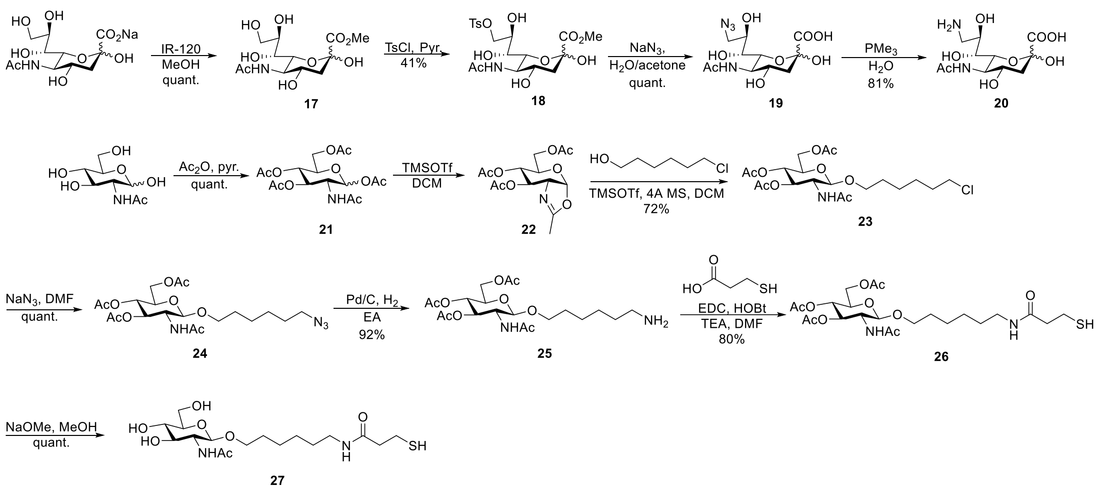

**Scheme S4.** Chemical synthesis of $9^{Am}$Neu5Ac **20** and GlcNAc-SH **27**.

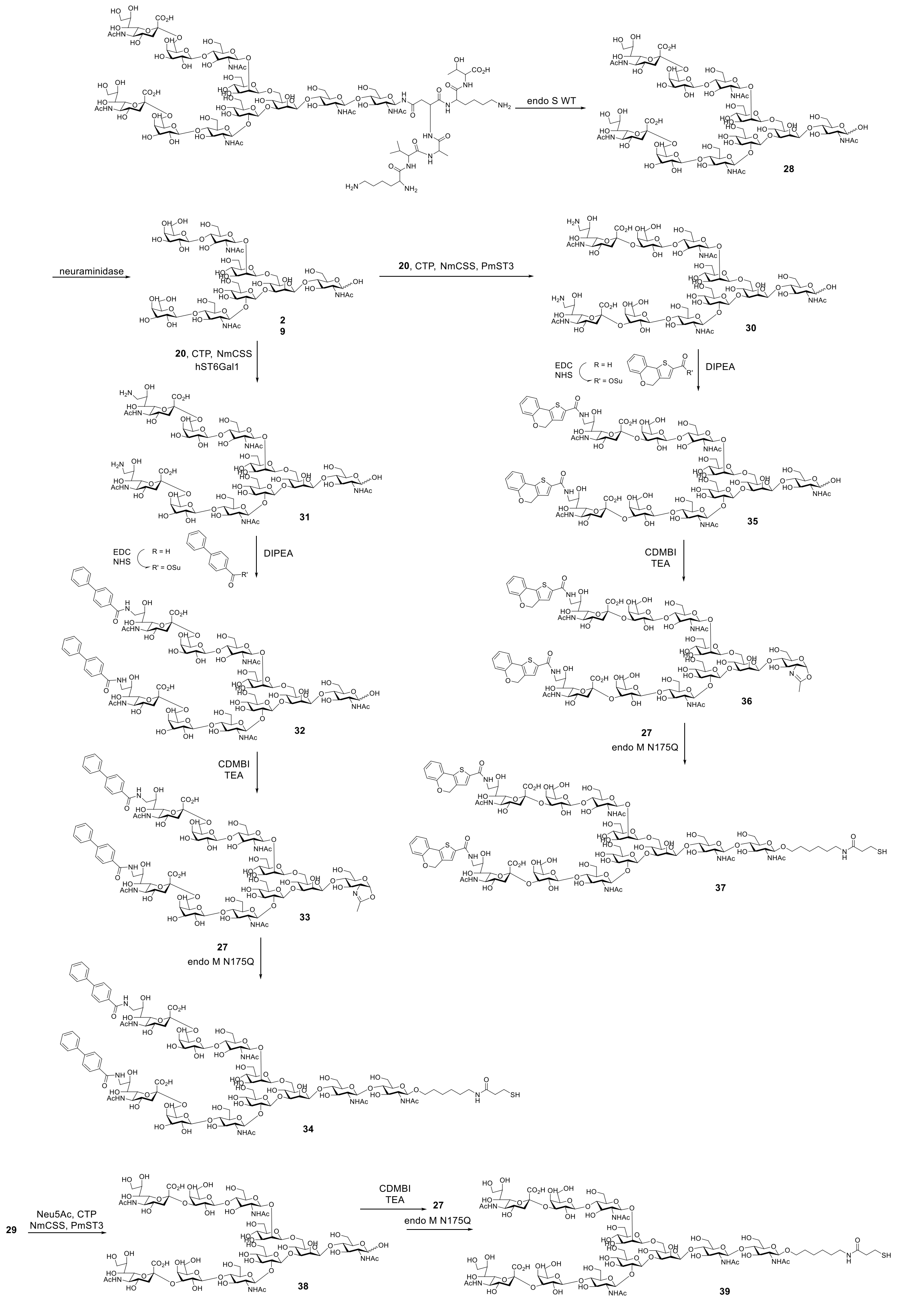

**Scheme S5.** Chemoenzymatic synthesis of $9^{BPC}$Neu5Ac-α2,6-SCT-SH **34**, $9^{TCC}$Neu5Ac-α2,3-SCT-SH **37** and Neu5Ac-α2,3-SCT-SH **39** N-glycan.

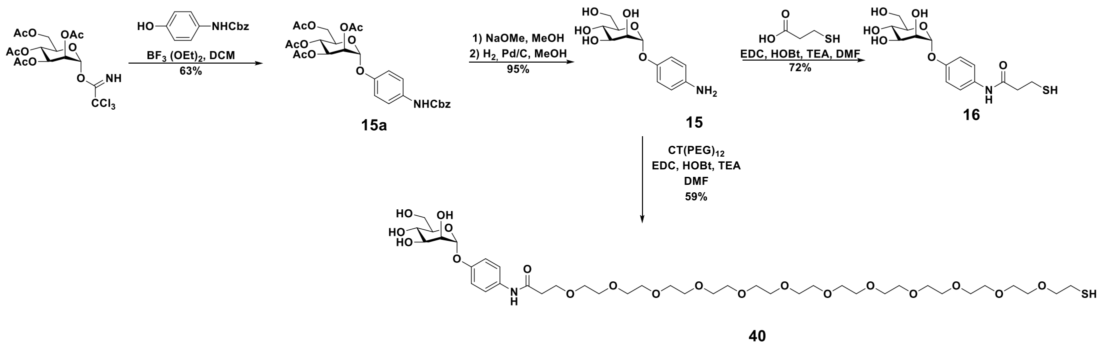
BF3 (OEt)2, DCM
63%
15a
1) NaOMe, MeOH
2) H2, Pd/C, MeOH
95%
15
EDC, HOBt, TEA, DMF
72%
16
CT(PEG)12
EDC, HOBt, TEA
DMF
59%
40

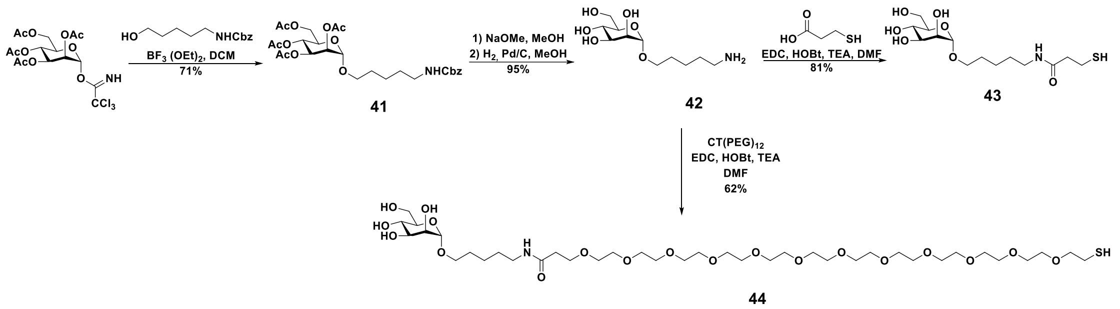
BF3 (OEt)2, DCM
71%
41
1) NaOMe, MeOH
2) H2, Pd/C, MeOH
95%
42
EDC, HOBt, TEA, DMF
81%
43
CT(PEG)12
EDC, HOBt, TEA
DMF
62%
44

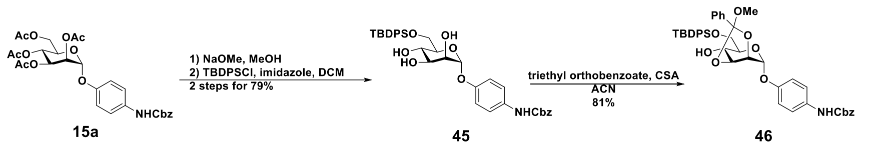
15a
1) NaOMe, MeOH
2) TBDPSCl, imidazole, DCM
2 steps for 79%
45
triethyl orthobenzoate, CSA
ACN
81%
46

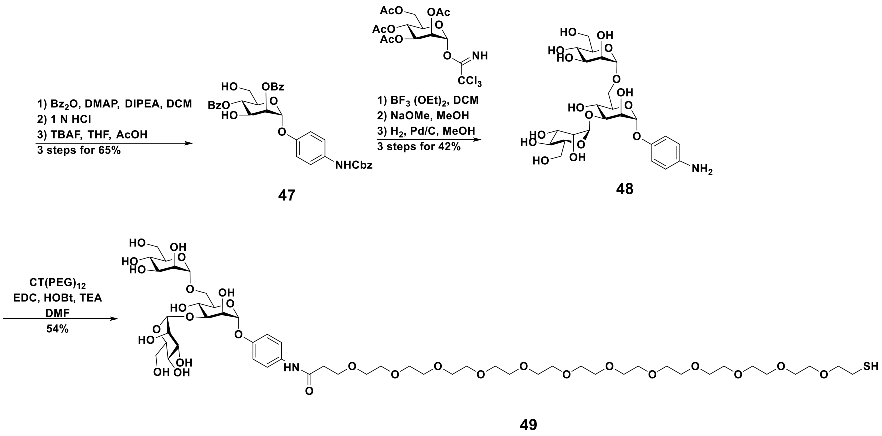
1) Bz2O, DMAP, DIPEA, DCM
2) 1 N HCl
3) TBAF, THF, AcOH
3 steps for 65%
47
1) BF3 (OEt)2, DCM
2) NaOMe, MeOH
3) H2, Pd/C, MeOH
3 steps for 42%
48
CT(PEG)12
EDC, HOBt, TEA
DMF
54%
49

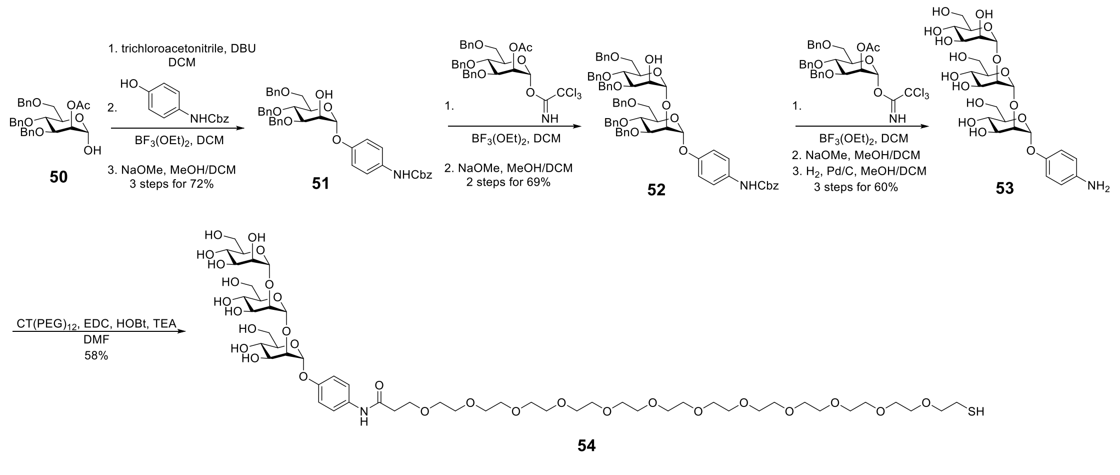
1. trichloroacetonitrile, DBU
DCM
2.
BF3(OEt)2, DCM
3. NaOMe, MeOH/DCM
3 steps for 72%
50
51
1.
BF3(OEt)2, DCM
2. NaOMe, MeOH/DCM
2 steps for 69%
52
1.
BF3(OEt)2, DCM
2. NaOMe, MeOH/DCM
3. H2, Pd/C, MeOH/DCM
3 steps for 60%
53
CT(PEG)12, EDC, HOBt, TEA
DMF
58%
54

**Scheme S6.** Synthetic procedure of **16, 40, 43, 44, 49** and **54**.

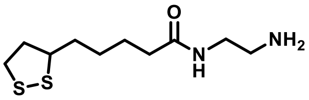

**Compound 1** [2]. Compound **1** was synthesized and characterized based on a published procedure. 1H NMR (600 MHz, $CDCl_3$): δ 5.92 (br, 1H), 3.60-3.57 (m, 1H), 3.29 (dt, *J* = 11.2 Hz, 2H), 3.21-3.10 (m, 2H), 2.86-2.79 (m, 2H), 2.55-2.31 (m, 1H), 2.21 (t, 2H, *J* = 7.4 Hz), 1.90-1.85 (m, 1H), 1.77-1.41 (m, 8H).

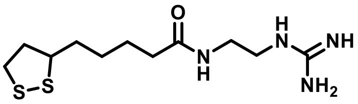

**Compound 2** [2]**.** Compound **2** was synthesized and characterized based on a published procedure. $^1H$ NMR (600 MHz, MeOD): δ 3.98 (s, 1H), 3.61-3.39 (m, 1H), 3.30-3.22 (m, 4H), 3.22-2.88 (m, 2H), 2.56-2.30 (m, 1H), 2.21 (t, *J* = 7.4 Hz, 2H), 1.98-1.72 (m, 1H), 1.79-1.30 (m, 6H).

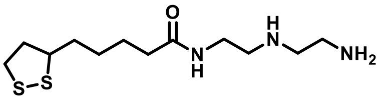

**Compound 3** [5]**.** Compound **3** was synthesized and characterized based on a published procedure. $^1H$ NMR (600 MHz, $CDCl_3$): δ 3.61 – 3.56 (m, 1H), 3.41 – 3.34 (m, 1H), 3.31 – 3.16 (m, 8H), 3.14 – 3.06 (m, 2H), 2.54 – 2.40 (m, 1H), 2.08 (t, *J* = 7.4 Hz, 2H), 1.95 – 1.86 (m, 1H), 1.69 (s, 1H), 1.55 (m, 3H), 1.47 – 1.31 (m, 2H).

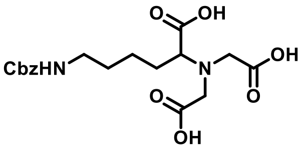

**Compound 4** [2]**.** Compound 4 was synthesized and characterized according to a published protocol. $^1H$ NMR (600 MHz, $DMSO\text{-}d_6$): δ 7.44-7.12 (m, 5H), 5.02 (s, 2H), 3.62-3.41 (m, 4H), 3.35 (t, 1H, *J* = 7.2 Hz), 2.98 (d, 2H, *J* = 5.9 Hz), 1.79-1.03 (m, 6H).

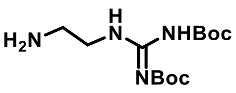

**Compound 5** [6]**.** Compound **5** was synthesized and characterized according to a published protocol. $^1H$ NMR (600 MHz, $CDCl_3$): δ 11.52 (br, 1H), 8.62 (s, 1H), 3.45

(q, $J$ = 4.00 Hz, 2H), 2.86 (t, $J$ = 4.00 Hz, 2H), 1.49 (s, 9H), 1.48 (s, 9H).

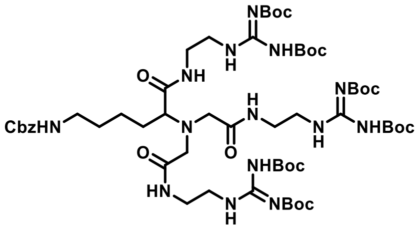


**Compound 6.** A solution of **4** (0.126 mmol) in DMF (1 mL) was preactivated with EDC (0.506 mmol), HOBt (0.506 mmol), and trimethylamine (0.57 mmol) under nitrogen for 30 min. **5** (0.506 mmol) in DMF (1 mL) was then added to the above solution and the resulting solution was stirred at rt for 12 h. The mixture was concentrated to dryness *in vacuo* and then diluted with ethyl acetate. The organic layer was washed with $H_2O$ for three times and was dried over $MgSO_4$, filtered, and concentrated. The crude product was purified by column chromatography on silica gel (MeOH/DCM 1:20) to yield **6** (134 mg, 74%). $^1$H NMR (600 MHz, $CDCl_3$): δ 11.45–11.33 (m, 3H), 8.53–8.45 (m, 3H), 8.35 (t, $J$ = 4.9 Hz, 2H), 7.33–7.25 (m, 5H), 5.04 (s, 2H), 3.59–3.19 (m, 16H), 3.15–3.12 (q, $J$ = 6.0 Hz, 2H), 3.06–3.04 (t, $J$ = 6.9 Hz, 1H), 1.77-1.71 (m, 1H), 1.55-1.32 (m, 59H). HRMS (ESI) calcd for $C_{57}H_{97}N_{14}O_{17}$ $[M+H]^+$ $m/z$ 1249.7156; found: 1249.7166.

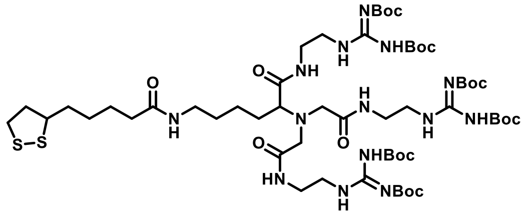


**Compound 7.** A solution of **6** (0.12 mmol) in MeOH (2 mL) was added palladium on charcoal (Pd/C, 10% Pd content, 13 mg). The mixture was stirred at rt under atmosphere of hydrogen gas for 6 h. The solution was filtered through a pad of Celite. The residue was concentrated to dryness in *vacuo* and the resulting residue was dissolved in DCM (2 mL). Lipoic acid (0.152 mmol), EDCI (0.304 mmol), HOBt (0.304 mmol), and TEA (0.304 mmol) was added to the mixture and then stirred at rt for 2 h. The mixture was concentrated to dryness *in vacuo* and then diluted with ethyl acetate. The organic layer was washed with $H_2O$ for three times and was dried over $MgSO_4$, filtered, and concentrated. The crude product was purified by column chromatography on silica gel (MeOH/DCM 1:50) to yield **7** (94 mg, 82%). $^1$H NMR (600 MHz, $CDCl_3$): δ 11.48–11.30 (m, 3H), 8.57–8.46 (m, 3H), 8.40 (t, $J$ = 5.2 Hz, 2H), 8.15-8.12

(m, 1H), 6.22–6.18 (m, 1H), 3.60–3.34 (m, 16H), 3.28–3.04 (m, 7H), 2.48-2.38 (m, 1H), 2.17-2.14 (t, $J$ = 7.4 Hz, 2H), 1.92-1.71 (m, 11H), 1.71-1.59 (m, 4H), 1.57-1.34 (m, 64H). HRMS (ESI) calcd for $C_{57}H_{103}N_{14}O_{16}S_2$ $[M+H]^+$ *m/z* 1303.7118; found: 1303.7129.

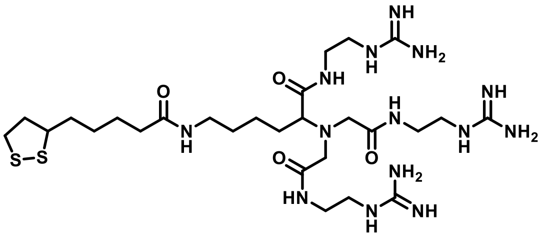

**Compound 8.** **7** (0.08 mmol) was added a solution of 4M HCl (0.5 mL) in 1,4-dioxane (0.5 mL), and the mixture was stirred at rt for 12 h. After that, the solution was removed and dried *in vacuo* to yield **8** (24 mg, 89%). $^1$H NMR (600 MHz, $D_2O$): δ 3.60-3.40 (m, 3H), 3.4-3.16 (m, 16H), 3.02 (s, 2H), 2.76 (s, 2H), 2.20-2.03 (m, 2H), 1.98-1.76 (m, 2H), 1.62-1.16 (m, 11H). $^{13}$C NMR (150 MHz, $D_2O$): δ 176.61, 175.62, 173.68, 172.92, 156.88 (x3), 66.01, 65.73, 56.57, 55.36, 55.24, 40.40 (x2), 40.27, 38.69, 37.99 (x3), 37.82, 35.44, 33.57, 28.66, 28.18, 27.73, 24.99, 22.71. HRMS (ESI) calcd for $C_{27}H_{55}N_{14}O_4S_2$ $[M+H]^+$ *m/z* 703.3967; found: 703.3995.

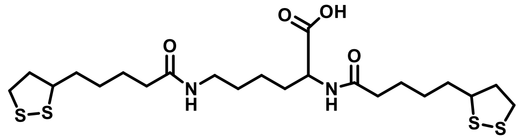

**Compound 9.** A solution of lipoic acid (2.5 mmol) in DMF (3 mL) was preactivated with EDCI (3 mmol), HOBt (3 mmol), and trimethylamine (3 mmol) under nitrogen for 30 min. Lysine (1 mmol) in DMF (3 mL) was then added to the above solution and the resulting solution was stirred at rt for 2 h. The mixture was concentrated to dryness *in vacuo* and then diluted with ethyl acetate. The organic layer was washed with $H_2O$ for three times and was dried over $MgSO_4$, filtered, and concentrated. The crude product was purified by column chromatography on silica gel (MeOH/DCM 1:20) to yield **9** (230 mg, 90%). $^1$H NMR (600 MHz, MeOD): δ 4.37–4.32 (dd, $J$ = 4.6, 9.2 Hz, 1H), 3.63-3.57 (qui, $J$ =6.8 Hz, 2H), 3.22-3.17 (m, 4H), 3.14-3.09 (m, 2H), 2.51-2.45 (m, 2H), 2.29 (t, $J$ = 7.0 Hz, 2H), 2.21 (t, $J$ = 7.0 Hz, 2H), 1.95-1.86 (m, 3H), 1.79-1.62 (m, 9H), 1.60-1.40 (m, 8H). HRMS (ESI) calcd for $C_{57}H_{97}N_{14}O_{17}$ $[M+H]^+$ *m/z* 1249.7156; found: 1249.7166.

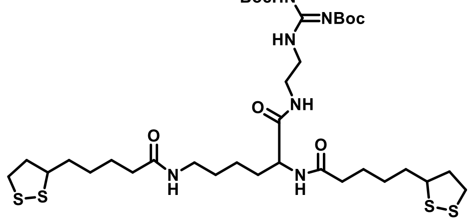

**Compound 10.** A solution of **9** (0.141 mmol) in DMF (1 mL) was preactivated with

EDC (0.211 mmol), HOBt (0.211 mmol), and trimethylamine (0.282 mmol) under nitrogen for 30 min. **5** (0.183 mmol) in DMF (1 mL) was then added to the above solution and the resulting solution was stirred at rt for 2 h. The mixture was concentrated to dryness *in vacuo* and then diluted with ethyl acetate. The organic layer was washed with $H_2O$ for three times and was dried over $MgSO_4$, filtered, and concentrated. The crude product was purified by column chromatography on silica gel (MeOH/DCM 1:20) to yield **10** (92 mg, 84%). $^1$H NMR (600 MHz, $CDCl_3$): δ 11.39 (s, 1H), 8.58 (t, $J$ = 6.0 Hz, 1H), 7.98 (t, $J$ = 4.6 Hz, 1H), 6.41 (d, $J$ = 7.4 Hz, 1H), 5.79-5.74, (m, 1H), 4.37 (q, $J$ = 7.5 Hz, 1H), 3.57-3.48 (m, 4H), 3.44-3.35 (m, 2H), 3.26-3.12 (m, 4H), 3.11-3.05 (m, 2H), 2.46-2.39 (m, 2H), 2.18 (t, $J$ = 7.4 Hz, 2H), 2.13 (t, $J$ = 7.4 Hz, 2H), 1.91-1.84 (m, 2H), 1.82-1.75 (m, 1H), 1.71-1.57 (m, 9H), 1.54-1.35 (m, 20H), 1.33-1.25 (m, 2H). $^{13}$C NMR (150 MHz, $CDCl_3$): δ 172.91, 172.55, 171.72, 162.87, 157.42, 153.03, 83.72, 79.97, 56.46, 56.44, 52.78, 41.19, 40.27, 40.26, 40.20, 38.96, 38.48 (x2), 36.48, 36.37, 36.30, 34.64, 34.61, 32.60, 29.18, 28.95, 28.88, 28.29 (x2), 28.05 (x2), 25.46, 25.36, 25.32, 22.29. HRMS (ESI) calcd for $C_{35}H_{63}N_6O_7S_4$ $[M+H]^+$ *m/z* 807.3641; found: 807.3655.

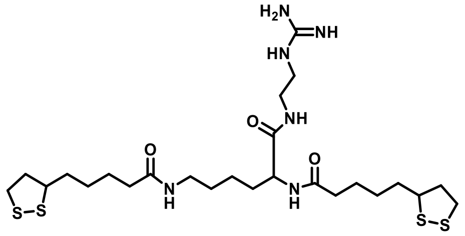

**Compound 11. 10** (0.1 mmol) was added a solution of 4 M HCl (0.5 mL) in 1,4-dioxane (0.5 mL), and the mixture was stirred at rt for 12 h. After that, the solution was removed and dried *in vacuo* to yield **8** (59 mg, quant.). The compound was pure to use directly without further purification.

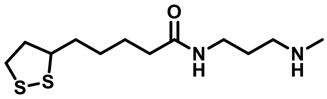

**Compound 12.** A solution of lipoic acid (3 mmol) and CDI (3.9 mmol) were dissolved in 25 ml of anhydrous DCM. This solution was added dropwise at 0 °C to 5 ml of anhydrous DCM containing 8 mmol of N-methyl-1,3-propanediamine. The reaction mixture was stirred 40 min at 0 °C and 30 min at room temperature, then it was washed with $H_2O$ for three times and was dried over $MgSO_4$, filtered, and concentrated to give 12 (589 mg, 66%). $^1$H NMR (600 MHz, $CDCl_3$): δ 3.57-3.50 (m, 1H), 3.31 (q, $J$ = 5.8 Hz, 2H), 3.17-3.12 (m, 1H), 3.11-3.05 (m, 1H), 2.65 (t, $J$ = 6.0 Hz, 2H), 2.46-2.40 (m, 1H), 2.39 (s, 3H), 2.15-2.11 (m, 2H), 1.91-1.84 (m, 1H), 1.72-1.54 (m, 6H), 1.49-1.37 (m, 2H). HRMS (ESI) calcd for $C_{12}H_{26}N_2OS_2$ $[M+H]^+$ *m/z* 277.1408; found: 277.1399.

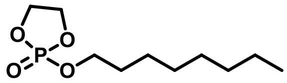

**Compound 13** [7]**.** Compound **13** was synthesized and characterized according to a published protocol. $^1$H NMR (600 MHz, $CDCl_3$): δ 4.50-4.38 (m, 4H), 4.19-4.15 (m, 2H), 1.74-1.70 (m, 2H), 1.38-1.20 (m, 12H), 0.89 (t, *J* = 7.2 Hz, 3H)

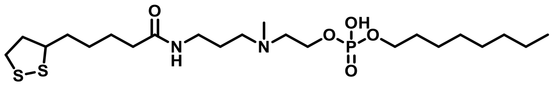

**Compound 14.** A solution of **12** (0.25 mmol) in 1 ml anhydrous DMF was added **13** (0.25 mmol), and the reaction was stirred at 70 °C for 24 h. The mixture was concentrated to dryness *in vacuo* to give **14** (116 mg, 90%). $^1$H NMR (600 MHz, MeOD): δ 7.90 (s, 1H), 4.14-4.01 (m, 1H), 3.91-3.83 (m, 1H), 3.73-3.67 (m, 1H), 3.62-3.55 (m, 1H), 3.28 (t, *J* = 6.6 Hz, 2H), 3.21-3.15 (m, 1H), 3.13-3.06 (m, 1H), 2.98 (t, *J* = 7.2 Hz, 2H), 2.69 (s, 3H), 2.50-2.41 (m, 1H), 2.24 (t, *J* = 7.2 Hz, 2H), 1.93-1.84 (m, 1H), 1.76-1.59 (m, 6H), 1.51-1.43 (m, 2H), 1.43-1.37 (m ,1H), 1.36-1.25 (m, 5H), 0.92-0.88 (m, 3H). $^{13}$C NMR (150 MHz, MeOD): δ 177.08, 79.68, 57.79, 48.91, 48.77, 48.04, 41.54, 39.55, 37.06, 36.91, 35.91, 33.85, 33.20, 32.09, 32.05, 30.44, 30.12, 27.73, 27.16, 26.81, 23.92, 14.65.HRMS (ESI) calcd for $C_{22}H_{44}N_2O_5PS_2$ [M-H]$^-$ *m/z* 511.2429; found: 511.2423.

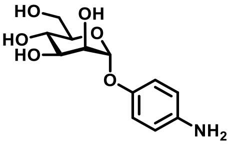

**Compound 15**[8]. Compound 15 was synthesized and characterized according to a published protocol. $^1$H NMR (600 MHz, MeOD): δ 6.93 (d, *J* = 8.6 Hz, 2H), 6.71 (d, *J* = 8.6 Hz, 2H), 5.3 (s, 1H), 4.00-3.99 (m, 1H), 3.89-3.88 (m, 1H), 3.81-3.79 (m, 1H), 3.79-3.70 (m, 2H), 3.69-3.67 (m, 1H).

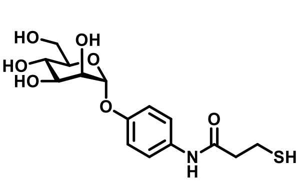

**Compound 16.** A solution of **15** (0.24 mmol) in DMF (2 mL) was added EDC (0.24 mmol), HOBt (0.24 mmol), trimethylamine (0.4 mmol), and 3-mercaptopropionic acid (0.2 mmol), and the resulting solution was stirred under nitrogen at rt for 2 h. The mixture was concentrated to dryness *in vacuo*, and the crude product was purified by column chromatography on silica gel (MeOH/DCM 1:2) to yield **16** (72%). $^1$H NMR (600 MHz, MeOD) δ 6.94 (d, *J* = 8.6 Hz, 2H), 6.72 (d, *J* = 8.6 Hz, 2H), 5.30 (s, 1H),

4.00 (dd, $J$ = 3.4, 1.8 Hz, 1H), 3.89 (dd, $J$ = 9.7, 3.4 Hz, 1H), 3.81-3.67 (m, 4H), 2.72 (t, $J$ = 6.8 Hz, 2H), 2.60 (t, $J$ = 6.8 Hz, 2H). $^{13}$C NMR (150 MHz, MeOD) δ 176.55, 151.13, 143.23, 119.25(x2), 117.93 (x2), 101.49, 101.32, 75.10, 72.43, 72.18, 68.42, 62.66, 40.34, 20.70. HRMS (ESI) calcd for $C_{15}H_{22}NO_7S$ $[M+H]^+$: 360.1117, found 360.1101.

**Compound 17-20** [9] were synthesized according to published article.
**Compound 21-25** [10] were synthesized according to published article.

**Compound 26.**
Compound **25**[11] (0.43 mmol) in DCM (5 mL) was added EDC (0.52 mmol), HOBt (0.52 mmol), trimethylamine (0.86 mmol), and 3-mercaptopropionic acid (0.47 mmol), and the resulting solution was stirred under nitrogen at rt for 2 h. The mixture was concentrated to dryness *in vacuo*, and the crude product was purified by column chromatography on silica gel (MeOH/DCM 1:10) to yield **26** (184 mg, 80%). $^1$H NMR (600 MHz, $CDCl_3$) δ 5.96 (d, $J$ = 8.6 Hz, 1H), 5.28 (t, $J$ = 9.6 Hz, 1H), 5.05 (t, $J$ = 9.4 Hz, 1H), 4.64 (d, $J$ = 8.4 Hz, 1H), 4.24 (dd, $J$ = 12.3, 4.7 Hz, 1H), 4.11 (dd, $J$ = 12.3, 2.5 Hz, 1H), 3.86-3.80 (m, 2H), 3.68-3.66 (m, 1H), 3.47-3.43 (m, 1H), 3.34-3.29 (m, 1H), 3.22-3.16 (m, 1H), 2.81 (t, $J$ = 8.2 Hz, 2H), 2.51-2.45 (m, 2H), 2.06 (s, 3 H), 2.01 (s, 3 H), 2.00 (s, 3 H), 1.93 (s, 3 H), 1.57-1.51 (m, 4H), 1.48 (t, $J$ = 7.0 Hz, 2H) 1.37-1.29 (m, 4H). HRMS (ESI) calcd for $C_{23}H_{38}N_2O_{10}S$ $[M+H]^+$: 535.2325, found 535.2314.

**Compound 27.**
Compound **26** in MeOH was added NaOMe and the resulting solution was stirred under nitrogen at rt for 2 h. The mixture was neutralized by IR-120, and then filtered, concentrated to dryness in vacuo to yield **27** (92 mg, quant.) $^1$H NMR (600 MHz, MeOD) δ 4.38 (d, $J$ = 9.2 Hz, 1H), 3.90-3.86 (m, 2H), 3.68 (dd, $J$ = 12.9, 5.9 Hz, 1H), 3.65-3.62 (m, 1H), 3.48-3.43 (m, 2H), 3.28-3.24 (m, 1H), 3.20-3.14 (m, 2H), 2.94 (t, $J$ = 7.3 Hz, 1H), 2.73 (t, $J$ = 7.3 Hz, 1H), 2.59 (t, $J$ = 7.3 Hz, 1H), 2.47 (t, $J$ = 7.3 Hz, 1H), 1.97 (s, 3 H), 1.56-1.48 (m, 4H), 1.39-1.35 (m, 4H). $^{13}$C NMR (150 MHz, MeOD): δ 173.70, 169.75, 102.74, 77.92, 76.07, 70.50, 62.76, 57.35, 41.02, 40.32, 36.45, 35.19, 30.49, 27.66, 26.73, 23.03. HRMS (ESI) calcd for $C_{17}H_{32}N_2O_{17}S$ $[M+H]^+$: 409.2008, found 409.2017.

**Preparation of $9^{Am}$Neu5Ac-α2,3-SCT 30, $9^{Am}$Neu5Ac-α2,6-SCT 31 and Neu5Ac-α2,3-SCT 38.**
30 mg sialylglycopeptide (SGP) was digested by Endo-S WT (300 μg) in Tris-HCl buffer at 37°C for 48 h, and purified by Sephadex G-25 gel filtration chromatography,

and the product was analyzed by ESI-MS to give SCT compound **28**.[12] Neuraminidase (5 U/ml, 12 μL) in Tris-HCl buffer was added to the mixture at 37°C for 12 h, and purified by Sephadex G-25 gel filtration chromatography to give desialylated N-glycan **29**. After that, the reaction was carried out in 0.5 mL of HEPES buffer (50 mM, pH 8.5) containing 100 mM of $9^{Am}$Neu5Ac (14.5 mg, 47 μmol), 110 mM of CTP (27.2 mg, 52 μmol), and 20 mM of $MgCl_2$. The pH of reaction mixture was adjusted to 8.5 by adding 2N NaOH. Then, 0.5 mg/mL of NmCSS was added to the above solution. The resulting mixture was incubated at 37 °C for 8 h and the formation of CMP-$9^{Am}$Neu5Ac was monitored by TLC analysis. For $9^{Am}$Neu5Ac-α2,6-SCT, hST6Gal-I (0.5 mg/mL) were added to the $9^{Am}$Neu5Ac reaction mixture with N-glycan and incubated at 37 °C.[13] For $9^{Am}$Neu5Ac-α2,3-SCT, PmST3 (0.3 mg/mL) were added to the $9^{Am}$Neu5Ac reaction mixture with N-glycan and incubated at 37 °C. For Neu5Ac-α2,3-SCT, PmST3 (0.3 mg/mL) were added to the Neu5Ac reaction mixture with N-glycan and incubated at 37 °C .The reactions above were monitored by mass spectrometry analysis and TLC. After the acceptor was consumed, the reaction was centrifuged, and the supernatant was subjected to a centrifuge filter with molecular mass cutoff 10 kDa (Amicon Ultra, Millipore) to remove proteins. The filtrate was purified with P-2 gel filtration chromatography to give $9^{Am}$Neu5Ac-α2,3-SCT **30**, $9^{Am}$Neu5Ac-α2,6-SCT **31** and Neu5Ac-α2,3-SCT **38**.

**Preparation of $9^{BPC}$Neu5Ac-α2,6-SCT-SH 34, $9^{TCC}$Neu5Ac-α2,3-SCT-SH 37 and Neu5Ac-α2,3-SCT-SH 39.**

$9^{Am}$Neu5Ac-α2,6-SCT and DIEA (5.0 eq) were dissolved in $H_2O$, followed with addition of biphenyl carboxylic acid-N-hydrozysuccinimide ester (BPC-NHS) (3 eq) in THF. The reaction was stirred at 0 °C until the starting material was consumed. Then the reaction was purified by Sep-Pak C18 column (2 g, Waters Corp.) and eluted with $H_2O$-MeOH to give compound **32** in 91% yield. $9^{Am}$Neu5Ac-α2,3-SCT was prepared similarly as described above by stirring with 4H-thieno[3,2-c]chromene-2-carbamoyl-NHS (TCC-NHS) in THF and $H_2O$ to give compound **35**. After purification, the product was afforded in 91% yield.

The mixture of $9^{BPC}$Neu5Ac-α2,6-SCT in water was added CDMBI and TEA and incubated at 4°C for 1 h[12] to generate corresponding oxazoline N-glycan **33**, which was purified by Sephadex G-25 gel filtration chromatography and characterized by ESI-MS. GlcNAc-SH **27** (0.25 mg) and Endo-M (N175Q) (1.6 U/mL) were added to a solution of 50 mM phosphate buffer (pH 7) with $9^{BPC}$Neu5Ac-α2,6-SCT-oxazoline and incubated at 30 °C for 30 min. The transglycosylation products was isolated by P-2 gel filtration chromatography to give compound **34** and characterized by ESI-MS. $9^{TCC}$Neu5Ac-α2,3-SCT-SH **37** and Neu5Ac-α2,3-SCT-SH **39** was prepared same as described above.

**$9^{BPC}$Neu5Ac-α 2,6-SCT-SH 34.**

$^1$H NMR (600 MHz, DMSO-d$_6$): δ = 8.33 (s, 3H, NH), 8.00 (d, *J* = 8.6 Hz, 4H), 7.72 (dd, *J* = 8.0, 14.2 Hz, 9H), 7.49 (t, *J* = 8.3 Hz, 4H), 7.40 (t, *J* = 8.3 Hz, 2H) , 4.98-4.95 (m, 2H), 4.76 (s, 1H), 4.54 (s, 1H), 4.44-4.39 (m, 2H), 4.25-4.22 (m, 3H), 3.99 (s, 1H), 3.87 (s, 1H), 3.83-3.73 (m, 5H), 3.41-3.25 (m, 58H), 3.21 – 3.16 (m, 3H), 3.11 – 2.99 (m, 4H), 2.87 (t, *J* = 6.9 Hz, 2H), 2.61 (d, *J* = 8.4 Hz, 2H), 2.44 (t, *J* = 7.4 Hz, 2H), 1.91-1.77 (m, 18H), 1.44 (t, *J* = 6.9 Hz, 2H), 1.39-1.28 (m, 4H), 1.28-1.19 (m, 4H), 1.12 (t, *J* = 6.9 Hz, 1H). HRMS (ESI) calcd for $C_{119}H_{171}N_9O_{63}S^{2-}$ $[M-2H]^{2-}$: 1383.0093, found 1383.0077.

**$9^{TCC}$Neu5Ac-α2,3-SCT-SH 37.**

$^1$H NMR (600 MHz, DMSO-d$_6$): δ = 8.22 (s, 3H, NH), 8.10-8.09 (m, 1H, NH), 7.96-7.93 (m, 1H, NH), 7.81-7.79 (m, 1H, NH), 7.74-7.72 (m, 1H, NH), 7.35 (d, *J* = 7.8 Hz, 2H), 7.29 (s, 2H), 7.20 (t, *J* = 7.8 Hz, 2H), 6.97 (t, *J* = 7.8 Hz, 2H), 6.92 (d, *J* = 7.9 Hz, 2H), 5.29 (s, 4H), 5.01-4.95 (m, 2H), 4.77 (s, 1H), 4.54 (s, 1H), 4.44-4.36 (m, 2H), 4.27-4.19 (m, 3H), 3.99 (s, 1H), 3.88 (s, 1H), 3.85-3.71 (m, 3H), 3.50-3.22 (m, 52H), 3.19 (d, *J* = 9.5 Hz, 2H), 3.10-3.00 (m, 4H), 2.87 (t, *J* = 6.9 Hz, 2H), 2.64 – 2.58 (m, 2H), 2.44 (t, *J* = 7.4 Hz, 2H), 1.93-1.78 (m, 18H), 1.45-1.39 (m, 2H), 1.39-1.29 (m, 4H), 1.28-1.20 (m, 4H), 1.16 (t, *J* = 6.9 Hz, 1H). HRMS (ESI) calcd for $C_{117}H_{169}N_9O_{65}S_3^{2-}$ $[M-2H]^{2-}$: 1416.9606, found 1416.9623.

**Neu5Ac-α2,3-SCT-SH 39.**

$^1$H NMR (600 MHz, DMSO-d$_6$): δ = 8.33 (s, 3H, NH), 8.10 (s, 1H, NH), 7.96-7.94 (m, 1H, NH), 7.81-7.80 (m, 1H, NH), 7.74-7.73 (m, 1H, NH), 4.98-4.95 (m, 2H), 4.76-4.75 (m, 1H), 4.54 (s, 1H), 4.46-4.38 (m, 2H), 4.25-4.22 (m, 3H), 3.99 (s, 1H), 3.87 (s, 1H), 3.80-3.71 (m, 4H), 3.41-3.25 (m, 59H), 3.23 – 3.18 (m, 3H), 3.08 – 3.01 (m, 3H), 2.87 (t, *J* = 6.9 Hz, 2H), 2.63-2.60 (m, 2H), 2.44 (t, *J* = 7.4 Hz, 2H), 1.87-1.79 (m, 18H), 1.44 (t, *J* = 6.9 Hz, 2H), 1.39-1.28 (m, 4H), 1.28-1.19 (m, 4H), 1.07 (t, *J* = 6.9 Hz, 1H). HRMS (ESI) calcd for $C_{93}H_{153}N_7O_{63}S^{2-}$ $[M-2H]^{2-}$: 1203.9357, found 1203.9377

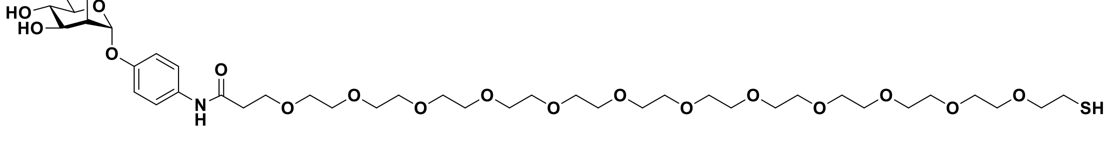

**Compound 40.**

Compound **15** (0.12 mmol) in DMF (1 mL) was added EDC (0.12 mmol), HOBt (0.12

mmol), DMAP(0.12 mmol), trimethylamine (0.2 mmol), and CT(PEG)$_{12}$ (0.1 mmol), and the resulting solution was stirred under nitrogen at rt for 12 h. The mixture was concentrated to dryness *in vacuo*, and the crude product was purified by column chromatography on silica gel to yield **40** (59%). 1H NMR (600 MHz, MeOD): δ 6.93 (d, *J* = 8.6 Hz, 2H), 6.71 (d, *J* = 8.6 Hz, 2H), 5.3 (s, 1H), 4.00-3.99 (m, 1H), 3.89-3.88 (m, 1H), 3.81-3.70 (m, 5H), 3.70-3.59 (m, 48H), 2.68 (t, *J* = 6.8 Hz, 2H), 2.50-2.47 (m, 2H). $^{13}$C NMR (150 MHz, MeOD): δ 170.17, 151.03, 143.50, 119.26(x2), 117.82(x2), 101.36, 75.10, 74.08, 72.45, 72.19, 71.45, 71.41, 71.37, 71.30, 71.24, 71.11, 70.93(x18), 68.43, 62.68, 24.67. HRMS (ESI) calcd for $C_{39}H_{70}NO_{19}S$ [M+H]$^+$: 888.4263, found 888.4241.

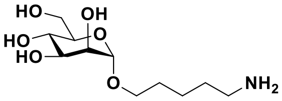


**Compound 40, 41**[14] were synthesized according to our published article.

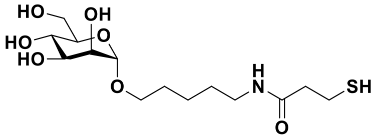


**Compound 43.** A solution of **42** (0.24 mmol) in DMF (2 mL) was added EDC (0.24 mmol), HOBt (0.24 mmol), trimethylamine (0.4 mmol), and 3-mercaptopropionic acid (0.2 mmol), and the resulting solution was stirred under nitrogen at rt for 2 h. The mixture was concentrated to dryness *in vacuo*, and the crude product was purified by column chromatography on silica gel (MeOH/DCM 1:3) to yield **16** (81%). $^1$H NMR (600 MHz, MeOD) δ 4.91 (s, 1H), 3.84 (dd, *J* = 3.4, 1.8 Hz, 1H), 3.82-3.76 (m, 2H), 3.75-3.67 (m, 2H), 3.62 (m, 1H), 3.56-3.41 (m, 2H), 2.94 (t, *J* = 7.0 Hz, 2H), 2.70 (t, *J* = 7.2 Hz, 2H), 2.48 (t, *J* = 7.2 Hz, 2H). HRMS (ESI) calcd for $C_{14}H_{28}NO_7S$ [M+H]$^+$: 354.1586, found 354.1602.

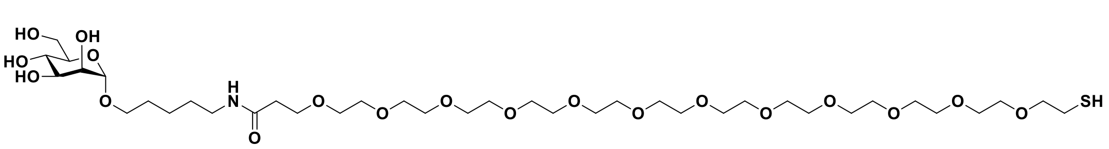


**Compound 44.** Compound **42** (0.12 mmol) in DMF (1 mL) was added EDC (0.12 mmol), HOBt (0.12 mmol), trimethylamine (0.2 mmol), and CT(PEG)$_{12}$ (0.1 mmol), and the resulting solution was stirred under nitrogen at rt for 12 h. The mixture was concentrated to dryness *in vacuo*, and the crude product was purified by column chromatography on silica gel to yield **44** (62%). $^1$H NMR (600 MHz, $D_2O$): δ 4.77 (d,

$J$ = 8.6 Hz, 2H), 3.85-3.82 (m, 1H), 3.81-3.77 (m, 1H), 3.69-3.67 (m, 1H), 3.66-3.64 (m, 2H), 3.63-3.57 (m, 41H), 3.55-3.45 (m, 4H), 2.88 (t, $J$ = 6.8 Hz, 2H), 2.57 (t, $J$ = 7.2 Hz, 2H), 2.37 (t, $J$ = 7.2 Hz, 2H), 1.66-1.51 (m, 4H), 1.40-1.31 (m, 2H). $^{13}$C NMR (150 MHz, $D_2O$): δ 171.01, 99.62, 72.72, 70.57, 69.99, 69.24 (x22), 69.04, 67.90, 67.39, 66.73, 60.92, 39.36, 37.57, 27.96, 26.70, 22.91, 22.43. HRMS (ESI) calcd for $C_{38}H_{76}NO_{19}S$ $[M+H]^+$: 882.4732, found 882.4750.

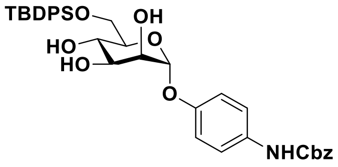


**Compound 45.** Compound **15b** (5 mmol) in MeOH was added NaOMe (0.2 eq) and the resulting solution was stirred under nitrogen at rt for 2 h. The mixture was neutralized by IR-120, and then filtered, concentrated to dryness in vacuo. It was then dissolved in anhydrous DCM (40 mL) and treated with imidazole (7.5 mmol) at 0 °C, followed by addition of TBDPSCl (5.5 mmol). The mixture was stirred at room temperature for 2.5 h under nitrogen atmosphere. The reaction was quenched by addition of MeOH. After stirring at room temperature for 10 min, the solvent was removed under reduced pressure to give a dry residue that was purified by column chromatography with MeOH/DCM (1/10) to give compound **45** (75%).$^1$H NMR (600 MHz, $CDCl_3$) δ 7.61 (m, 4H), 7.41-7.29 (m, 11H), 7.23-7.18 (m, 2H), 6.92 (d, $J$ = 9.3 Hz, 2H), 5.41 (s, 1H), 5.16 (s, 2H), 4.09 (s, 1H), 4.02-4.00 (dd, $J$ = 9.3, 3.4 Hz, 1H), 3.91 (t, $J$ = 9.3 Hz, 1H), 3.85 (d, $J$ = 5.1 Hz, 2H), 3.72-3.69 (m, 1H), 1.01, (s, 9H). $^{13}$C NMR (150 MHz, $CDCl_3$): 162.61, 135.64(x4), 135.54(x4), 132.73, 129.95(x4), 128.64(x4), 128.38, 128.33, 127.84(x2), 127.81(x2), 98.09, 71.41, 71.22, 70.22, 70.13, 64.89, 36.53, 31.48, 26.83(x3), 19.19. HRMS (ESI) calcd for $C_{36}H_{42}NO_8Si$ $[M+H]^+$: 644.2680, found 644.2699.

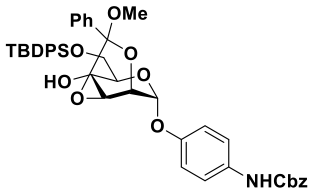


**Compound 46.** To a solution of compound **45** (3 mmol) and a catalytic amount of CSA (0.3 mmol) in $CH_3CN$ (60 mL) was added trimethyl orthobenzoate (9 mmol) at room temperature under atmospheric pressure of nitrogen. After stirring for 30 min, $Et_3N$ was added to quench the reaction and the resulting mixture was dried under reduced pressure.

The residue was purified by column chromatography with EA/Hex (1/2) to give compound **46** (81%).

$^{1}$H NMR (600 MHz, $CDCl_3$) δ 7.65-7.59 (m, 2H), 7.57-7.52 (m, 4H), 7.41-7.27 (m, 14H), 7.26-7.20 (m, 2H), 6.93 (d, *J* = 9.2 Hz, 2H), 5.77 (s, 1H), 5.17 (s, 2H), 4.70 (d, *J* = 6.1 Hz,1H), 4.58 (dd, *J* = 9.3, 3.4 Hz, 1H), 3.79-3.76 (m, 2H), 3.74-3.70 (m, 1H), 3.69-3.66 (m, 1H), 3.22 (s, 3H), 2.53 (d, *J* = 3.9 Hz, 1H), 0.93, (s, 9H). $^{13}$C NMR (150 MHz, $CDCl_3$): 171.23, 153.49, 152.24, 137.09, 136.09, 135.68(x4), 135.48(x4), 132.98, 132.72, 129.86, 129.84(x2), 129.18, 128.65(x2), 128.39, 128.37, 128.34, 127.78(x2), 127.71(x2), 126.23, 121.11, 117.18, 95.69, 79.52, 69.57, 69.45, 67.03, 63.75, 60.44, 51.16, 26.76(x3), 19.15, 14.22. HRMS (ESI) calcd for $C_{44}H_{48}NO_9Si$ $[M+H]^+$:762.3098, found 762.3072.

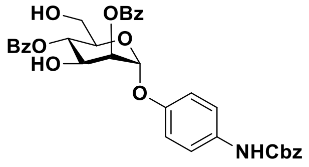


**Compound 47.** Compound **46** (2 mmol) was dissolved in DCM (20 mL) and sequentially mixed with DIPEA (6 mmol), benzoic anhydride (4 mmol) and DMAP (0.2 mmol). After stirring for 30 min, the solvent was evaporated under reduced pressure to give a dry residue and then poured into EA (20 mL) and 2 N HCl (20 mL) with vigorous stirring for 30 min. The solvent was removed by evaporation, followed by extraction with EA. The collected organic layer was washed with ice-cold saturated $NaHCO_3$(aq), water and brine, and dried over $MgSO_4$. The filtrate was evaporated under reduced pressure and redissolved in THF (20 mL). It was added AcOH (4 mmol) and 1 M TBAF (2.4 mmol in THF) at 0 °C. The resulting mixture was warmed up to room temperature gradually, stirred for another 2 h, and then diluted with EA. The organic layer was washed with saturated $NaHCO_3$(aq), water and brine, dried with anhydrous $MgSO_4$, and concentrated under reduced pressure. The dry residue was purified by column chromatography with EA/Hex (1/2) to give compound **47** (65%). $^{1}$H NMR (600 MHz, $CDCl_3$) δ 8.10-8.05 (m, 4H), 7.61-7.56 (m, 2H), 7.48-7.41 (m, 4H), 7.39-7.27 (m, 7H), 7.03 (d, *J* = 9.2 Hz, 2H), 5.69 (s, 1H), 5.62-5.55 (m, 2H), 5.16, (s, 2H), 4.62 (dd, *J* = 9.3, 3.4 Hz, 1H), 4.06-4.01 (m, 1H), 3.76-3.67 (m, 2H).$^{13}$C NMR (150 MHz, $CDCl_3$): 167.42, 166.05, 153.55, 152.01, 136.03, 133.84, 133.76, 130.00 (x4), 129.09, 128.96, 128.69(x4), 128.65(x4), 128.62(x4), 128.34, 117.11 (x2), 96.18, 72.61, 71.30, 70.01, 68.53, 61.16. HRMS (ESI) calcd for $C_{34}H_{32}NO_{10}$ $[M+H]^+$:614.2026, found 614.2038.

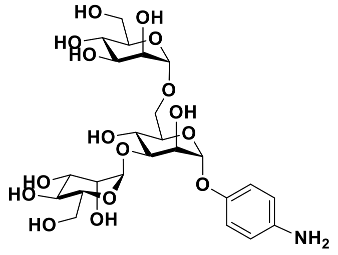

**Compound 48.**

To a stirred solution of **47** (0.2 mmol) and 4 A molecular sieve (0.2 g) in anhydrous DCM (2 mL) was cooled to -40°C and then $BF_3(OEt)_2$ (0.02 mmol) was added dropwise to the solution. A solution of **15a** in anhydrous DCM was added dropwise to the above mixture and stirred for 1 h at -40°C. After that, the reaction was gradually warmed to room temperature and stirred for another 1 h. The solution was quenched by adding triethylamine, then filtered, added sat. $NaHCO_3$ aq. and extracted with DCM. The organic layer was dried with $MgSO_4$ and evaporated to dryness. The residue was purified by flash column chromatography on silica gel to give trisacchride product. The product was then dissolved in MeOH and NaOMe (0.2 eq) was added and the resulting solution was stirred at rt for 2 h. The mixture was neutralized by IR-120, and then filtered, concentrated to dryness in vacuo. The deacetylated mixture was purified by Bio-Gel P-2 Gel (Biorad) with $H_2O$ as eluent to obtain a pure trisaccharide. The compound was lyophilized and then dissolved in MeOH (2 mL), 10% Pd-C (30 mg) was added and stirred vigorously under $H_2$ atmosphere overnight. The solution was filtered by celite and concentrated to dryness to give compound **48** (42%). $^1$H NMR (600 MHz, $D_2O$) δ 7.03 (d, $J$ = 9.2 Hz, 2H), 6.88 (d, $J$ = 9.2 Hz, 2H), 5.48 (s, 1H), 5.19 (s, 1H), 4.76 (s, 1H), 4.32 (s, 1H), 4.14 (dd, $J$ = 9.3, 3.0 Hz, 1H), 4.11 (s, 1H), 3.95-3.63 (m, 15H). $^{13}$C NMR (150 MHz, $D_2O$): 151.38, 143.39, 121.23, 120.70, 105.19, 101.60, 101.39, 80.93, 76.13, 75.38, 74.04, 73.27, 73.12, 72.79, 72.66, 72.22, 69.50, 69.41, 68.70, 67.90, 63.71, 63.65. HRMS (ESI) calcd for $C_{24}H_{37}NO_{16}Na[M+Na]^+$:618.2010, found 618.2029.

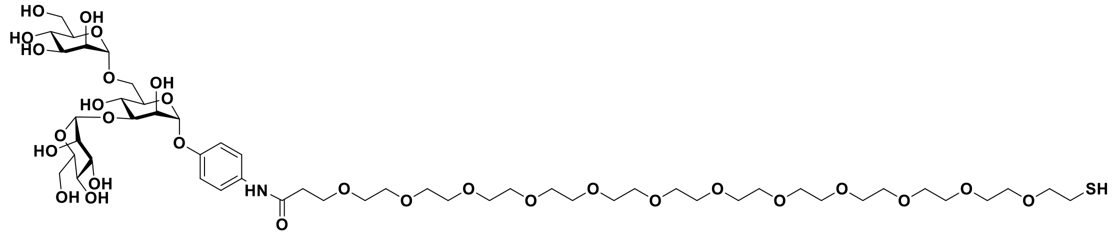

**Compound 49.**

Compound **48** (0.12 mmol) in DMF (1 mL) was added EDC (0.12 mmol), HOBt (0.12 mmol), DMAP(0.12 mmol), trimethylamine (0.2 mmol), and CT(PEG)$_{12}$ (0.1 mmol), and the resulting solution was stirred under nitrogen at rt for 12 h. The mixture was concentrated to dryness *in vacuo*, and the crude product was purified Bio-Gel P-2 Gel with $H_2O$ as eluent to to yield **49** (54%).$^1$H NMR (600 MHz, $D_2O$): 7.20 (d, $J$ = 9.2 Hz, 2H), 7.13 (d, $J$ = 9.2 Hz, 2H), 5.53 (s, 1H), 5.09 (s, 1H), 4.65 (s, 1H), 4.25 (s, 1H), 4.06 (dd, $J$ = 9.3, 3.0 Hz, 1H), 4.01 (m, 1H), 3.83-3.67 (m, 11H), 3.61-3.56 (m, 52H), 2.64 (t, $J$ = 6.4 Hz, 2H), 2.48 (t, $J$ = 6.4 Hz, 2H).$^{13}$C NMR (150 MHz, $D_2O$): 178.06, 153.84, 149.84, 123.02, 118.11, 120.42, 98.8, 97.78, 78.05, 73.39, 72.61, 72.17, 71.47, 70.52, 70.35, 70.01, 69.85, 69.55, 69.38, 69.31, 69.17, 66.97, 66.74, 66.60, 65.82, 65.10, 60.96, 60.86, 35.82, 23.03. HRMS (ESI) calcd for $C_{51}H_{89}NO_{29}SNa[M+Na]^+$:1234.5139, found 1234.5114.

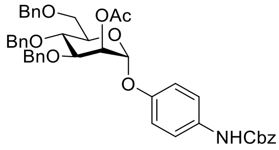


**Compound 51.**

To a stirred solution of **50** (1 mmol) in anhydrous DCM (10 mL) was added trichloroacetonitrile and DBU and the solution was stirred for 2 h at rt. The solvent was removed and the residue was purified by flash column chromatography on silica gel to give imidate product. To a stirred solution of benzyl (4-hydroxyphenyl)carbamate (1.2 mmol) and 4 A molecular sieve (1 g) in anhydrous DCM (10 mL) was cooled to -40°C and then $BF_3(OEt)_2$ (0.1 mmol) was added dropwise to the solution. A solution of imidate donor (1 mmol) in anhydrous DCM was added dropwise to the above mixture and stirred for 1 h at -40°C. After that, the reaction was gradually warmed to room temperature and stirred for another 1 h. The solution was quenched by adding triethylamine, then filtered, added sat. $NaHCO_3$ aq. and extracted with DCM. The organic layer was dried with $MgSO_4$ and evaporated to dryness. The residue was purified by flash column chromatography on silica gel to give compound **51** (72%). $^1$H NMR (600 MHz, $CDCl_3$) δ 7.40-7.28 (m, 18H), 7.19 (d, $J$ = 7.9 Hz, 2H), 7.10 (d, $J$ = 8.1 Hz, 2H), 7.00 (d, $J$ = 8.1 Hz, 2H), 5.57-5.55 (m, 2H), 5.12 (s, 2H), 4.91 (d, $J$ = 10.5 Hz, 1H), 4.80 (d, $J$ = 10.5 Hz, 1H), 4.69 (d, $J$ = 10.5 Hz, 1H), 4.65 (d, $J$ = 10.5 Hz, 1H), 4.53 (d, $J$ = 10.5 Hz, 1H), 4.46 (d, $J$ = 10.5 Hz, 1H), 4.22 (dd, $J$ = 9.4, 3.6 Hz, 1H), 4.07-4.04 (t, $J$ = 9.7 Hz, 1H), 3.93 (d, $J$ = 9.5 Hz, 1H), 3.83 (dd, $J$ = 10.9, 3.9 Hz, 1H), 3.68 (d, $J$ = 10.8 Hz, 1H), 2.21 (s, 3H).$^{13}$C NMR (150 MHz, $CDCl_3$):170.44, 156.25, 154.62, 138.27, 138.04, 137.82, 136.51, 132.71, 129.79, 128.50, 128.43, 128.31, 128.27, 128.10, 128.08, 127.83, 127.80, 127.64, 127.60, 116.63, 96.13, 77.95, 76.66, 75.22, 74.00, 73.34, 71.99, 71.92, 68.55, 68.5, 66.63, 21.09. HRMS (ESI) calcd for

$C_{43}H_{43}NO_{10}$ $[M+H]^+$:718.3016, found 718.3041.

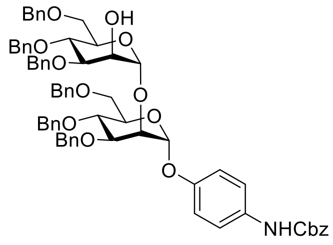


**Compound 52.**

To a stirred solution of **51** (0.6 mmol) in MeOH was added NaOMe (0.1 eq) and the resulting solution was stirred at rt for 1 h. The mixture was neutralized by IR-120, and then filtered, concentrated to dryness in vacuo. The deacetylated product was then dissolved in anhydrous DCM (5 mL) with 4 A molecular sieve (0.5 g) added. The solution was cooled to -40°C and then $BF_3(OEt)_2$ (0.05 mmol) was added dropwise to it. A solution of imidate donor (0.5 mmol) in anhydrous DCM was added dropwise to the above mixture and stirred for 1 h at -40°C. After that, the reaction was gradually warmed to room temperature and stirred for another 1 h. The solution was quenched by adding triethylamine, then filtered, added sat. $NaHCO_3$ aq. and extracted with DCM. The organic layer was dried with $MgSO_4$ and evaporated to dryness. The product was then dissolved in MeOH and NaOMe (0.1 eq) was added and the resulting solution was stirred at rt for 2 h. The mixture was neutralized by IR-120, and then filtered, concentrated to dryness in vacuo. The residue was purified by flash column chromatography on silica gel to give compound **52** (69%). $^1$H NMR (600 MHz, $CDCl_3$) δ 7.36-7.25 (m, 25H), 7.22-7.13 (m, 10H), 6.99-6.93 (m, 4H), 5.66 (s, 1H), 5.17 (s, 1H), 5.06 (s, 2H), 4.86 (d, *J* = 10.5 Hz, 1H), 4.79 (d, *J* = 10.5 Hz, 1H), 4.73, (s, 2H), 4.65 (d, *J* = 10.5 Hz, 1H), 4.58-4.57 (m, 2H), 4.55-4.52 (m, 2H), 4.47-4.43 (m, 3H), 4.19-4.14 (m, 2H), 3.99-3.96 (m, 2H), 3.88 (dd, *J* = 9.1, 3.1 Hz, 1H), 3.84-3.76 (m, 3H), 3.67-3.63 (m, 3H).$^{13}$C NMR (150 MHz, $CDCl_3$): 156.22, 154.70, 138.47, 138.35, 138.18, 138.14, 138.05, 137.90, 136.51, 132.30, 129.74, 129.63, 128.46, 128.44, 128.31, 128.28, 128.26, 128.19, 128.06, 127.93, 127.85, 127.83, 127.74, 127.68, 127.60, 127.54, 127.46, 127.38, 127.32, 116.63, 101.12, 96.95, 79.97, 79.42, 77.21, 77.00, 76.78, 75.12, 75.02, 74.66, 74.45, 74.33, 73.23, 73.16, 72.44, 72.41, 72.16, 71.69, 68.99, 68.45, 66.57. HRMS (ESI) calcd for $C_{68}H_{70}NO_{13}$ $[M+H]^+$:1108.4847, found 1108.4819.

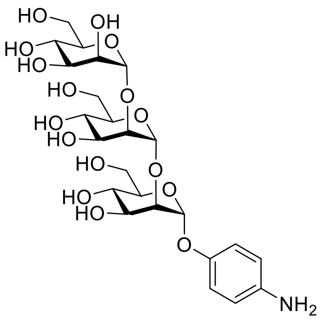


**Compound 53.**

To a stirred solution of **52** (0.3 mmol) in anhydrous DCM (2.5 mL) with 4 A molecular sieve (0.25 g) added. The solution was cooled to -40°C and then $BF_3(OEt)_2$ (0.03 mmol) was added dropwise to it. A solution of imidate donor (0.3 mmol) in anhydrous DCM was added dropwise to the above mixture and stirred for 1 h at -40°C. After that, the reaction was gradually warmed to room temperature and stirred for another 1 h. The solution was quenched by adding triethylamine, then filtered, added sat. $NaHCO_3$ aq. and extracted with DCM. The organic layer was dried with $MgSO_4$ and evaporated to dryness. The residue was purified by flash column chromatography on silica gel to give the trisaccharide product. The product was then dissolved in MeOH and NaOMe (0.1 eq) was added and the resulting solution was stirred at rt for 2 h. The mixture was neutralized by IR-120, and then filtered, concentrated to dryness in vacuo. The compound was then dissolved in MeOH (2 mL), 10% Pd-C (30 mg) was added and stirred vigorously under $H_2$ atmosphere overnight. The solution was filtered by celite and concentrated to dryness to give compound **53** (60%). $^1$H NMR (600 MHz, $D_2O$) δ 7.03 (d, *J* = 9.0 Hz, 2H), 6.83 (d, *J* = 9.0 Hz, 2H), 5.09 (s, 1H), 4.81 (s, 1H), 3.94-3.55 (m, 16H), 3.48 (t, *J* = 9.6 Hz, 1H), 3.29-3.27 (m, 1H).$^{13}$C NMR (150 MHz, $D_2O$): 151.36, 143.37, 121.20, 120.67, 101.59, 101.23, 96.33, 78.46, 75.34, 74.69, 73.51, 73.11, 72.97, 72.52, 71.53, 69.28, 69.14, 68.90, 65.51, 63.26, 62.91. HRMS (ESI) calcd for $C_{24}H_{38}NO_{16}$ $[M+H]^+$:596.2191, found 596.2044.

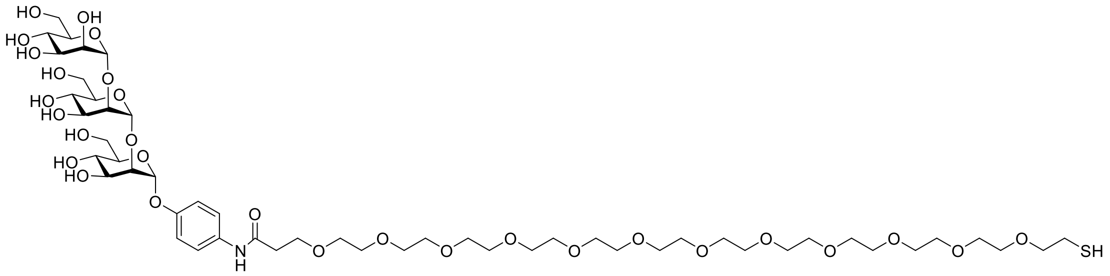


Compound **54**.

Compound **53** (0.02 mmol) in DMF (0.2 mL) was added EDC (0.02 mmol), HOBt (0.02 mmol), DMAP(0.02 mmol), trimethylamine (0.04 mmol), and $CT(PEG)_{12}$ (0.02 mmol), and the resulting solution was stirred under nitrogen at rt for 12 h. The mixture was concentrated to dryness *in vacuo*, and the crude product was purified Bio-Gel P-2 Gel with $H_2O$ as eluent to to yield **54** (58%).$^1$H NMR (600 MHz, $D_2O$): 7.03 (d, $J$ = 9.2 Hz, 2H), 6.82 (d, $J$ = 9.2 Hz, 2H), 5.09 (s, 1H), 4.81 (s, 1H), 3.84-3.46 (m, 65H), 3.30-3.26 (m, 1H), 2.65 (t, $J$ = 6.5 Hz, 2H), 2.52 (s, 2H).$^{13}$C NMR (150 MHz, $D_2O$):173.22, 151.36, 143.37, 121.37, 120.59, 101.20, 101.67, 97.04, 79.17, 76.06, 75.40, 75.22, 74.23, 73.69, 73.23, 72.60, 72.45, 72.22, 71.68, 71.21, 69.85, 69.61, 63.97, 38.52, 26.08. HRMS (ESI) calcd for $C_{51}H_{90}NO_{29}S[M+H]^+$:1212.5319, found 1212.5146.

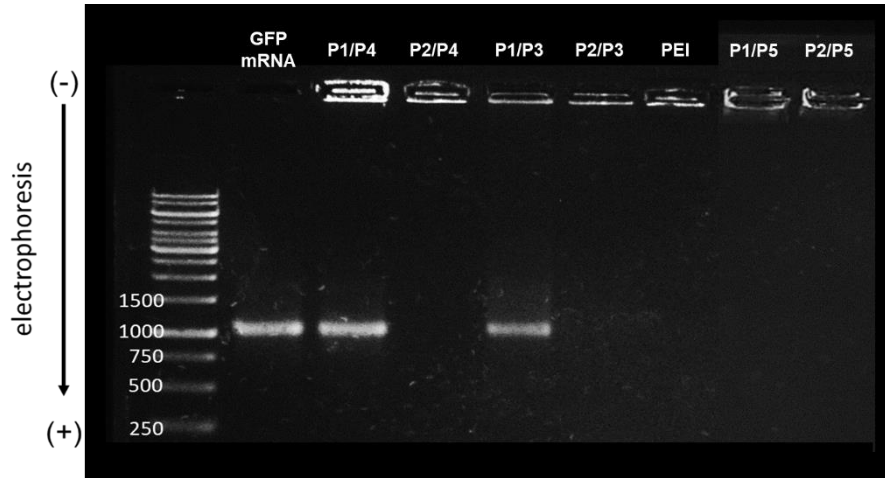


**Figure S1.** Agarose gel electrophoresis assay of copolymers with GFP mRNA (N/P ratio = 3:1).

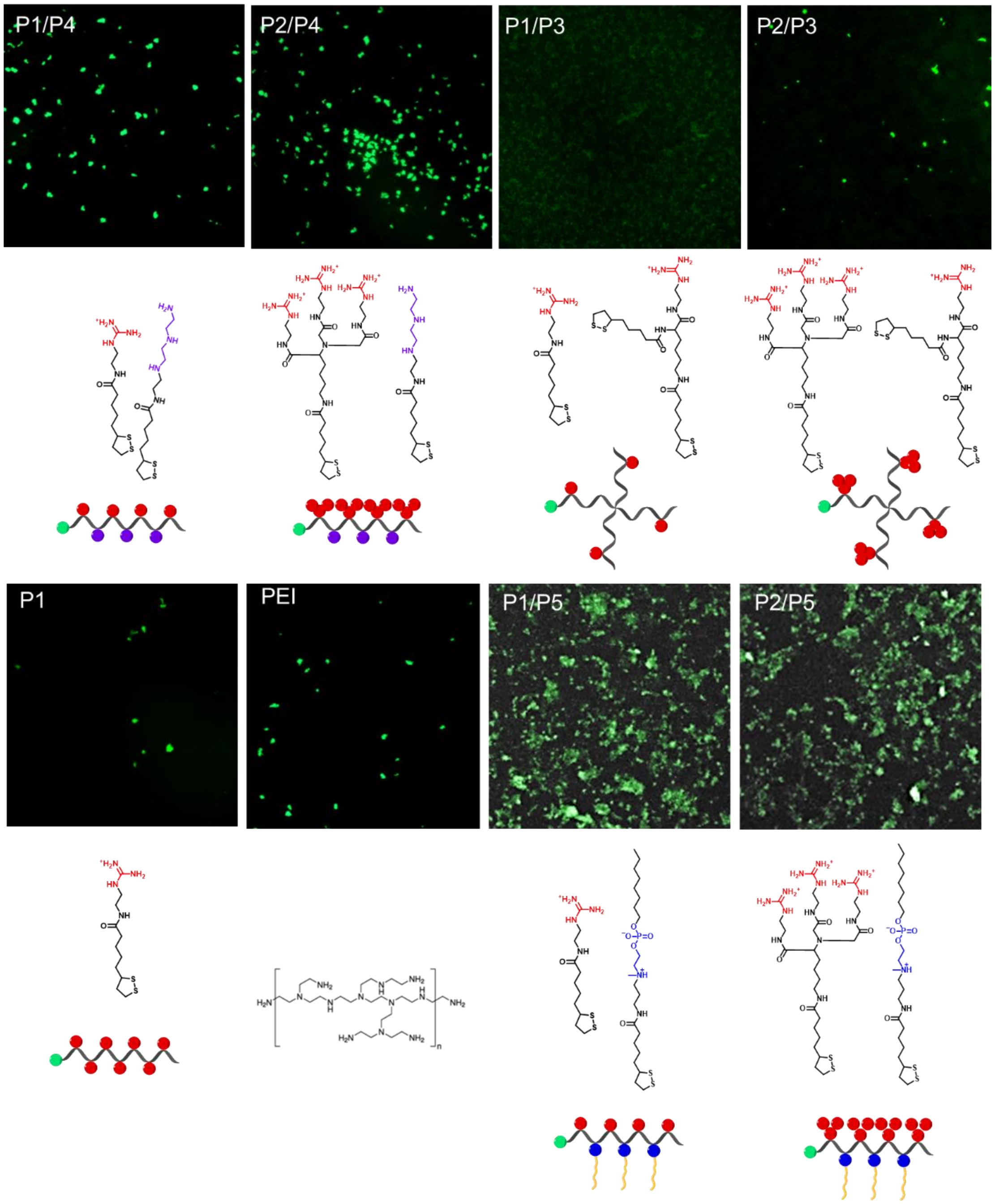


**Figure S2.** The polymers used in this study and the fluorescent image of GFP protein translated by GFP mRNA using different poly(disulfide)s for encapsulation in HEK293T cells are shown. GFP mRNA was complexed with **P1/P4, P2/P4, P1/P3, P2/P3, P1, PEI** (MW: 25kDa)**, P1/P5,** or **P2/P5** .

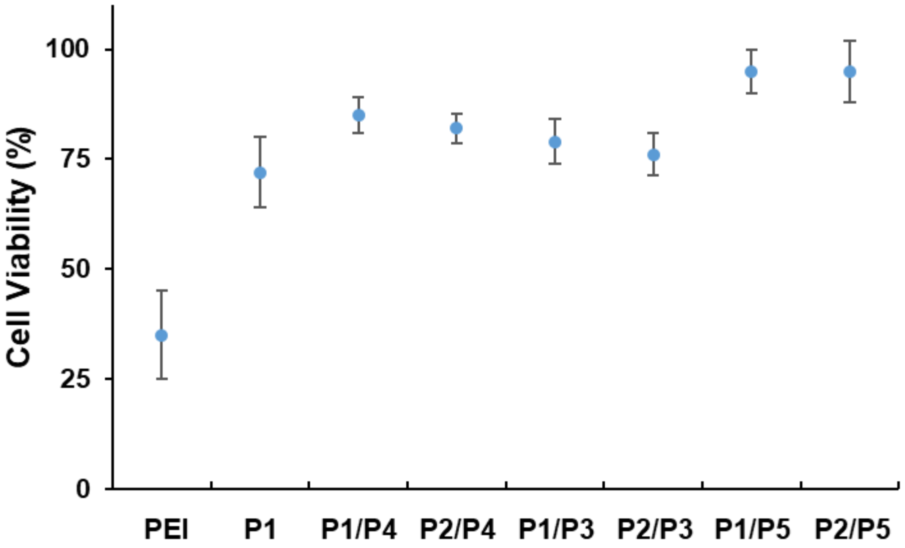


**Figure S3.** Cell viability assay of HEK293T cells after PNP treatment. The N/P ratio of mRNA to **Px** was fixed at 3:1 and 1 µg/well mRNA was used. Error bar represent the standard error (mean ± S.D.,n=3)

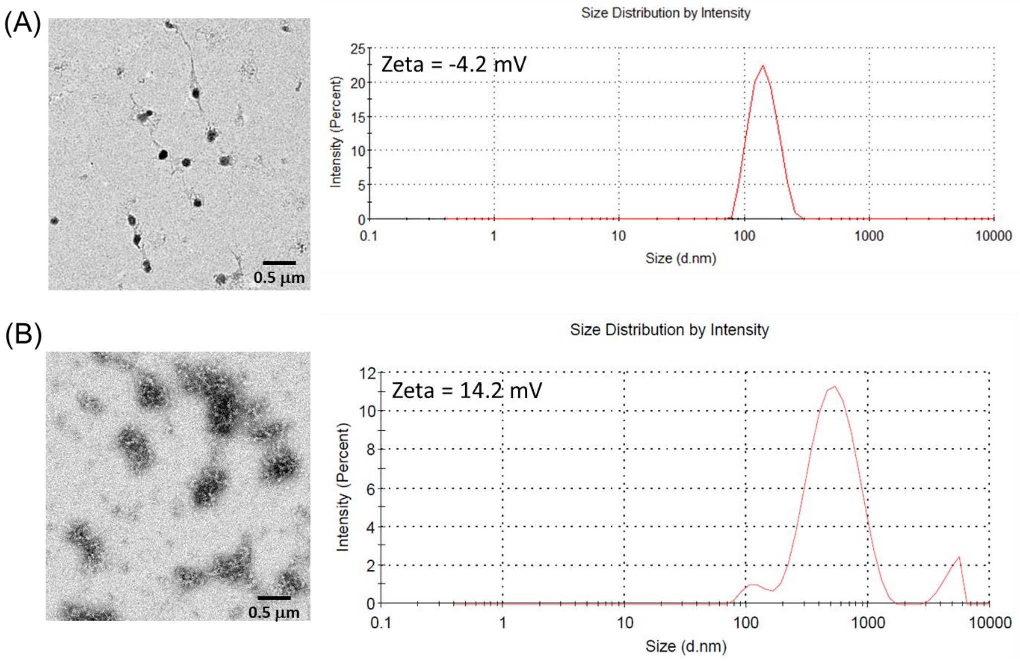


**Figure S4.** Size and zeta potential of the (A) mRNA-PNP (**I1-P1/P5**) (B) mRNA-PNP (**I1-P1**) nanoparticles measured by TEM and dynamic light scattering. All the samples and experiments were maintained at 25 °C.

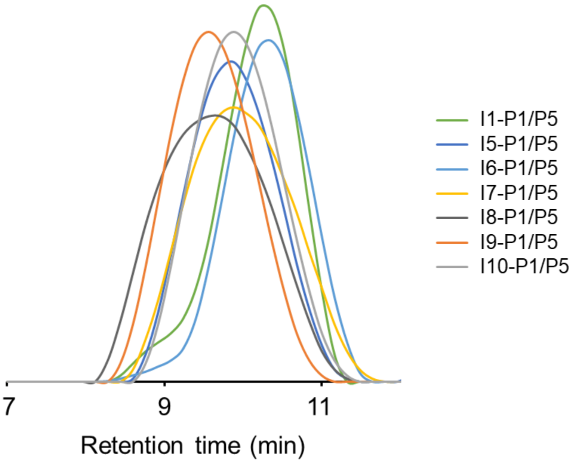


| Samples | MW (kDa) | PDI |
|---|---|---|
| I1-P1/P5 | 10.2 | 1.33 |
| I5-P1/P5 | 11.8 | 1.24 |
| I6-P1/P5 | 11.2 | 1.39 |
| I7-P1/P5 | 12.6 | 1.61 |
| I8-P1/P5 | 14.1 | 1.17 |
| I9-P1/P5 | 13.8 | 1.37 |
| I10-P1/P5 | 12.3 | 1.22 |

**Figure S5.** GPC characterization and determination of PDI and MW of synthesized polymers.

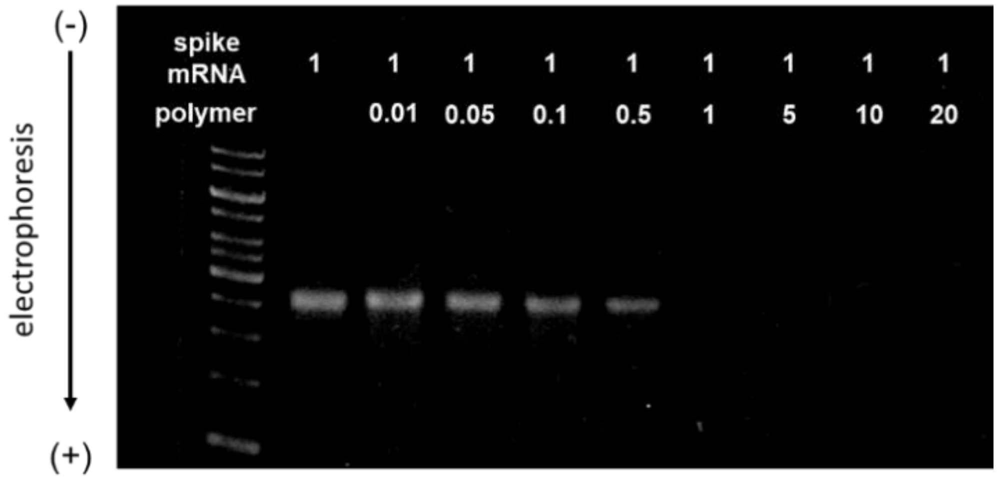


**Figure S6.** Agarose gel electrophoresis assay of spike mRNA-PNP at different N/P ratio, the ratio of positively-chargeable polymer amine (N = nitrogen) groups to negatively-charged nucleic acid phosphate (P) groups

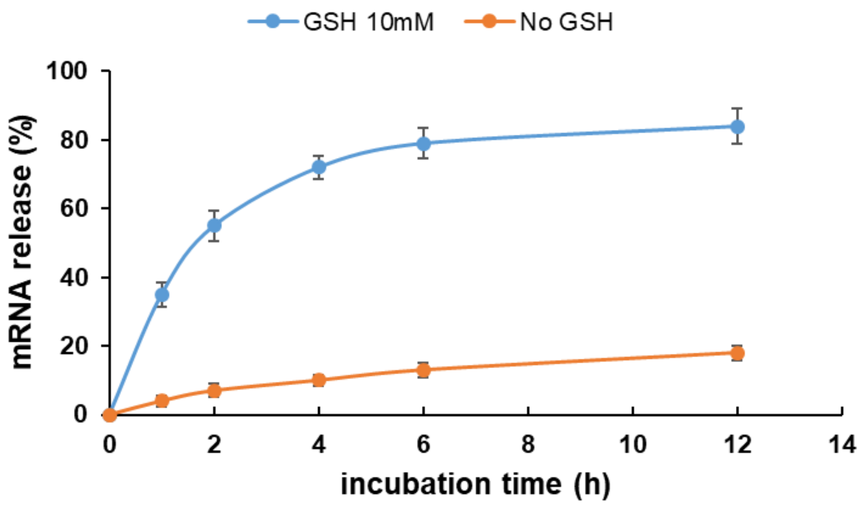


**Figure S7.** Detection of spike mRNA release from mRNA-PNP (**I1-P1/P5**) in addition of GSH (10 mM) or in PBS buffer for 0-12 h.

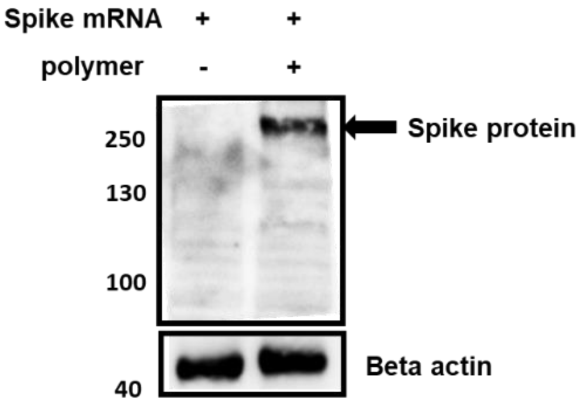


**Figure S8.** Chemiluminescent imaging of spike protein expression mediated by spike mRNA-PNP (**I1-P1/P5**) in HEK293T cells.

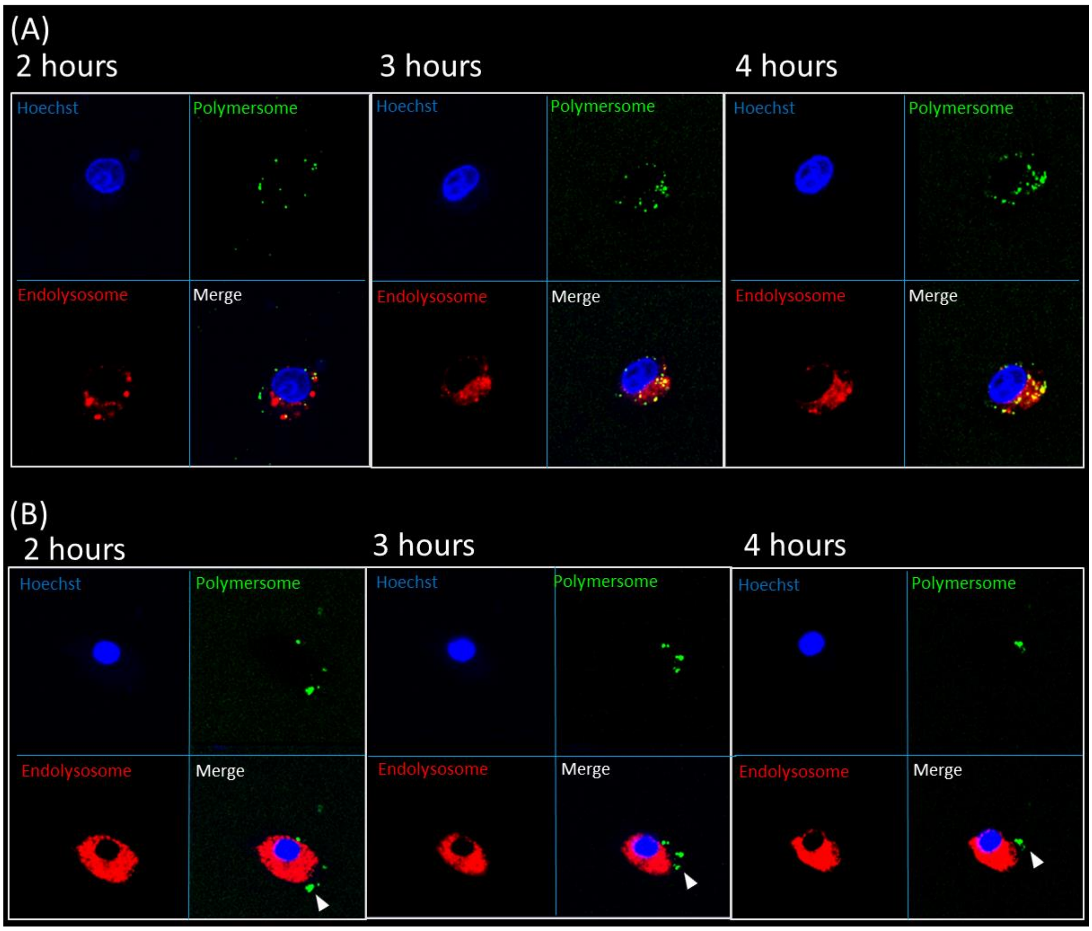


**Figure S9.** Endosomal escape and cellular uptake fluorescence images of dendritic cells treated with spike mRNA-PNP for 2-4 h (A) mRNA-PNP **(I1-P1/P4)** (B) mRNA-PNP(**I1-P1/P4/P5)**. FITC conjugated-mRNA-PNP (green), endo/lysosomes (red), and nuclei (blue) were shown in the images. The white arrow indicated mRNA escape from endo/lysosomes.

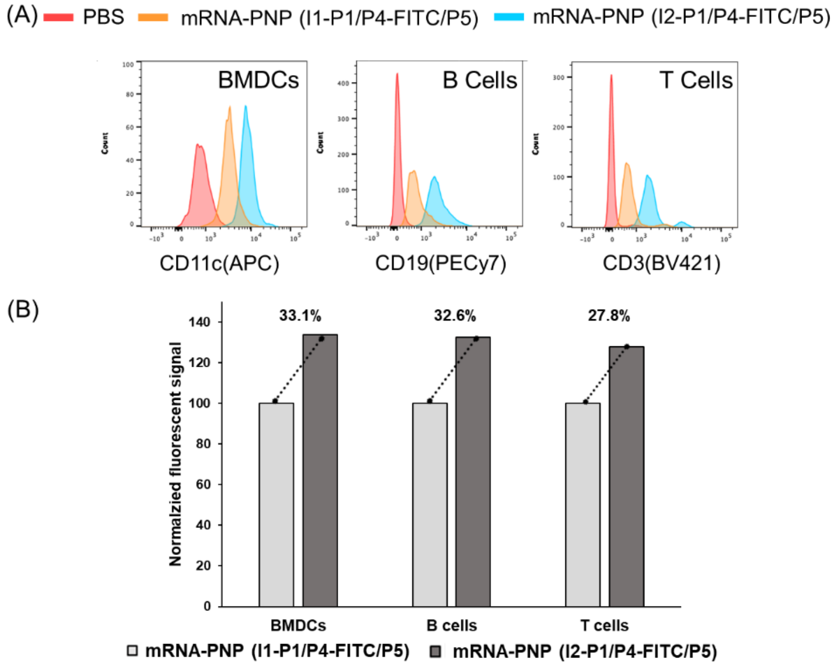


**Figure S10.** (A) Flow cytometry analysis revealed increased uptake of Siglec-2-targeting PNP (blue) to BMDCs, B cells, and T cells from mouse splenocytes compared to nontargeted PNP (orange) with PBS as control (pink). (B) Cellular uptake fluorescence signal of BMDC, B cells, and T cells treated with spike mRNA-PNP (I1-P1/P4-FITC/P5) or spike mRNA-PNP (I2-P1/P4-FITC/P5) after 1 h.

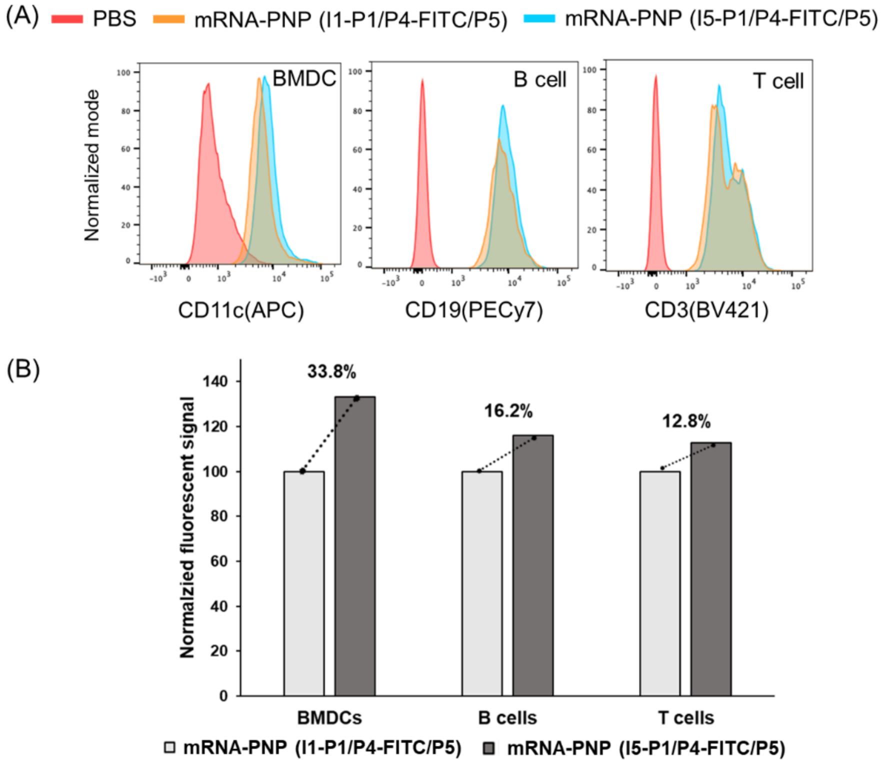

**Figure S11.** (A) Flow cytometry analysis revealed increased uptake of DC-SIGN-targeted PNP (blue) to BMDCs, B cells, and T cells from mouse splenocytes compared to non-targeted PNP (orange) with PBS as control (pink). (B) Cellular uptake of spike mRNA-PNP (I1-P1/P4-FITC/P5) or spike mRNA-PNP (I5-P1/P4-FITC/P5) after 1 h incubation with BMDC, B cells, and T cells and measured by fluorescence signal.

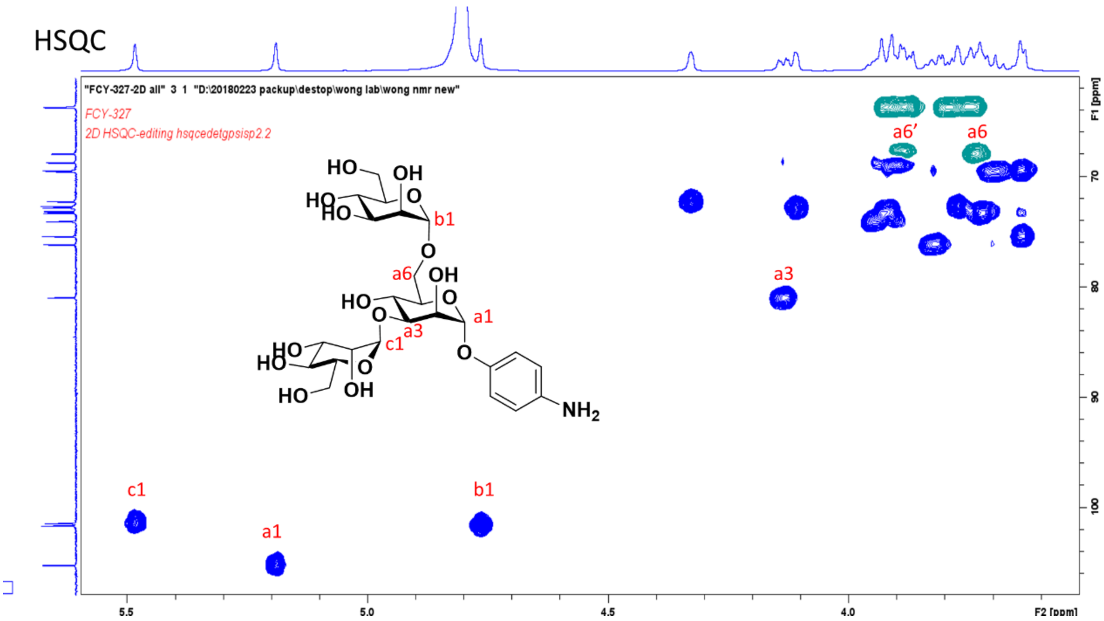


**Figure S12.** HSQC NMR spectrum of phenyltrimannose **48**.

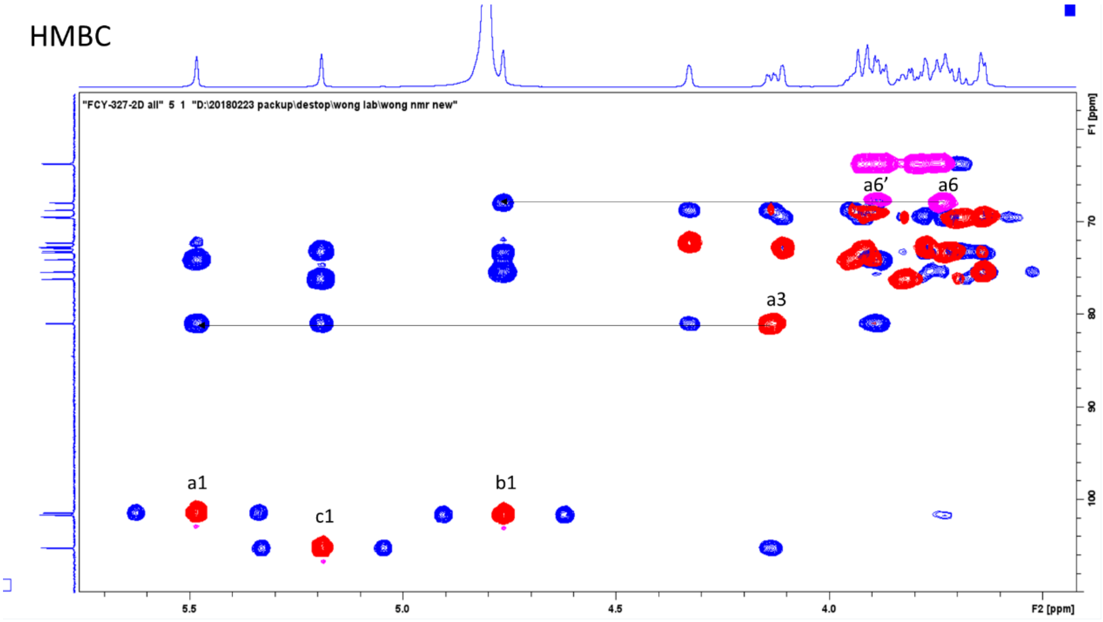

**Figure S13.** HMBC NMR spectrum of phenyltrimannose **48**.

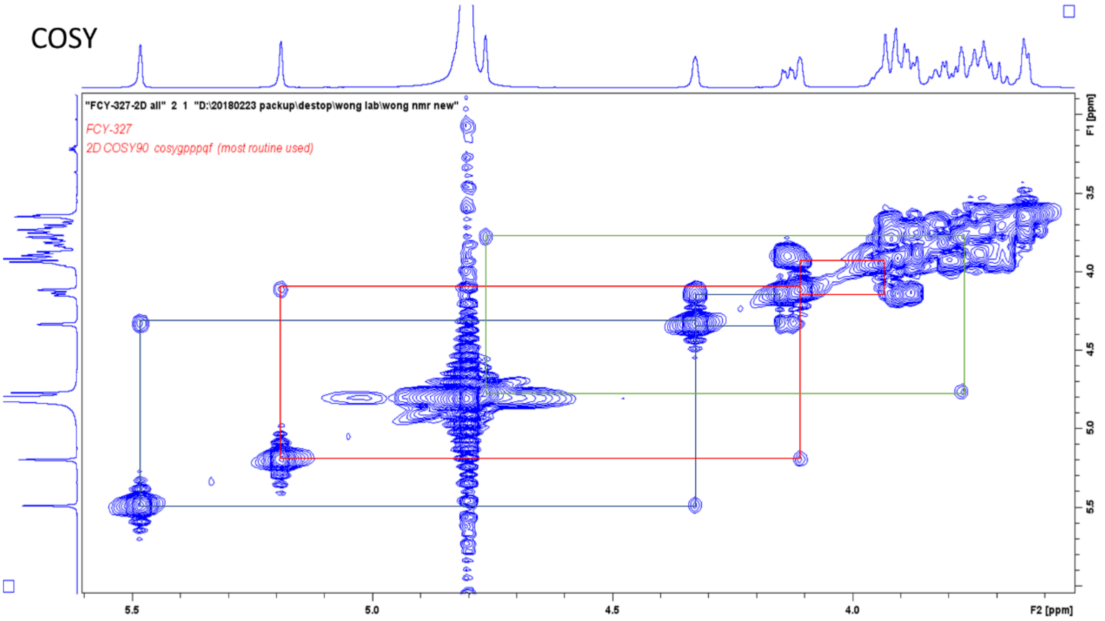


**Figure S14.** 2D COSY NMR spectrum of phenyltrimannose **48**.

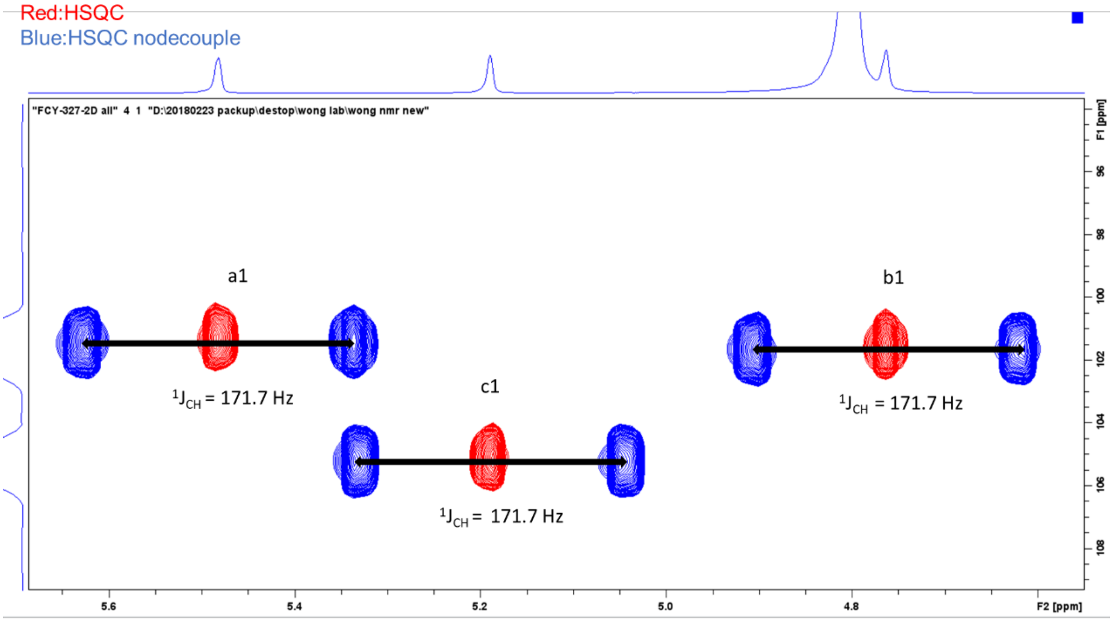


**Figure S15.** HSQC and HSQC nodecouple mode NMR spectrum of phenyltrimannose **48**.

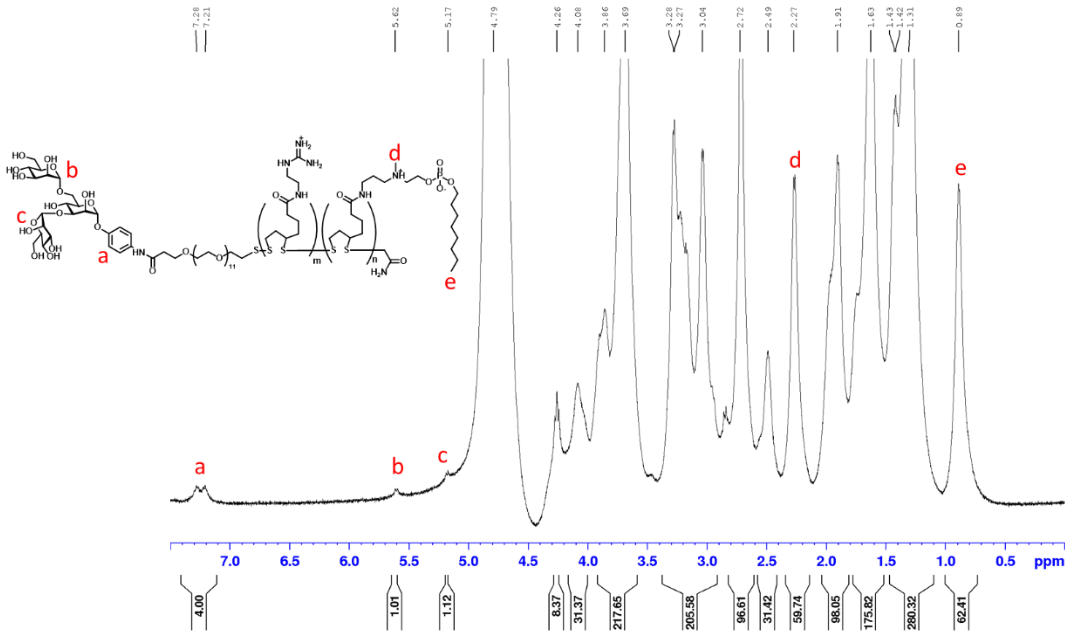


**Figure S16.** $^{1}$H NMR characterization of synthesized copolymers **I9-P1/P5**.

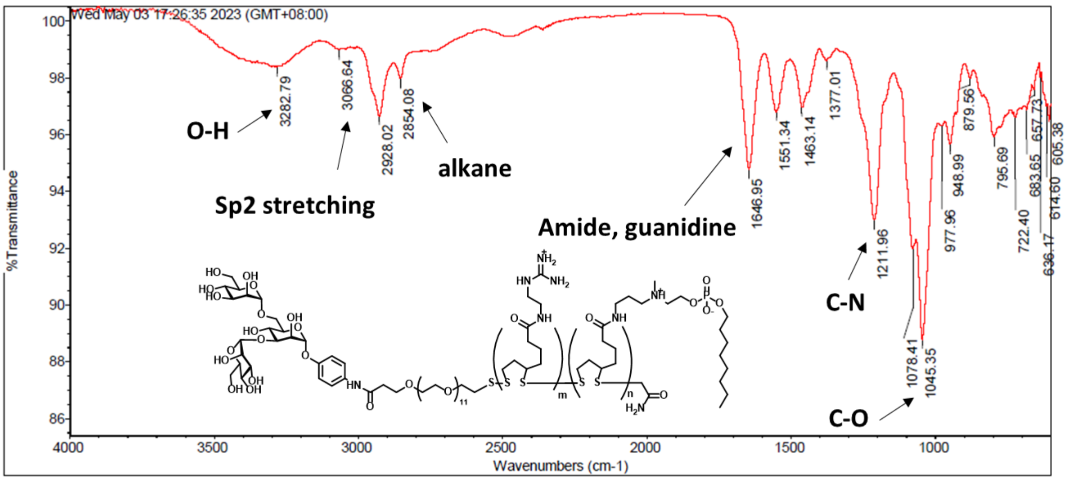


**Figure S17.** IR characterization of synthesized copolymers **I9-P1/P5**.

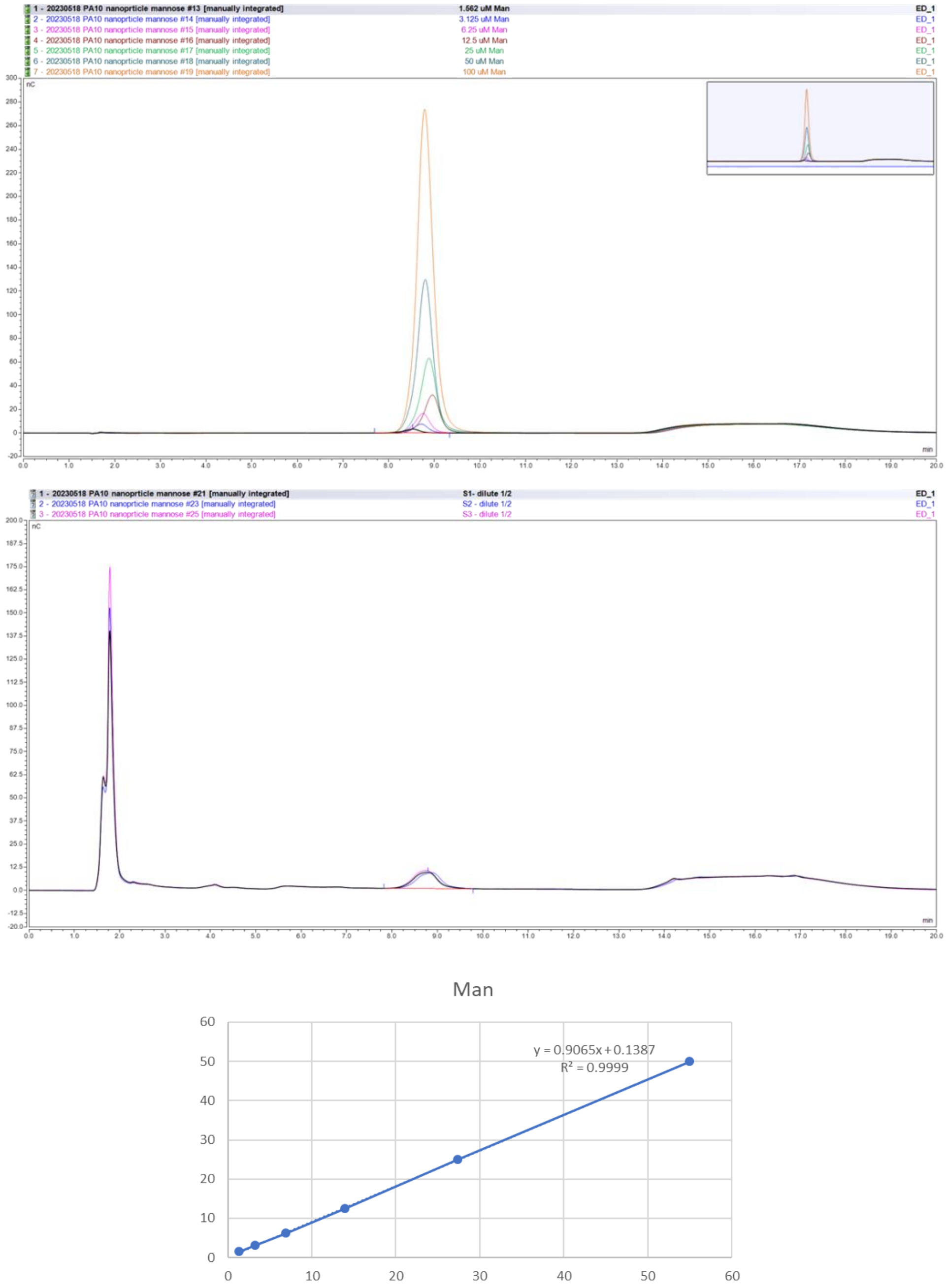


| | con (uM) | 1.562 | 3.125 | 6.25 | 12.5 | 25 | 50 |
|---|---|---|---|---|---|---|---|
| (Area) | Man | 1.3615 | 3.2053 | 6.9001 | 13.9073 | 27.3202 | 54.9801 |

| Peak Name | Ret.Time( min) | Area( nC*min) | Conc. ( µM) |
|---|---|---|---|
| S1 dilute 1/2 | 8.792 | 5.6082 | 5.22 |
| S2 dilute 1/2 | 8.875 | 5.4404 | 5.07 |
| S3 dilute 1/2 | 8.767 | 6.1019 | 5.67 |

| Sample Name | Conc. ( µM) | nmol/mg NP |
|---|---|---|
| S1 | 10.45 | 107.1 |
| S2 | 10.14 | 103.9 |
| S3 | 11.34 | 116.2 |

**Figure S18.** Glycan quantification on mRNA-PNP (I9-P1/P5). (a) HPAEC-PAD chromatogram recorded with the peak of mannose appearing at the retention time of 8.8 min (b) standard curve of free mannose and the calculation for surface glycan on

PNP.

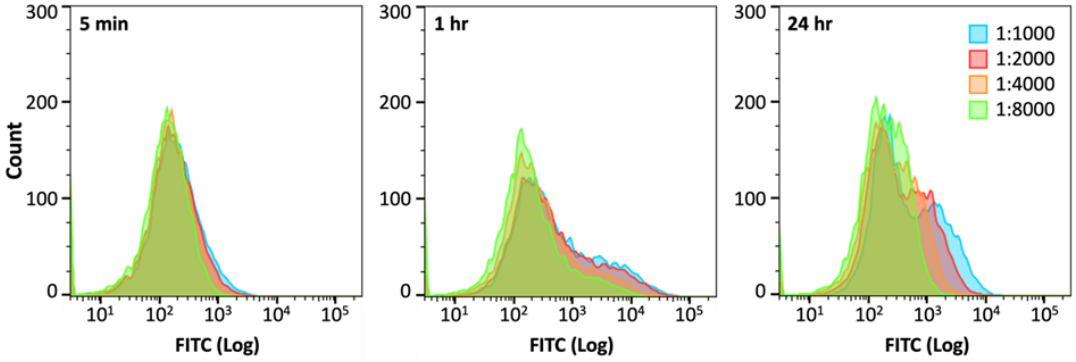


**Figure S19.** Flow cytometry analysis of FITC signals at different dilutions and incubation time points with mRNA-PNP (I9-P1/P5). mRNA-PNP concentration (dilution compared to 10 mg/mL stock): 1:1000, 1:2000, 1:4000, 1:8000. Incubation time: 5 min, 1 hr, 24 hr.

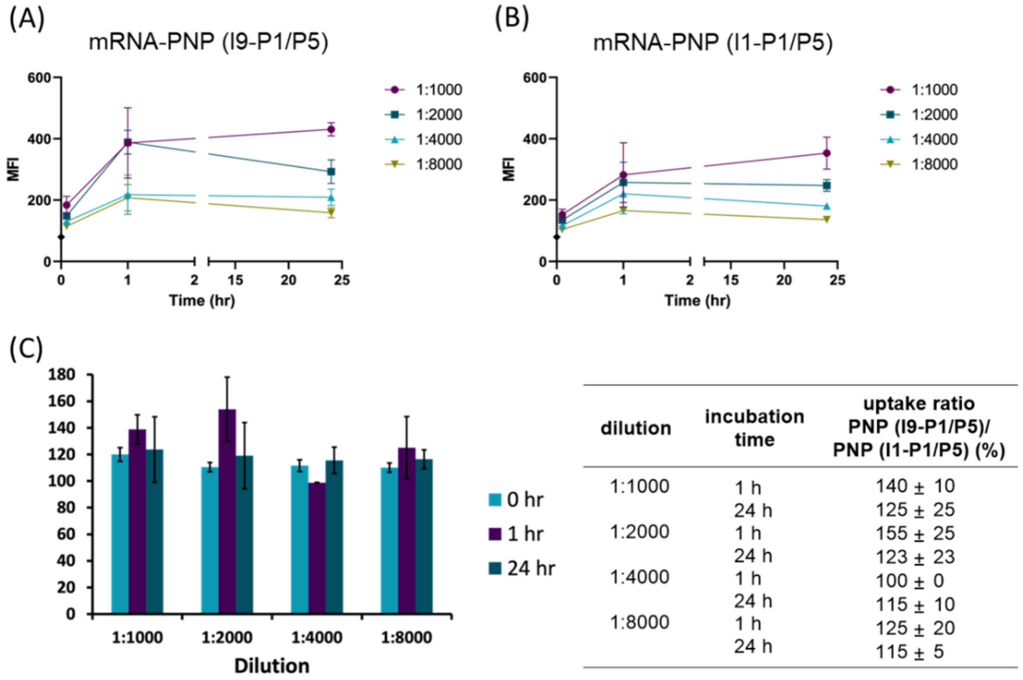


| dilution | incubation time | uptake ratio PNP (I9-P1/P5)/ PNP (I1-P1/P5) (%) |
|---|---|---|
| 1:1000 | 1 h | 140 ± 10 |
| | 24 h | 125 ± 25 |
| 1:2000 | 1 h | 155 ± 25 |
| | 24 h | 123 ± 23 |
| 1:4000 | 1 h | 100 ± 0 |
| | 24 h | 115 ± 10 |
| 1:8000 | 1 h | 125 ± 20 |
| | 24 h | 115 ± 5 |

**Figure S20.** FITC signals of BMDCs after mRNA-PNP incubation. BMDC incubation with (A) mRNA-PNP (I9-P1/P5) and (B) mRNA-PNP (I1-P1/P5) under different dilutions at 5 min, 1 hr and 24 hr. Each data point is the averaged geometric mean of FITC signals (MFI) measured by flow cytometry under the same conditions done on the same day (N = 2), and the error bars represent standard deviations. (C) Averaged ratios of mRNA-PNP (I9-P1/P5) and mRNA-PNP (I1-P1/P5) FITC signals (N = 2), and the error bars represent standard deviations.

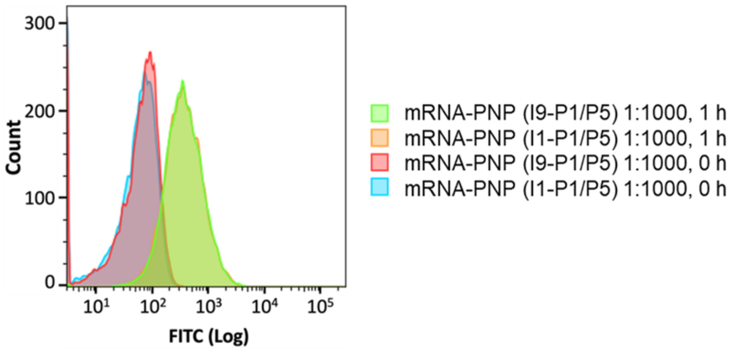


**Figure S21.** C2C12 myoblast FITC signal after incubation with mRNA-PNP measured by flow cytometry.

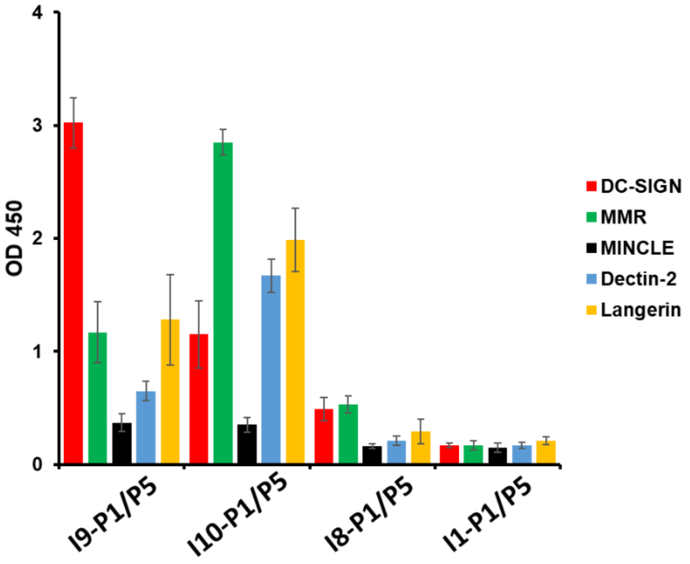


**Figure S22.** Binding analysis of DC-SIGN, MMR, MINCLE, Dectin-2 and Langerin (0.625 µg/mL) at pH 7.4 to PNP.

**References.**

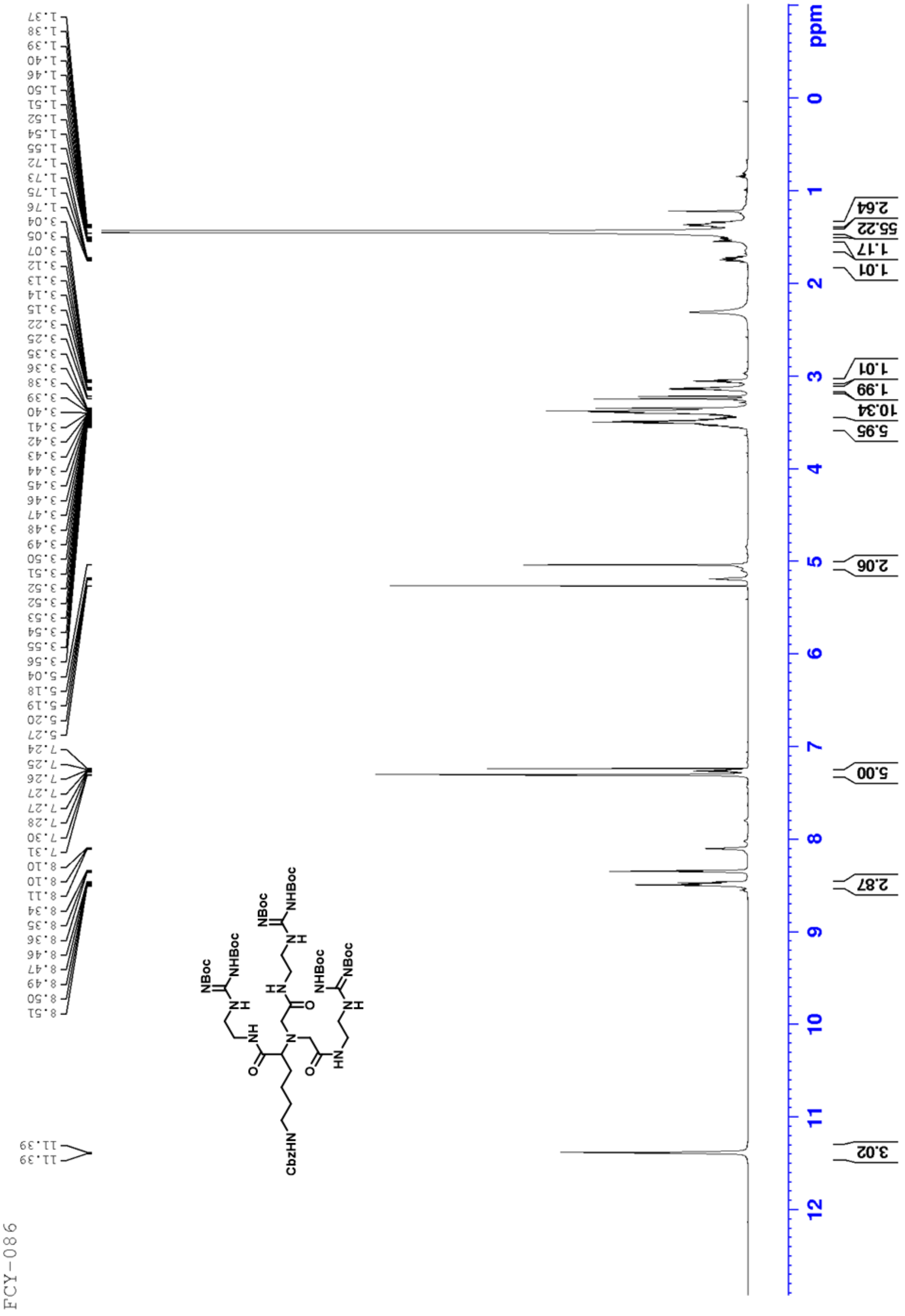
FCY-086
1.37 1.38 1.39 1.40 1.46 1.50 1.51 1.52 1.54 1.55 1.72 1.73 1.75 1.76 3.04 3.05 3.07 3.12 3.13 3.14 3.15 3.22 3.25 3.35 3.36 3.38 3.39 3.40 3.41 3.42 3.43 3.44 3.45 3.46 3.47 3.48 3.49 3.50 3.51 3.52 3.52 3.53 3.54 3.55 3.56 5.04 5.18 5.19 5.20 5.27 7.24 7.25 7.26 7.27 7.27 7.28 7.30 7.31 8.10 8.10 8.11 8.34 8.35 8.36 8.46 8.47 8.49 8.50 8.51 11.39 11.39
CbzHN
NH
NBoc
NHBoc
HN
O
N
ppm
2.64
55.22
1.17
1.01
1.01
1.99
10.34
5.95
2.06
5.00
2.87
3.02

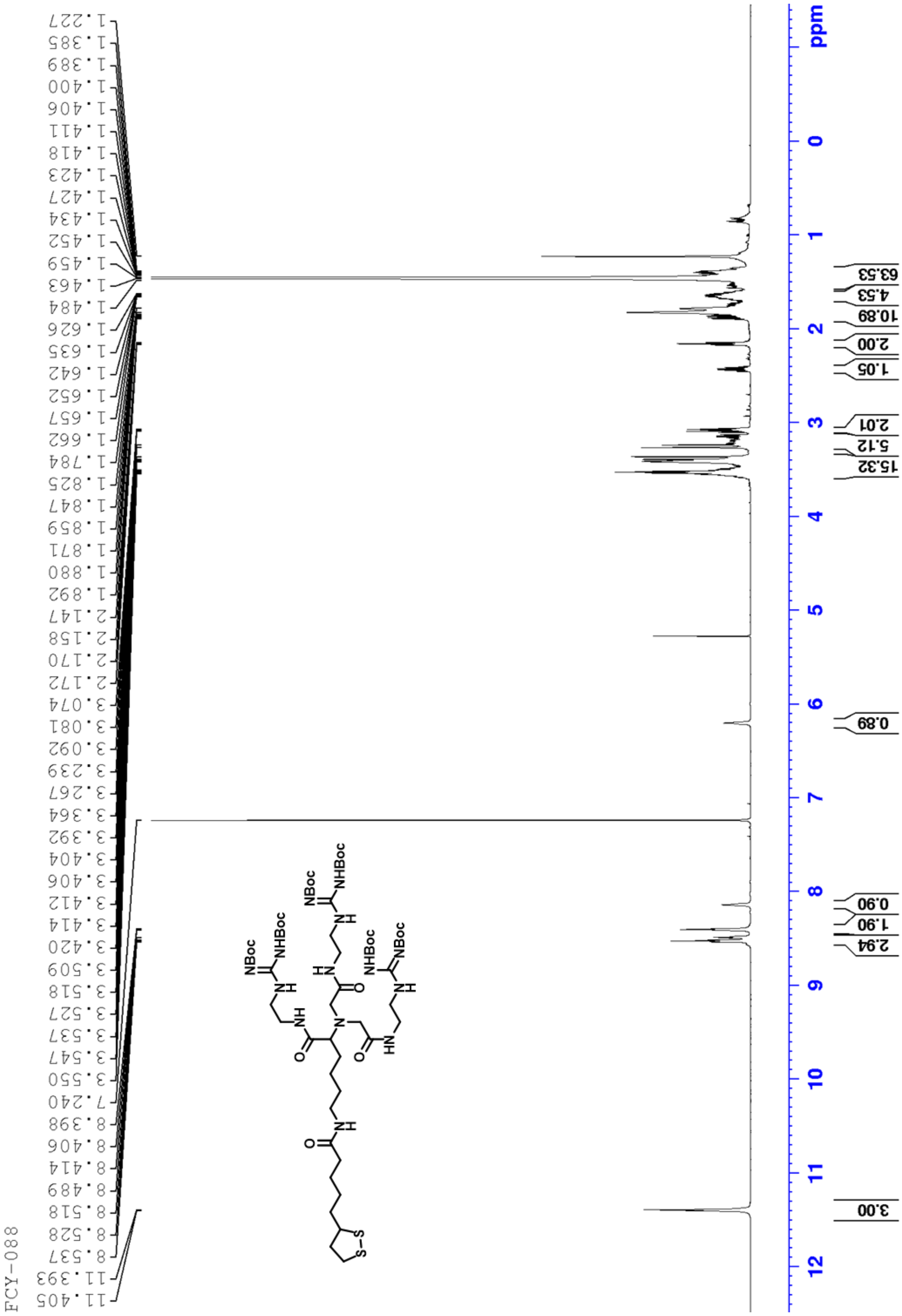
FCY-088
1.227
1.385
1.389
1.400
1.406
1.411
1.418
1.423
1.427
1.434
1.452
1.459
1.463
1.484
1.626
1.635
1.642
1.652
1.657
1.662
1.784
1.825
1.847
1.859
1.871
1.880
1.892
2.147
2.158
2.170
2.172
3.074
3.081
3.092
3.239
3.267
3.364
3.392
3.404
3.406
3.412
3.414
3.420
3.509
3.518
3.527
3.537
3.547
3.550
7.240
8.398
8.406
8.414
8.489
8.518
8.528
8.537
11.393
11.405
NBoc
NHBoc
S-S
63.53
4.53
10.89
2.00
1.05
2.01
5.12
15.32
0.89
0.90
1.90
2.94
3.00
ppm

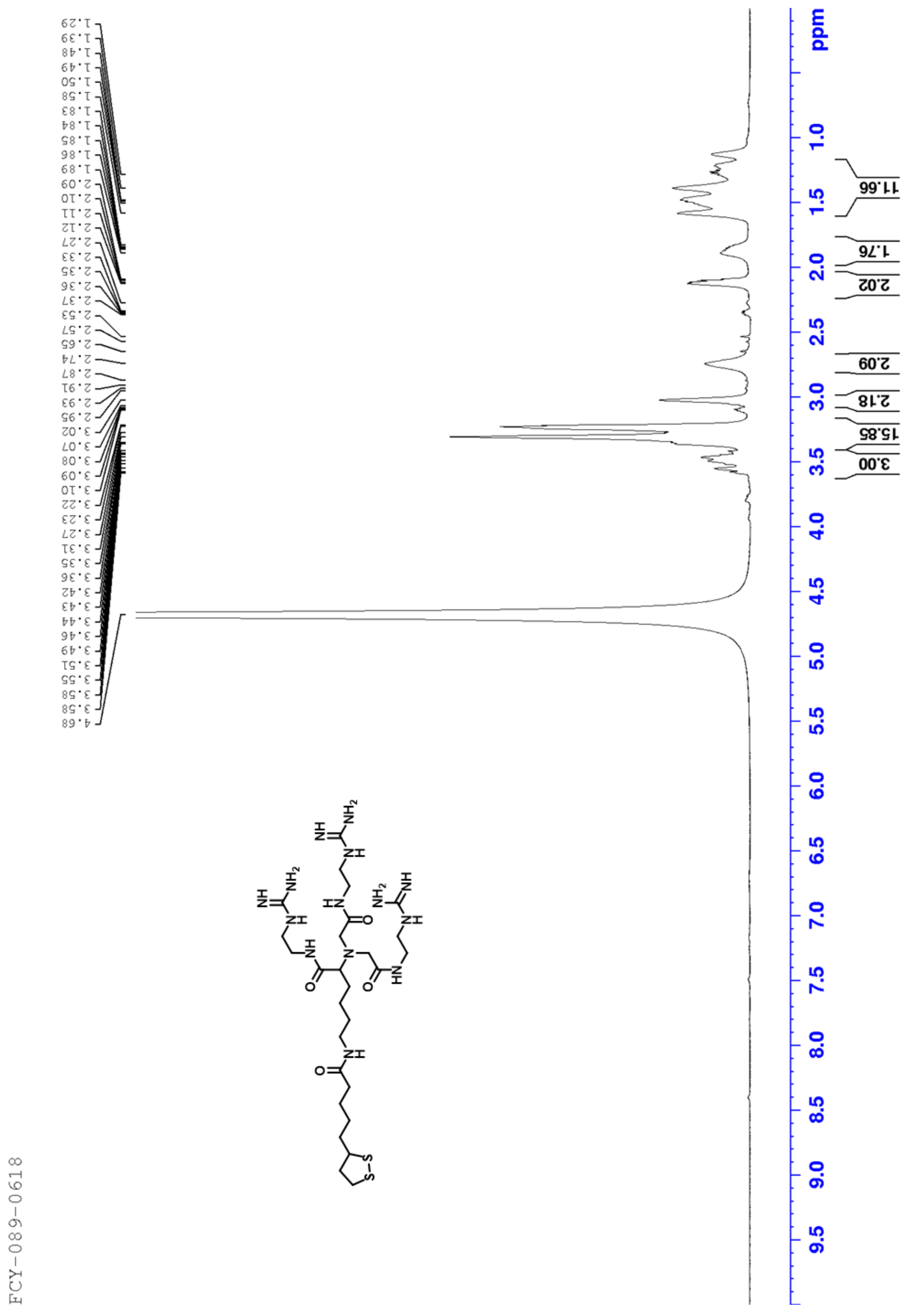

FCY-089-0618
1.29 1.39 1.48 1.49 1.50 1.58 1.83 1.84 1.85 1.86 1.89 2.09 2.10 2.11 2.12 2.27 2.33 2.35 2.36 2.37 2.53 2.57 2.65 2.74 2.87 2.91 2.93 2.95 3.02 3.07 3.08 3.09 3.10 3.22 3.23 3.27 3.31 3.35 3.36 3.42 3.43 3.44 3.46 3.49 3.51 3.55 3.58 3.58 4.68
11.66
1.76
2.02
2.09
2.18
15.85
3.00
ppm
1.0 1.5 2.0 2.5 3.0 3.5 4.0 4.5 5.0 5.5 6.0 6.5 7.0 7.5 8.0 8.5 9.0 9.5

FCY-089-0618-C

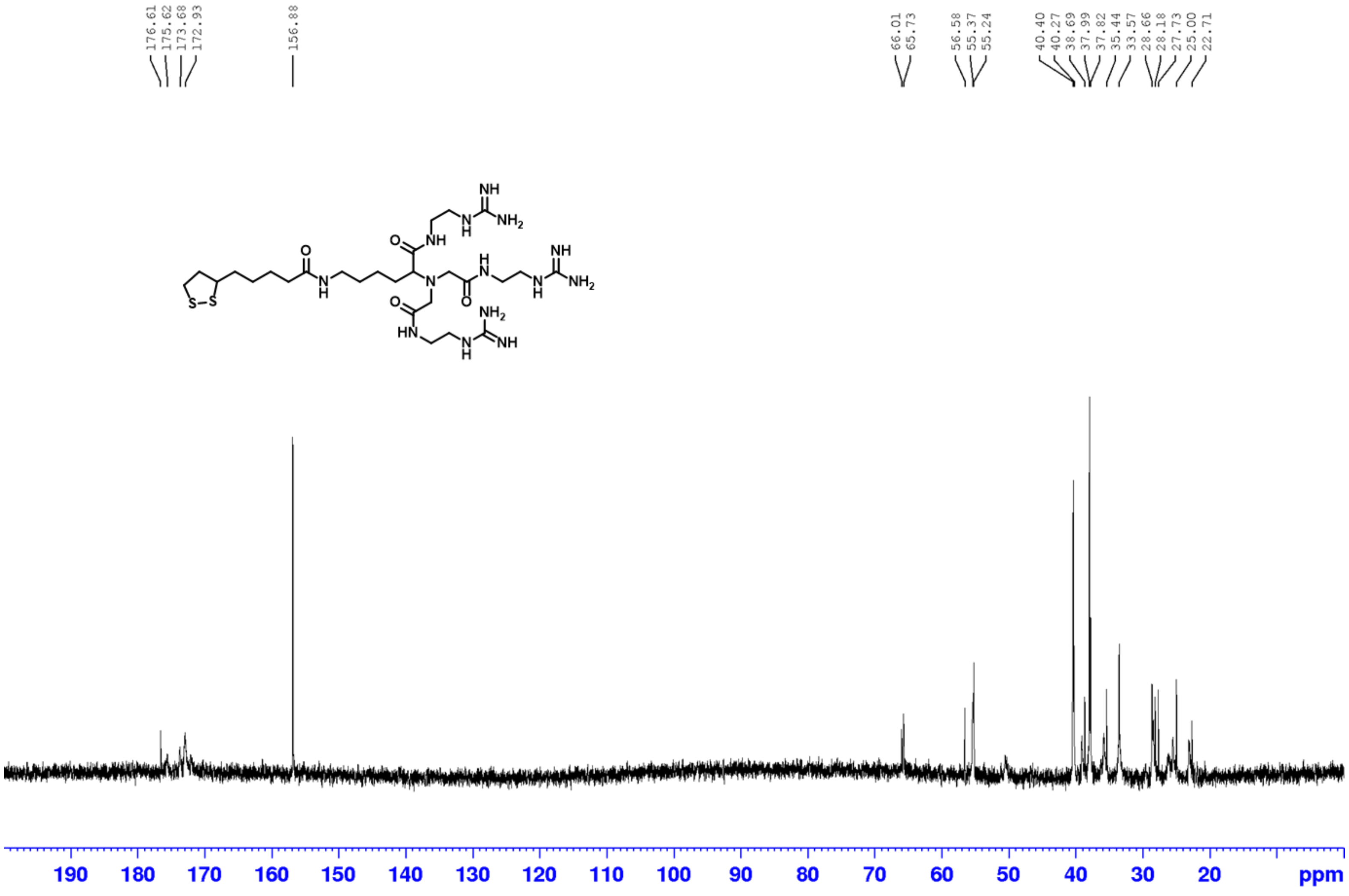

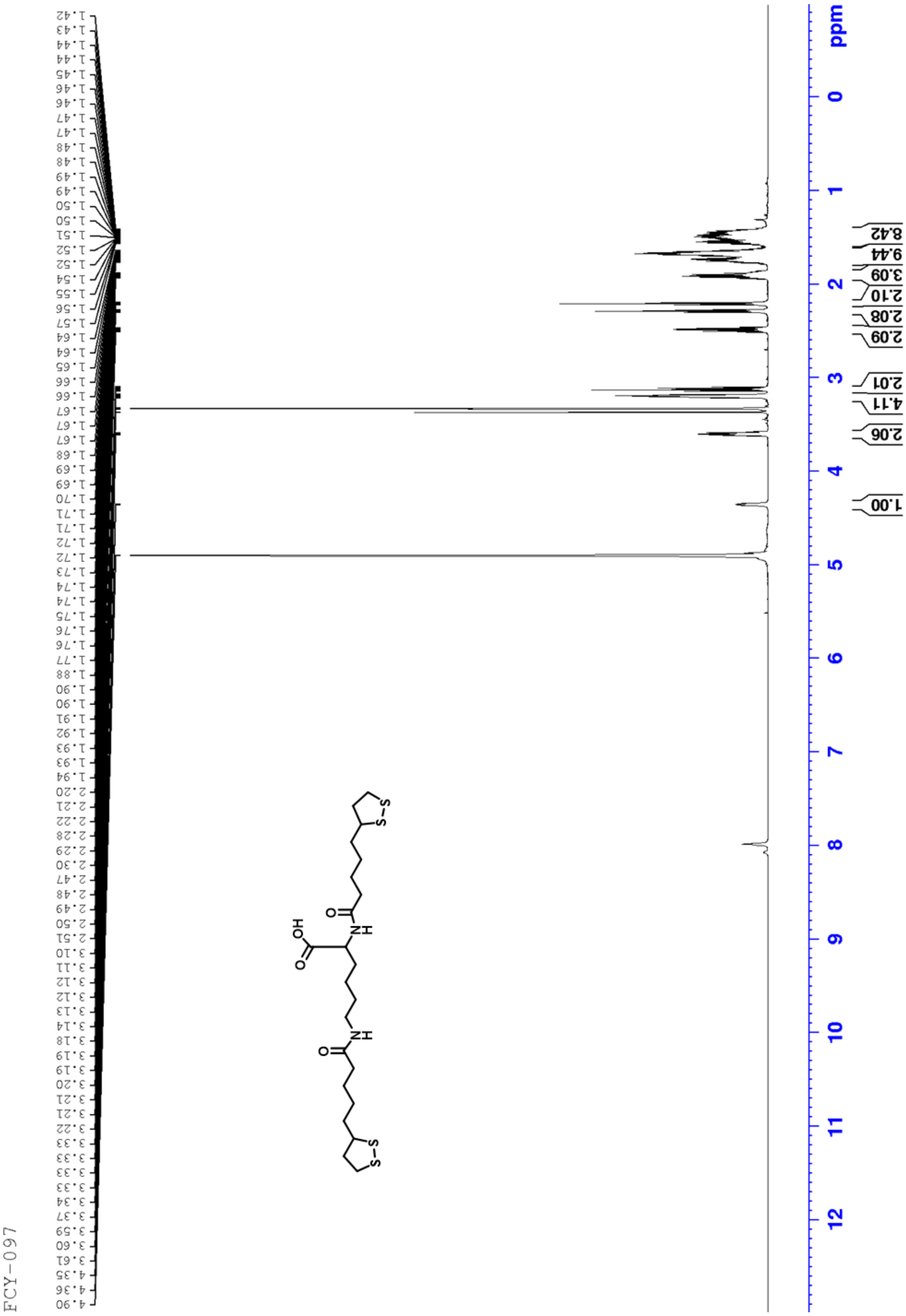
FCY-097
1.42
1.43
1.44
1.44
1.45
1.46
1.46
1.47
1.47
1.48
1.48
1.49
1.49
1.50
1.50
1.51
1.52
1.52
1.54
1.55
1.56
1.57
1.64
1.64
1.65
1.66
1.66
1.67
1.67
1.67
1.68
1.69
1.69
1.70
1.71
1.71
1.72
1.72
1.73
1.74
1.74
1.75
1.76
1.76
1.77
1.88
1.90
1.90
1.91
1.92
1.93
1.93
1.94
2.20
2.21
2.22
2.28
2.29
2.30
2.47
2.48
2.49
2.50
2.51
3.10
3.11
3.12
3.12
3.13
3.14
3.18
3.19
3.19
3.20
3.21
3.21
3.22
3.33
3.33
3.33
3.33
3.34
3.37
3.59
3.60
3.61
4.35
4.36
4.90
8.42
9.44
3.09
2.10
2.08
2.09
2.01
4.11
2.06
1.00
ppm
0
1
2
3
4
5
6
7
8
9
10
11
12

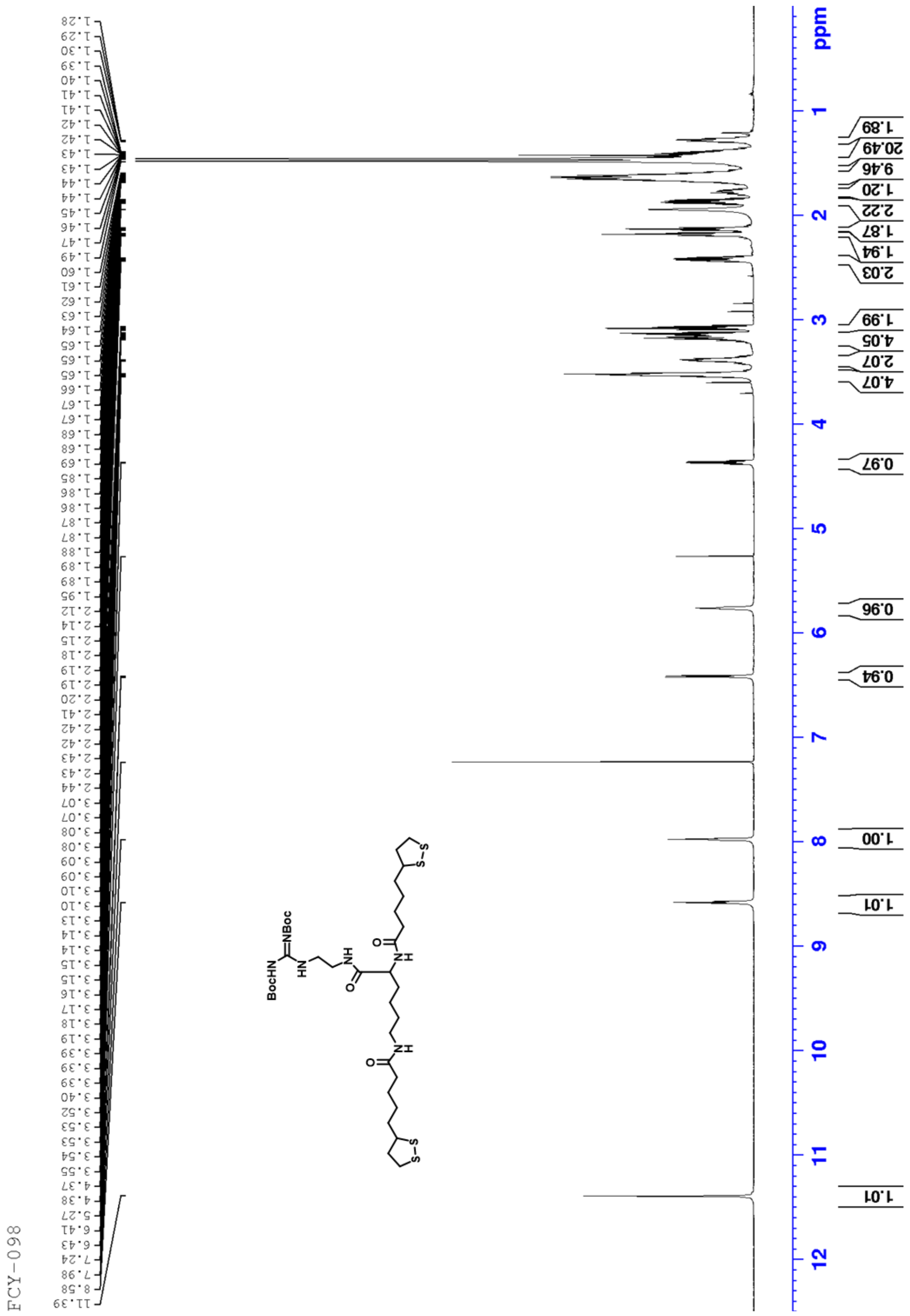

FCY-098
ppm
1 2 3 4 5 6 7 8 9 10 11 12
1.28 1.29 1.30 1.39 1.40 1.41 1.41 1.42 1.42 1.43 1.43 1.44 1.44 1.45 1.46 1.47 1.49 1.60 1.61 1.62 1.63 1.64 1.65 1.65 1.65 1.66 1.67 1.67 1.68 1.68 1.69 1.85 1.86 1.86 1.87 1.87 1.88 1.89 1.89 1.95 2.12 2.14 2.15 2.18 2.19 2.19 2.20 2.41 2.42 2.42 2.43 2.43 2.44 3.07 3.07 3.08 3.08 3.09 3.09 3.10 3.10 3.13 3.14 3.14 3.15 3.15 3.16 3.17 3.18 3.19 3.39 3.39 3.39 3.40 3.52 3.53 3.53 3.54 3.55 4.37 4.38 5.27 6.41 6.43 7.24 7.98 8.58 11.39
1.89 20.49 9.46 1.20 2.22 1.87 1.94 2.03 1.99 4.05 2.07 4.07 0.97 0.96 0.94 1.00 1.01 1.01
BocHN
NBoc
HN
NH
O
S-S

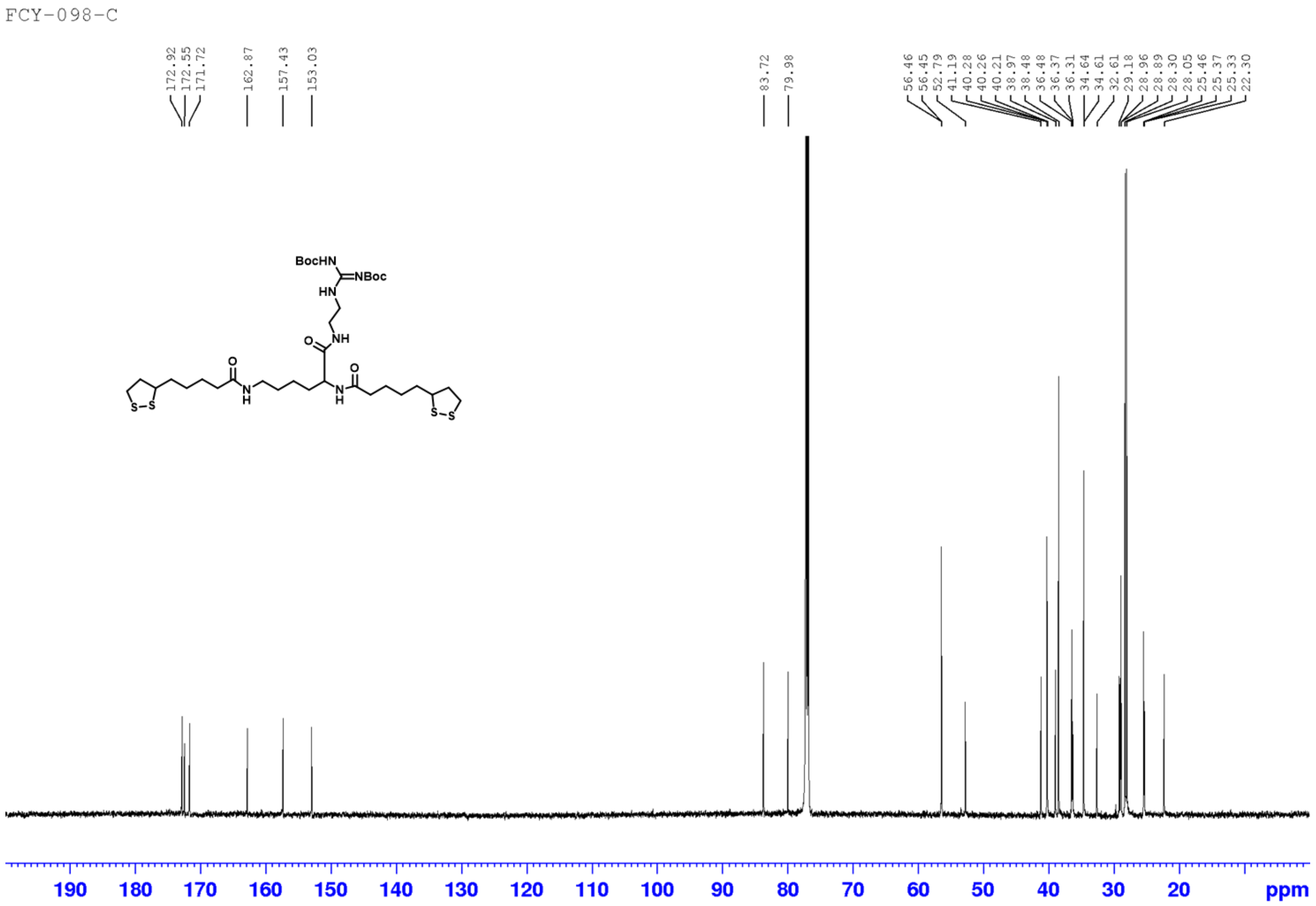
FCY-098-C
172.92
172.55
171.72
162.87
157.43
153.03
83.72
79.98
56.46
56.45
52.79
41.19
40.28
40.26
40.21
38.97
38.48
36.48
36.37
36.31
34.64
34.61
32.61
29.18
28.96
28.89
28.30
28.05
25.46
25.37
25.33
22.30
BocHN
NBoc
HN
NH
O
N
H
S-S
190
180
170
160
150
140
130
120
110
100
90
80
70
60
50
40
30
20
ppm

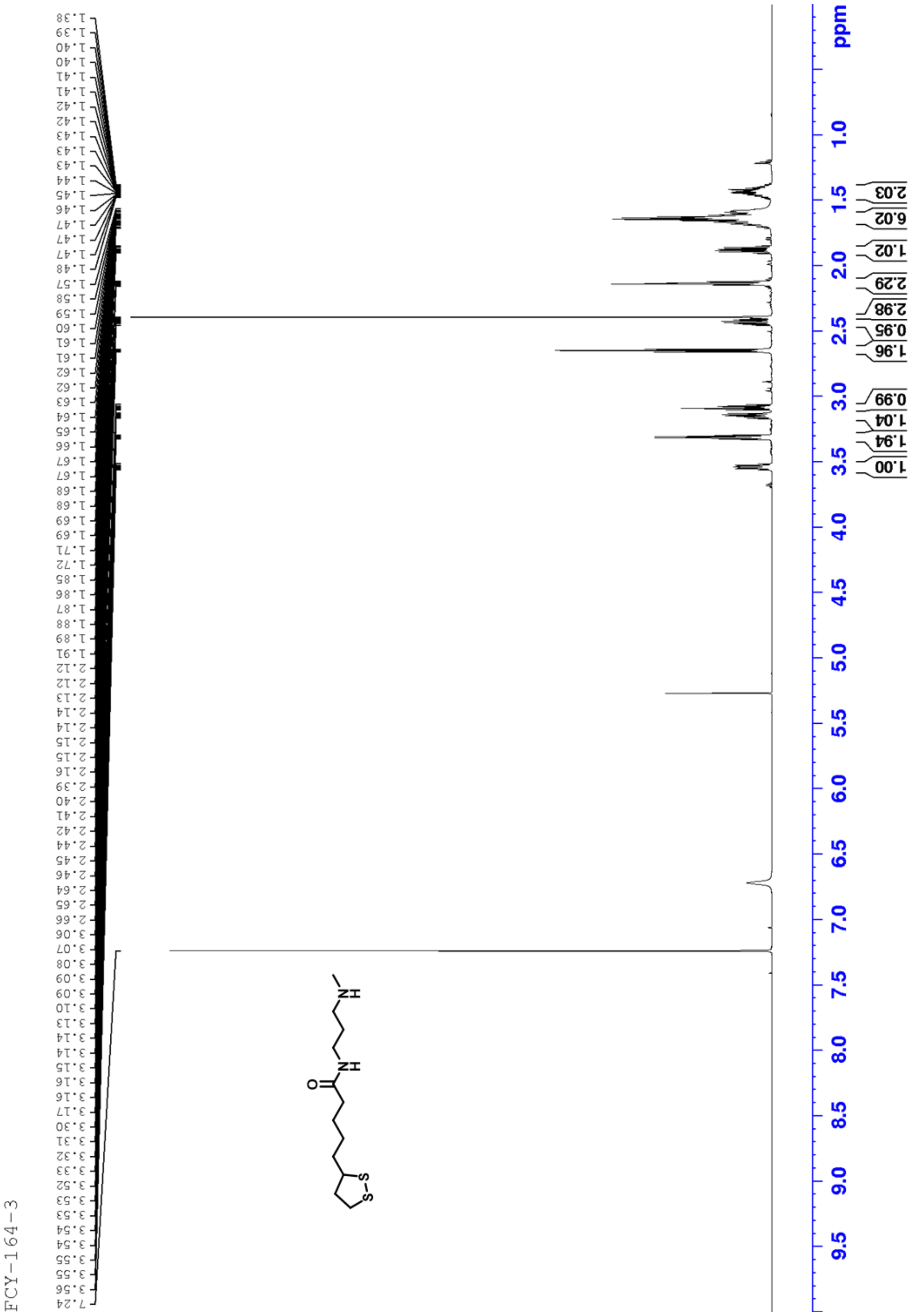
FCY-164-3
1.38 1.39 1.40 1.40 1.41 1.41 1.42 1.42 1.43 1.43 1.43 1.44 1.45 1.46 1.47 1.47 1.47 1.48 1.57 1.58 1.59 1.60 1.61 1.61 1.62 1.62 1.63 1.64 1.65 1.66 1.67 1.67 1.68 1.68 1.69 1.69 1.71 1.72 1.85 1.86 1.87 1.88 1.89 1.91 2.12 2.12 2.13 2.14 2.14 2.15 2.15 2.16 2.39 2.40 2.41 2.42 2.44 2.45 2.46 2.64 2.65 2.66 3.06 3.07 3.08 3.09 3.09 3.10 3.13 3.14 3.14 3.15 3.16 3.16 3.17 3.30 3.31 3.32 3.33 3.52 3.53 3.53 3.54 3.54 3.55 3.55 3.56 7.24
2.03
6.02
1.02
2.29
2.98
0.95
1.96
0.99
1.04
1.94
1.00
ppm
1.0 1.5 2.0 2.5 3.0 3.5 4.0 4.5 5.0 5.5 6.0 6.5 7.0 7.5 8.0 8.5 9.0 9.5

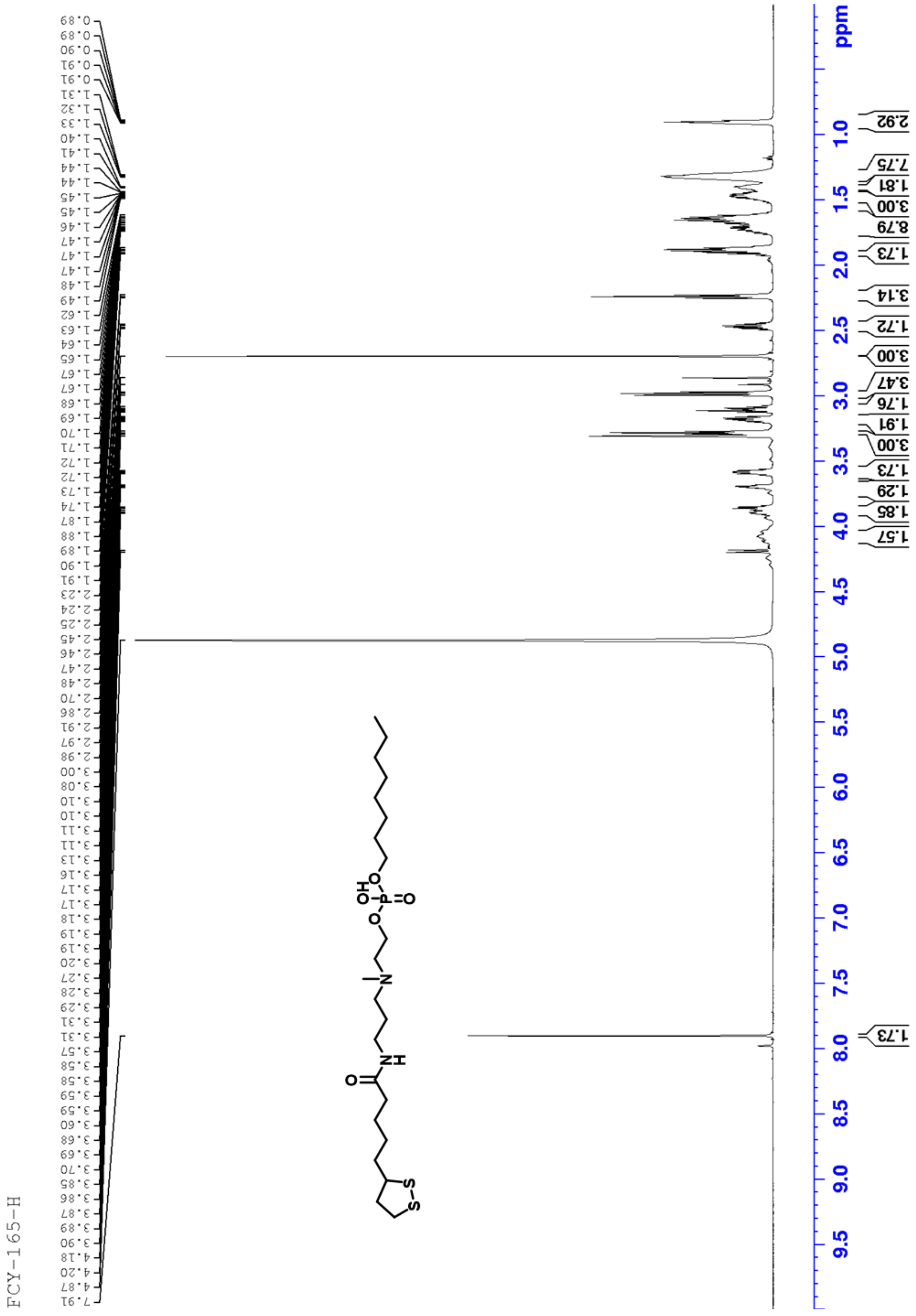
FCY-165-H
ppm

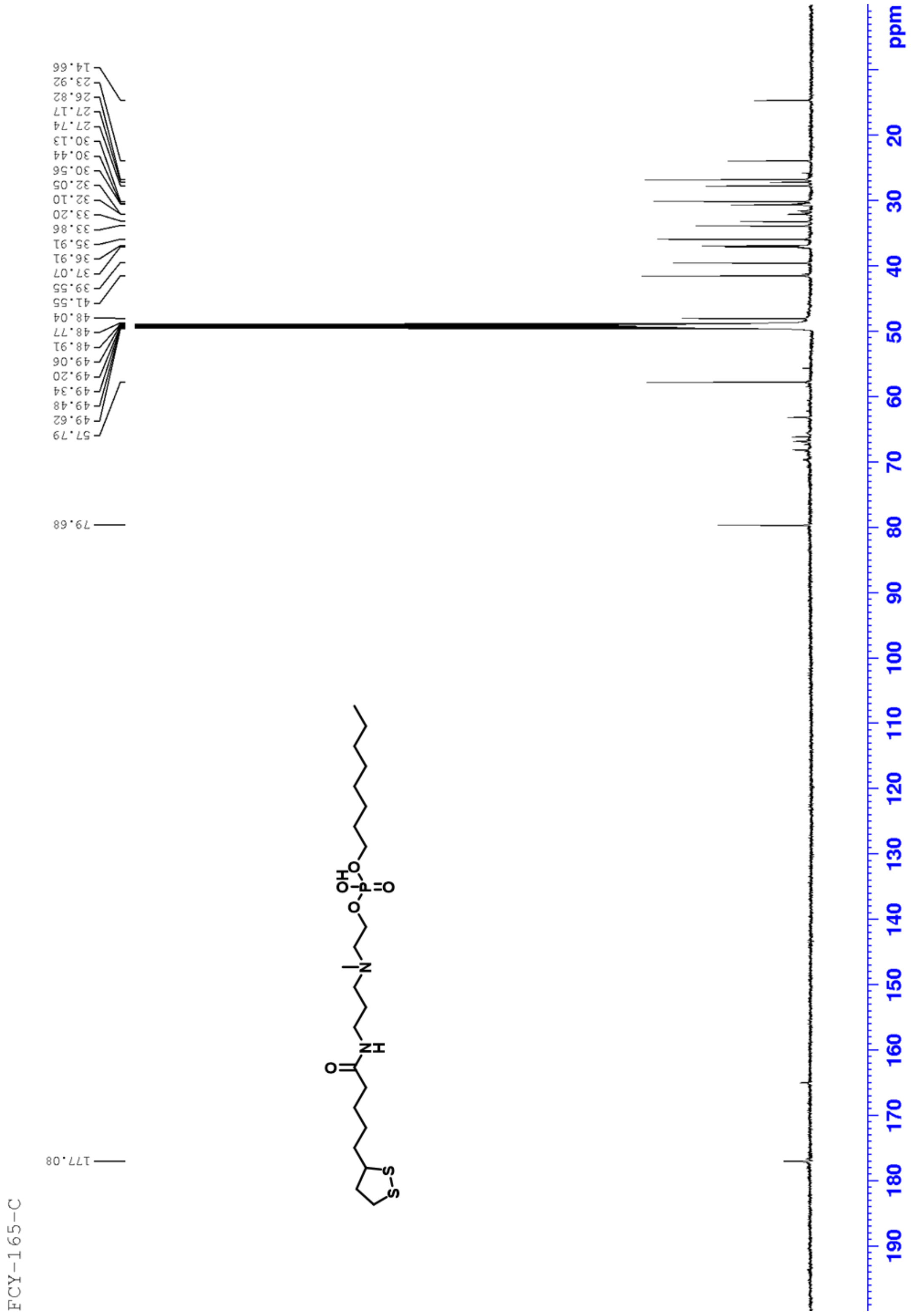
FCY-165-C
14.66
23.92
26.82
27.17
27.74
30.13
30.44
30.56
32.05
32.10
33.20
33.86
35.91
36.91
37.07
39.55
41.55
48.04
48.77
48.91
49.06
49.20
49.34
49.48
49.62
57.79
79.68
177.08
ppm
20
30
40
50
60
70
80
90
100
110
120
130
140
150
160
170
180
190

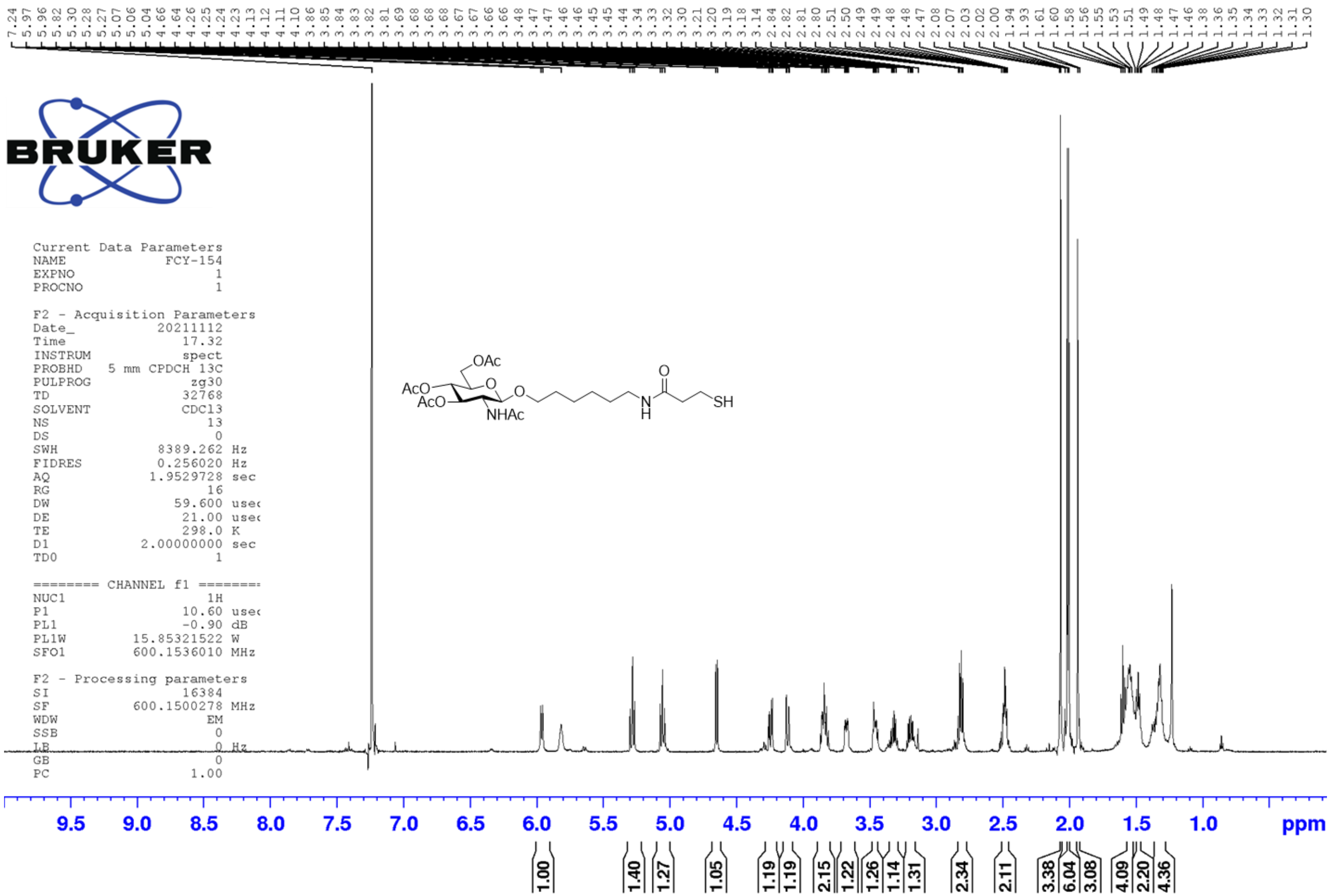
7.24 5.97 5.96 5.82 5.30 5.28 5.27 5.07 5.06 5.04 4.66 4.64 4.26 4.25 4.24 4.23 4.13 4.12 4.11 4.10 3.86 3.85 3.84 3.83 3.82 3.81 3.69 3.68 3.68 3.68 3.67 3.67 3.66 3.66 3.48 3.47 3.47 3.46 3.46 3.45 3.45 3.44 3.34 3.33 3.32 3.30 3.21 3.20 3.19 3.18 3.14 2.84 2.82 2.81 2.80 2.51 2.50 2.49 2.49 2.48 2.48 2.47 2.08 2.07 2.03 2.02 2.00 1.94 1.93 1.61 1.60 1.58 1.56 1.55 1.53 1.51 1.49 1.48 1.47 1.46 1.38 1.36 1.35 1.34 1.33 1.32 1.31 1.30
BRUKER
Current Data Parameters
NAME FCY-154
EXPNO 1
PROCNO 1
F2 - Acquisition Parameters
Date_ 20211112
Time 17.32
INSTRUM spect
PROBHD 5 mm CPDCH 13C
PULPROG zg30
TD 32768
SOLVENT CDCl3
NS 13
DS 0
SWH 8389.262 Hz
FIDRES 0.256020 Hz
AQ 1.9529728 sec
RG 16
DW 59.600 usec
DE 21.00 usec
TE 298.0 K
D1 2.00000000 sec
TD0 1
======== CHANNEL f1 ========
NUC1 1H
P1 10.60 usec
PL1 -0.90 dB
PL1W 15.85321522 W
SFO1 600.1536010 MHz
F2 - Processing parameters
SI 16384
SF 600.1500278 MHz
WDW EM
SSB 0
LB 0 Hz
GB 0
PC 1.00
OAc
AcO
AcO
O
NHAc
O
O
N
H
SH
9.5 9.0 8.5 8.0 7.5 7.0 6.5 6.0 5.5 5.0 4.5 4.0 3.5 3.0 2.5 2.0 1.5 1.0 ppm
1.00 1.40 1.27 1.05 1.19 1.19 2.15 1.22 1.26 1.14 1.31 2.34 2.11 3.38 6.04 3.08 4.09 2.20 4.36

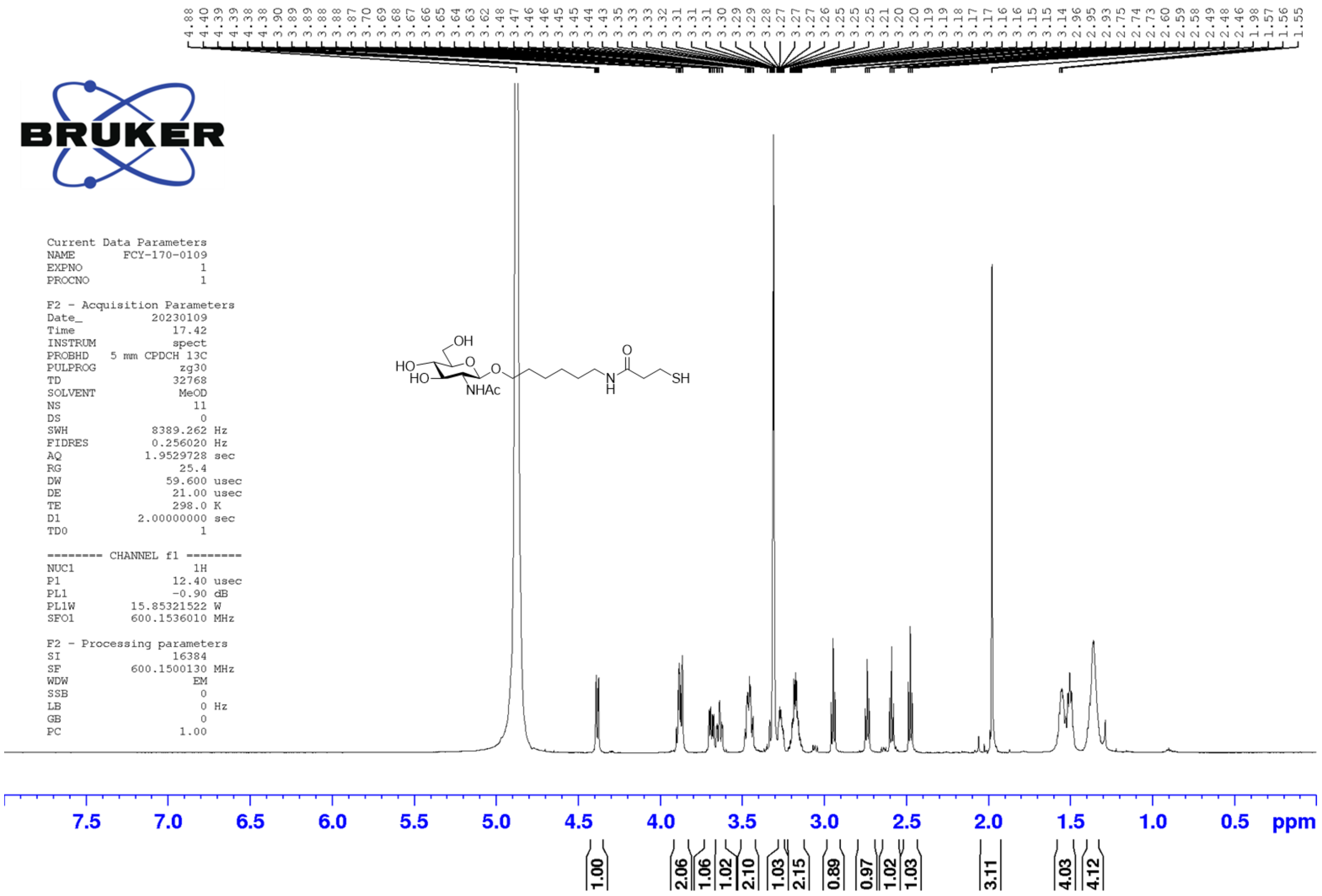
4.88 4.40 4.39 4.39 4.38 4.38 3.90 3.89 3.89 3.88 3.88 3.87 3.70 3.69 3.68 3.67 3.66 3.65 3.64 3.63 3.62 3.48 3.47 3.46 3.46 3.45 3.45 3.44 3.43 3.35 3.33 3.33 3.32 3.31 3.31 3.31 3.30 3.29 3.29 3.28 3.27 3.27 3.27 3.26 3.25 3.25 3.21 3.20 3.20 3.19 3.19 3.18 3.17 3.17 3.16 3.16 3.15 3.15 3.14 2.96 2.95 2.93 2.75 2.74 2.73 2.60 2.59 2.58 2.49 2.48 2.46 1.98 1.57 1.56 1.55
BRUKER
Current Data Parameters
NAME FCY-170-0109
EXPNO 1
PROCNO 1
F2 - Acquisition Parameters
Date_ 20230109
Time 17.42
INSTRUM spect
PROBHD 5 mm CPDCH 13C
PULPROG zg30
TD 32768
SOLVENT MeOD
NS 11
DS 0
SWH 8389.262 Hz
FIDRES 0.256020 Hz
AQ 1.9529728 sec
RG 25.4
DW 59.600 usec
DE 21.00 usec
TE 298.0 K
D1 2.00000000 sec
TD0 1
======== CHANNEL f1 ========
NUC1 1H
P1 12.40 usec
PL1 -0.90 dB
PL1W 15.85321522 W
SFO1 600.1536010 MHz
F2 - Processing parameters
SI 16384
SF 600.1500130 MHz
WDW EM
SSB 0
LB 0 Hz
GB 0
PC 1.00
OH
HO
HO
O
O
NHAc
N
H
O
SH
7.5 7.0 6.5 6.0 5.5 5.0 4.5 4.0 3.5 3.0 2.5 2.0 1.5 1.0 0.5 ppm
1.00 2.06 1.06 1.02 2.10 1.03 2.15 0.89 0.97 1.02 1.03 3.11 4.03 4.12

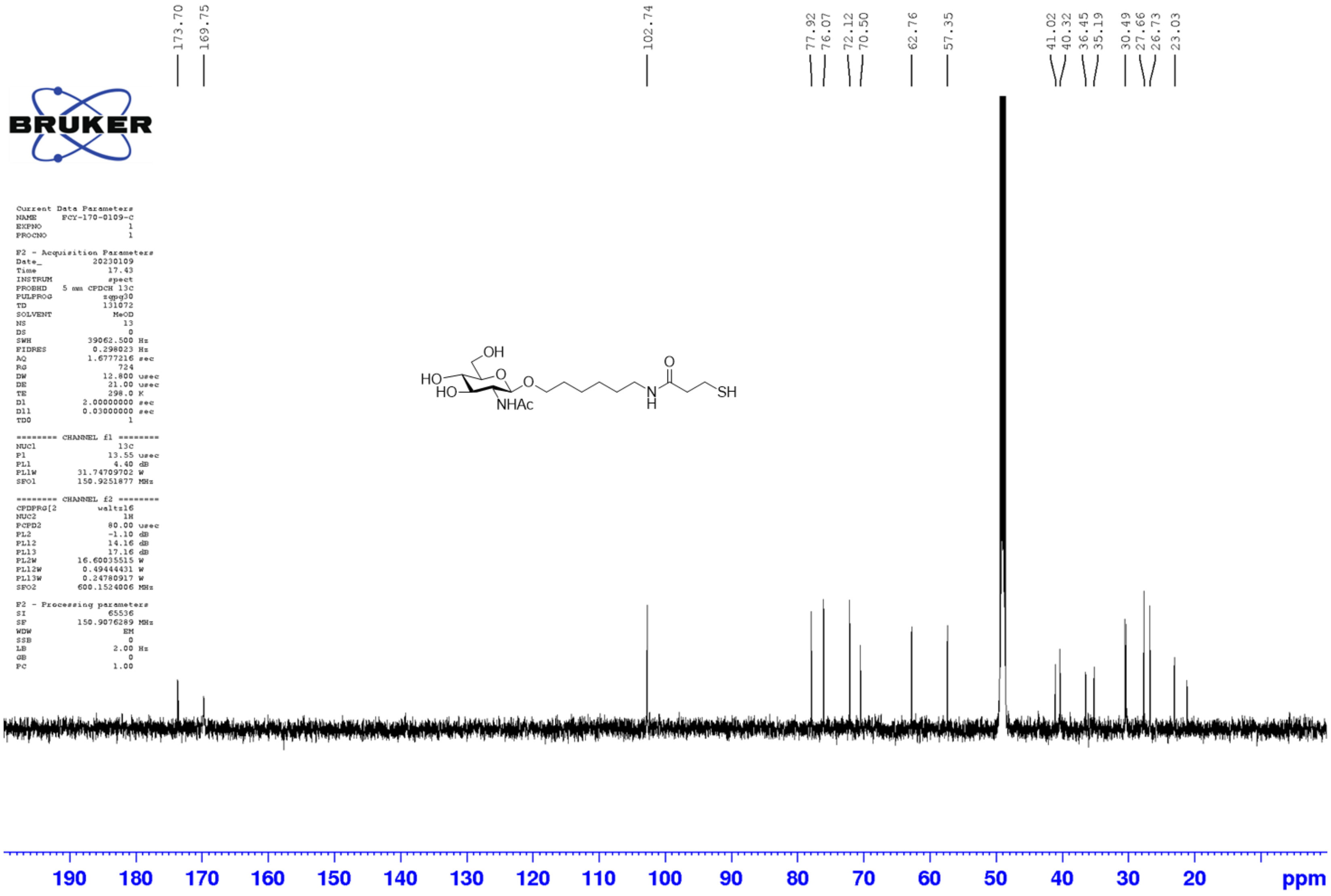
173.70
169.75
102.74
77.92
76.07
72.12
70.50
62.76
57.35
41.02
40.32
36.45
35.19
30.49
27.66
26.73
23.03
BRUKER
Current Data Parameters
NAME FCY-170-0109-c
EXPNO 1
PROCNO 1
F2 - Acquisition Parameters
Date_ 20230109
Time 17.43
INSTRUM spect
PROBHD 5 mm CPDCH 13C
PULPROG zgpg30
TD 131072
SOLVENT MeOD
NS 13
DS 0
SWH 39062.500 Hz
FIDRES 0.298023 Hz
AQ 1.6777216 sec
RG 724
DW 12.800 usec
DE 21.00 usec
TE 298.0 K
D1 2.00000000 sec
D11 0.03000000 sec
TD0 1
======== CHANNEL f1 ========
NUC1 13C
P1 13.55 usec
PL1 4.40 dB
PL1W 31.74709702 W
SFO1 150.9251877 MHz
======== CHANNEL f2 ========
CPDPRG[2 waltz16
NUC2 1H
PCPD2 80.00 usec
PL2 -1.10 dB
PL12 14.16 dB
PL13 17.16 dB
PL2W 16.60035515 W
PL12W 0.49444431 W
PL13W 0.24780917 W
SFO2 600.1524006 MHz
F2 - Processing parameters
SI 65536
SF 150.9076289 MHz
WDW EM
SSB 0
LB 2.00 Hz
GB 0
PC 1.00
OH
HO
HO
O
O
NHAc
O
N
H
SH
190
180
170
160
150
140
130
120
110
100
90
80
70
60
50
40
30
20
ppm

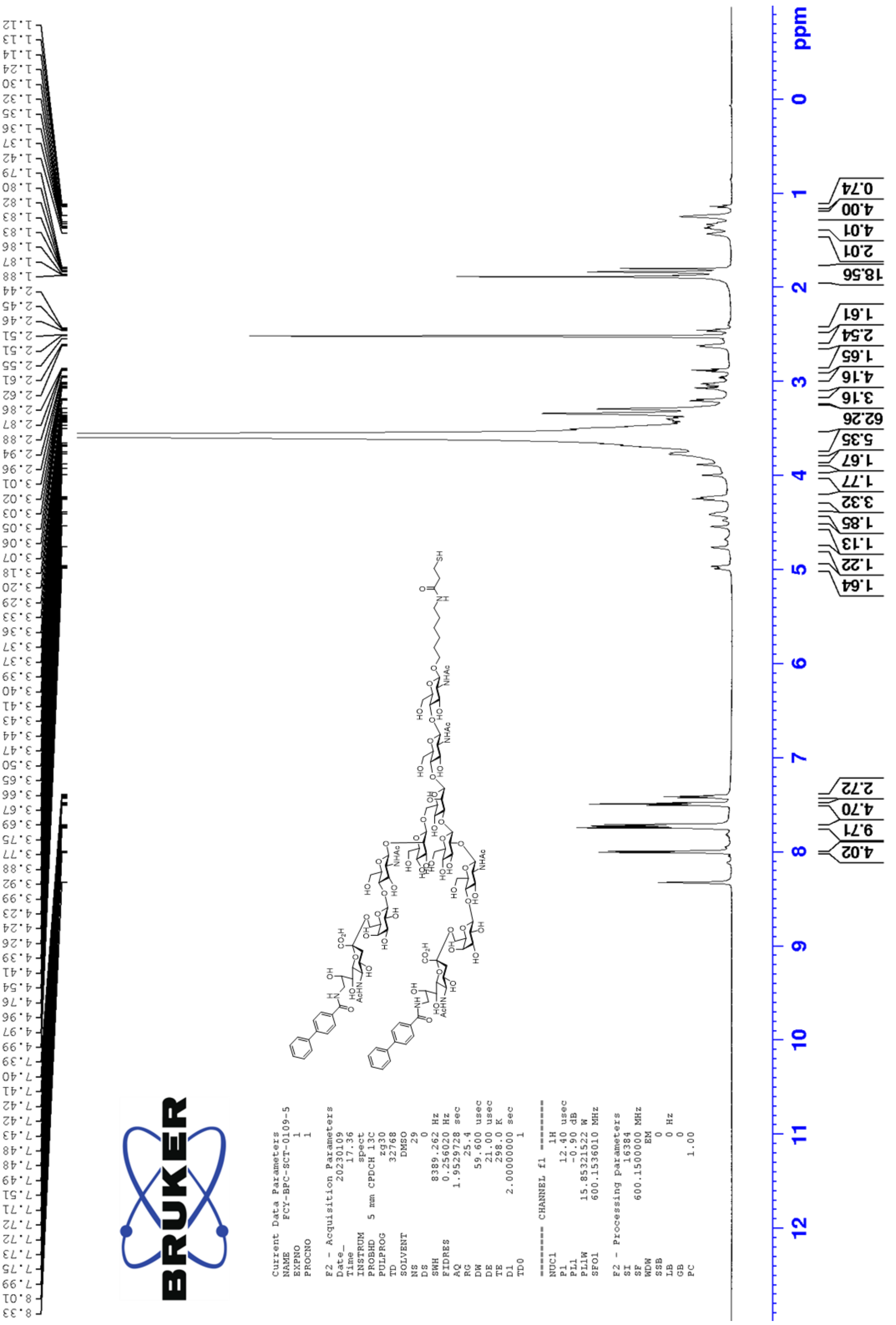

Current Data Parameters
NAME FCY-BPC-SCT-0109-5
EXPNO 1
PROCNO 1
F2 - Acquisition Parameters
Date_ 20230109
Time 17.36
INSTRUM spect
PROBHD 5 mm CPDCH 13C
PULPROG zg30
TD 32768
SOLVENT DMSO
NS 29
DS 0
SWH 8389.262 Hz
FIDRES 0.256020 Hz
AQ 1.9529728 sec
RG 25.4
DW 59.600 usec
DE 21.00 usec
TE 298.0 K
D1 2.00000000 sec
TD0 1
======== CHANNEL f1 ========
NUC1 1H
P1 12.40 usec
PL1 -0.90 dB
PL1W 15.85321522 W
SFO1 600.1536010 MHz
F2 - Processing parameters
SI 16384
SF 600.1500000 MHz
WDW EM
SSB 0
LB 0 Hz
GB 0
PC 1.00
ppm

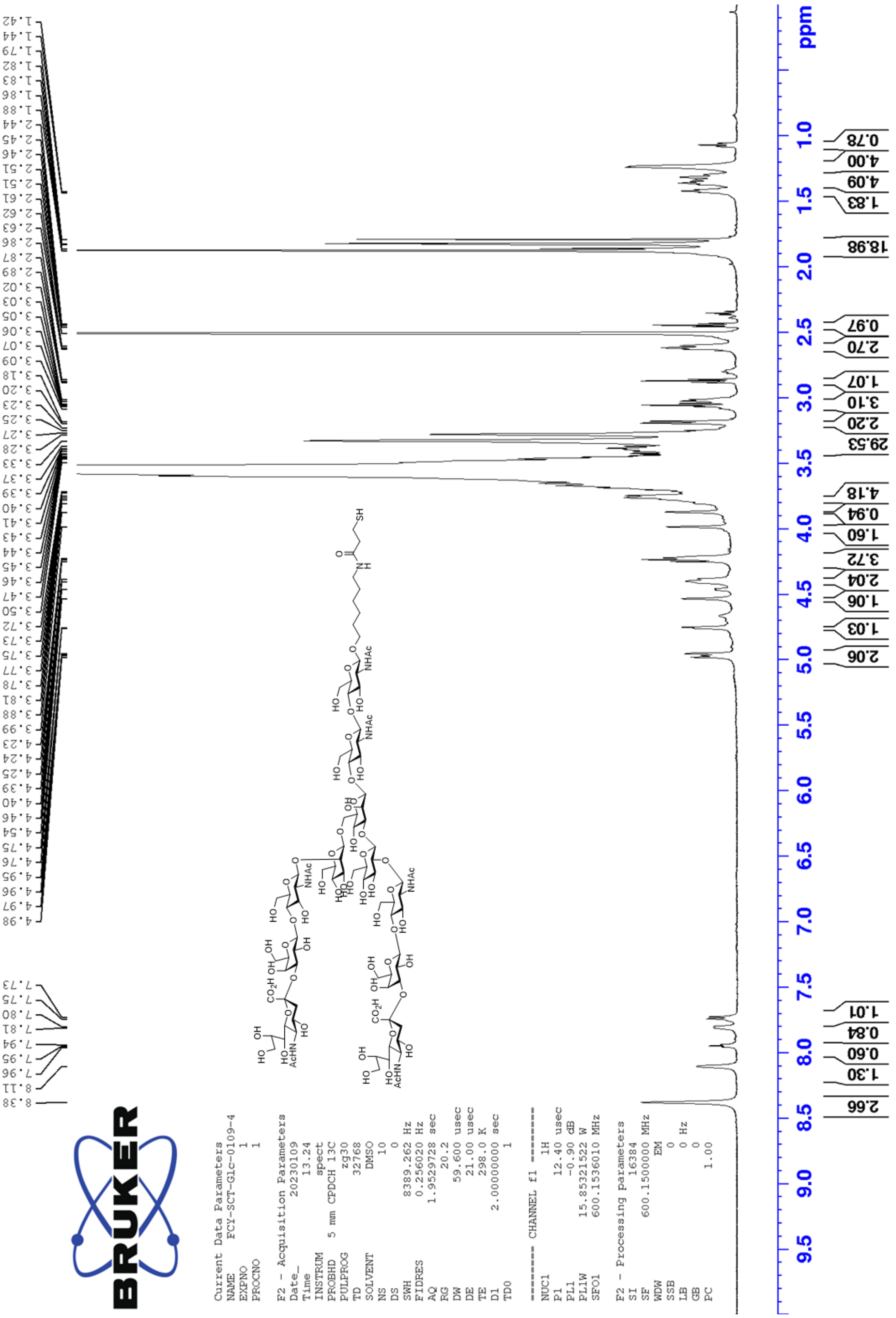

ppm
Current Data Parameters
NAME FCY-SCT-Glc-0109-4
EXPNO 1
PROCNO 1
F2 - Acquisition Parameters
Date_ 20230109
Time 13.24
INSTRUM spect
PROBHD 5 mm CPDCH 13C
PULPROG zg30
TD 32768
SOLVENT DMSO
NS 10
DS 0
SWH 8389.262 Hz
FIDRES 0.256020 Hz
AQ 1.9529728 sec
RG 20.2
DW 59.600 usec
DE 21.00 usec
TE 298.0 K
D1 2.00000000 sec
TD0 1
======== CHANNEL f1 ========
NUC1 1H
P1 12.40 usec
PL1 -0.90 dB
PL1W 15.85321522 W
SFO1 600.1536010 MHz
F2 - Processing parameters
SI 16384
SF 600.1500000 MHz
WDW EM
SSB 0
LB 0 Hz
GB 0
PC 1.00

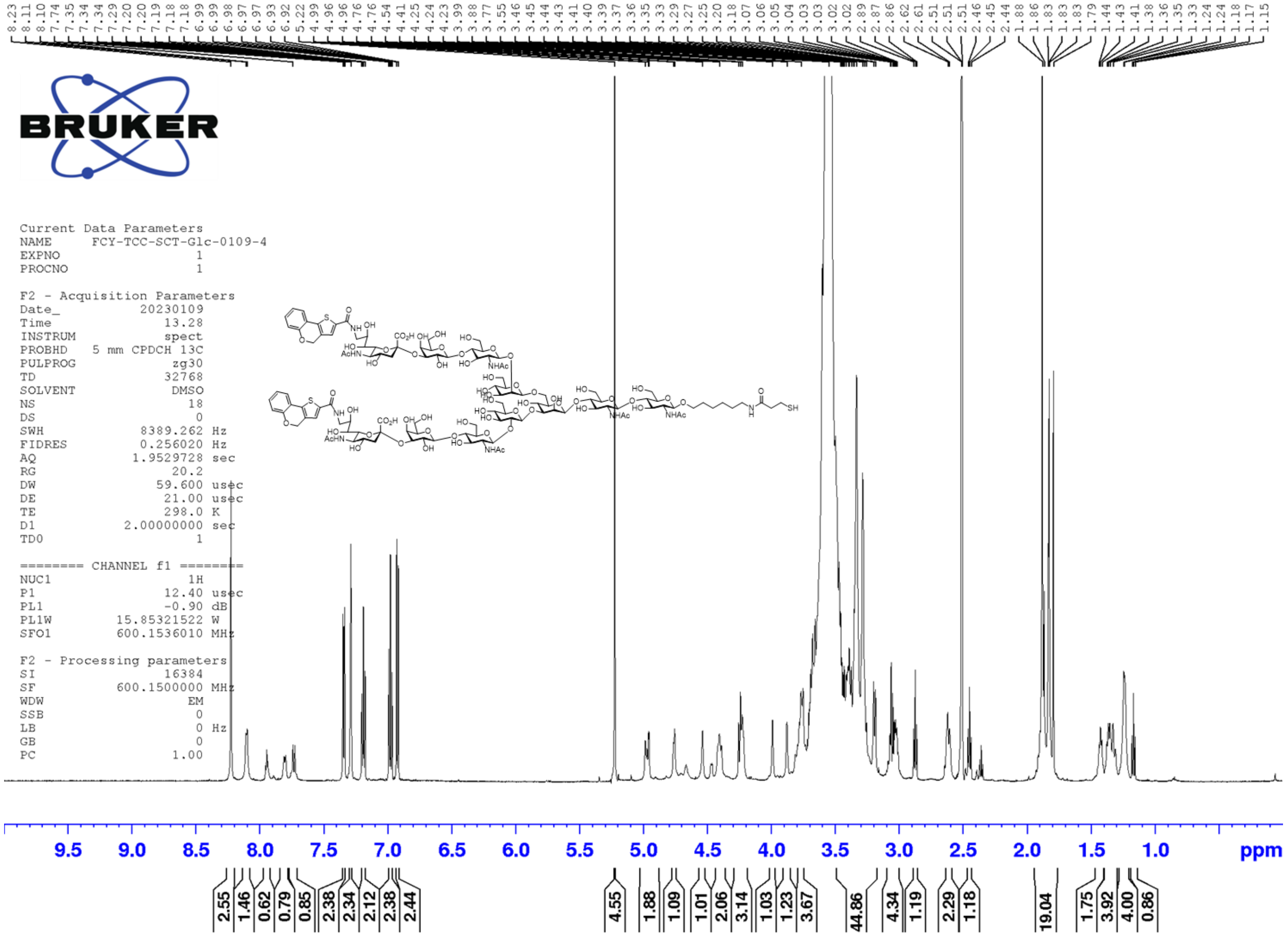
8.23 8.11 8.10 7.74 7.35 7.34 7.34 7.29 7.20 7.20 7.19 7.18 7.18 6.99 6.99 6.98 6.97 6.97 6.93 6.92 5.22 4.99 4.96 4.96 4.76 4.76 4.54 4.41 4.25 4.24 4.23 3.99 3.88 3.77 3.55 3.46 3.45 3.44 3.43 3.41 3.40 3.39 3.37 3.36 3.35 3.33 3.29 3.27 3.25 3.20 3.18 3.07 3.06 3.05 3.04 3.03 3.03 3.02 3.02 2.89 2.87 2.86 2.62 2.61 2.51 2.51 2.51 2.46 2.45 2.44 1.88 1.86 1.83 1.83 1.83 1.79 1.44 1.43 1.41 1.38 1.36 1.35 1.33 1.24 1.24 1.18 1.17 1.15
BRUKER
Current Data Parameters
NAME FCY-TCC-SCT-Glc-0109-4
EXPNO 1
PROCNO 1
F2 - Acquisition Parameters
Date_ 20230109
Time 13.28
INSTRUM spect
PROBHD 5 mm CPDCH 13C
PULPROG zg30
TD 32768
SOLVENT DMSO
NS 18
DS 0
SWH 8389.262 Hz
FIDRES 0.256020 Hz
AQ 1.9529728 sec
RG 20.2
DW 59.600 usec
DE 21.00 usec
TE 298.0 K
D1 2.00000000 sec
TD0 1
======== CHANNEL f1 ========
NUC1 1H
P1 12.40 usec
PL1 -0.90 dB
PL1W 15.85321522 W
SFO1 600.1536010 MHz
F2 - Processing parameters
SI 16384
SF 600.1500000 MHz
WDW EM
SSB 0
LB 0 Hz
GB 0
PC 1.00
9.5 9.0 8.5 8.0 7.5 7.0 6.5 6.0 5.5 5.0 4.5 4.0 3.5 3.0 2.5 2.0 1.5 1.0 ppm
2.55 1.46 0.62 0.79 0.85 2.38 2.34 2.12 2.38 2.44 4.55 1.88 1.09 1.01 2.06 3.14 1.03 1.23 3.67 44.86 4.34 1.19 2.29 1.18 19.04 1.75 3.92 4.00 0.86

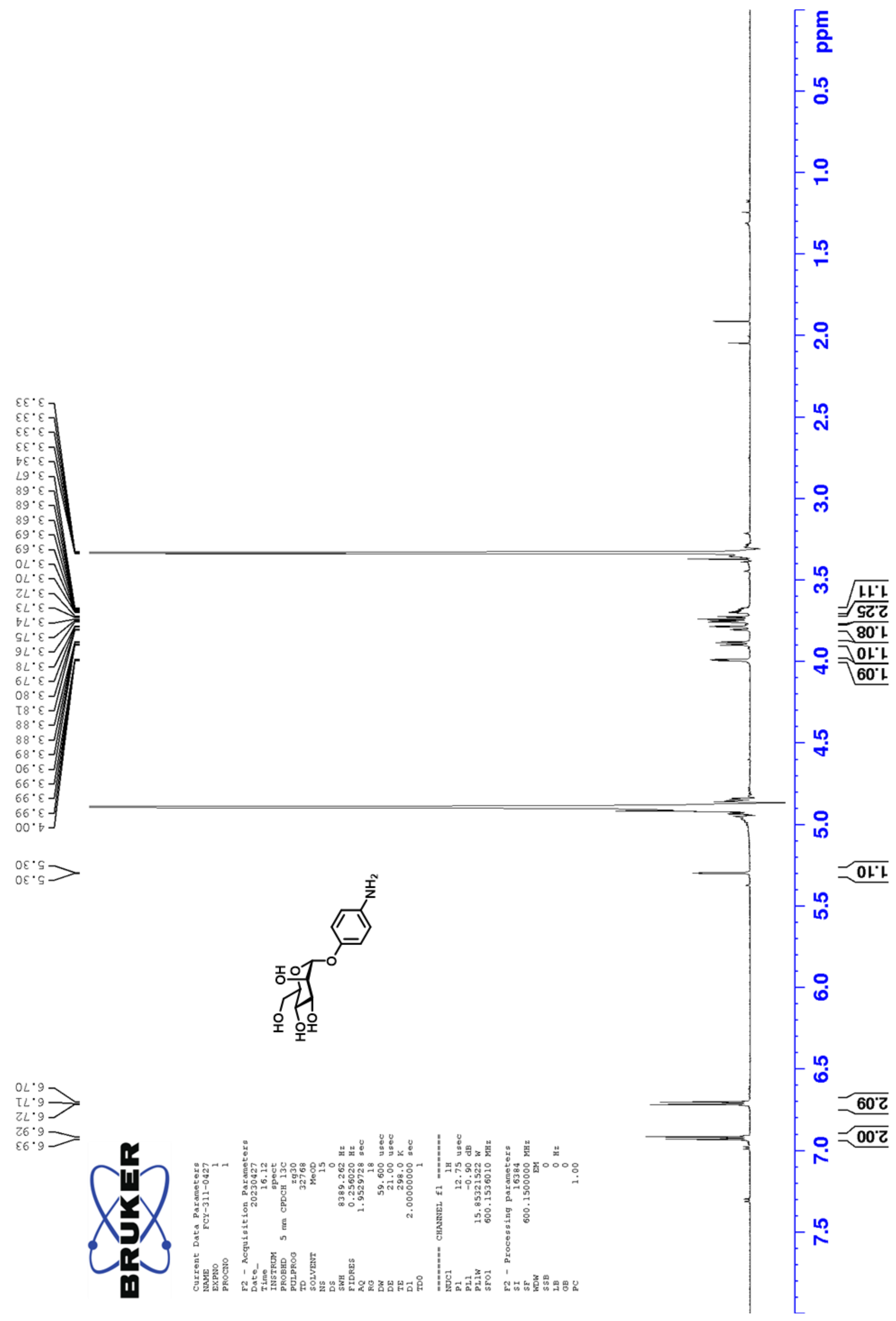
3.33
3.33
3.33
3.33
3.34
3.67
3.68
3.68
3.68
3.69
3.69
3.70
3.70
3.72
3.73
3.74
3.75
3.76
3.78
3.79
3.80
3.81
3.88
3.88
3.89
3.90
3.99
3.99
3.99
4.00
5.30
5.30
6.70
6.71
6.72
6.92
6.93
NH2
HO
HO
HO
OH
O
O
Current Data Parameters
NAME FCY-311-0427
EXPNO 1
PROCNO 1
F2 - Acquisition Parameters
Date_ 20230427
Time 16.12
INSTRUM spect
PROBHD 5 mm CPDCH 13C
PULPROG zg30
TD 32768
SOLVENT MeOD
NS 15
DS 0
SWH 8389.262 Hz
FIDRES 0.256020 Hz
AQ 1.9529728 sec
RG 18
DW 59.600 usec
DE 21.00 usec
TE 298.0 K
D1 2.00000000 sec
TD0 1
======== CHANNEL f1 ========
NUC1 1H
P1 12.75 usec
PL1 -0.90 dB
PL1W 15.85321522 W
SFO1 600.1536010 MHz
F2 - Processing parameters
SI 16384
SF 600.1500000 MHz
WDW EM
SSB 0
LB 0 Hz
GB 0
PC 1.00
1.11
2.25
1.08
1.10
1.09
1.10
2.09
2.00
ppm
0.5
1.0
1.5
2.0
2.5
3.0
3.5
4.0
4.5
5.0
5.5
6.0
6.5
7.0
7.5

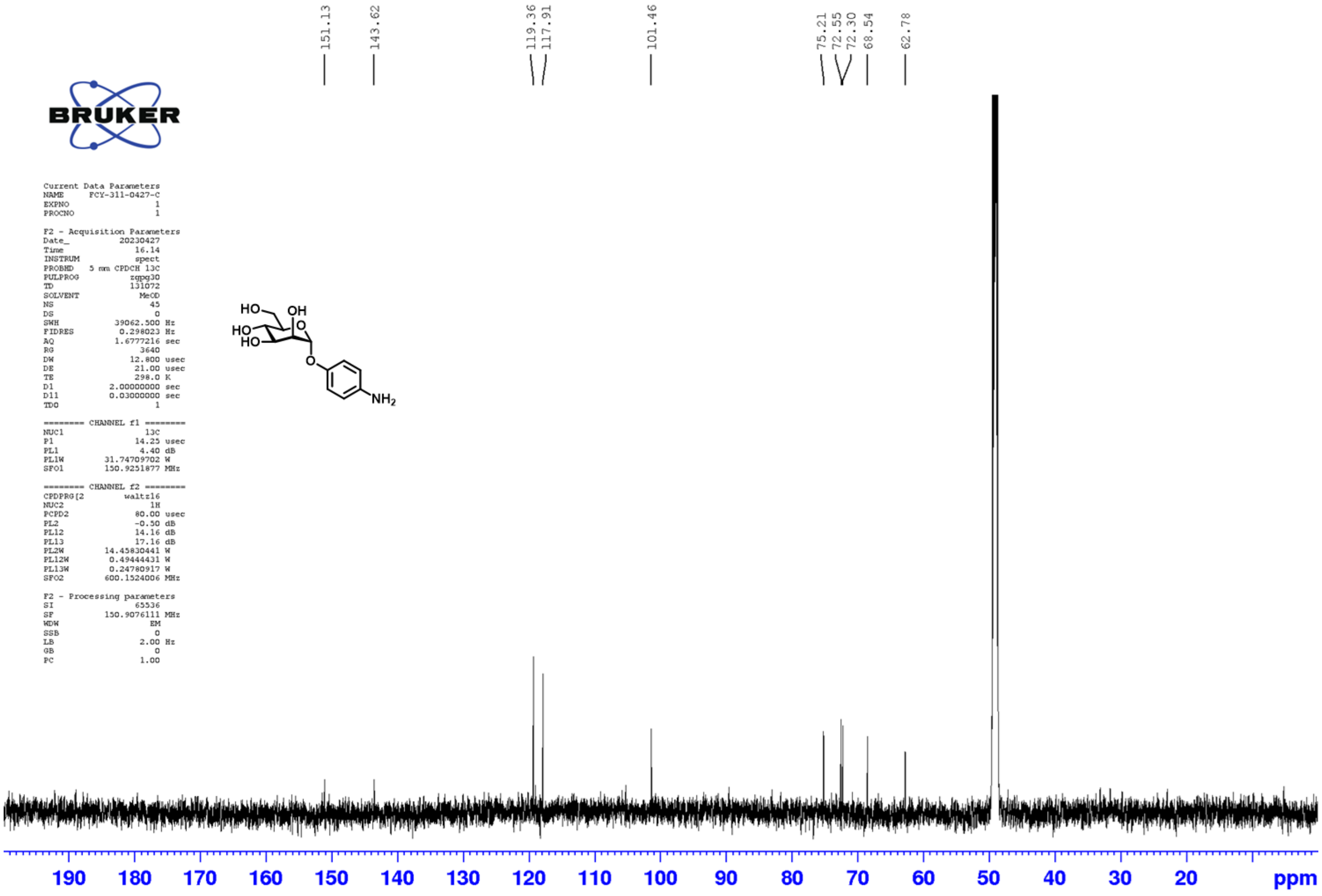
151.13
143.62
119.36
117.91
101.46
75.21
72.55
72.30
68.54
62.78
BRUKER
Current Data Parameters
NAME FCY-311-0427-C
EXPNO 1
PROCNO 1
F2 - Acquisition Parameters
Date_ 20230427
Time 16.14
INSTRUM spect
PROBHD 5 mm CPDCH 13C
PULPROG zgpg30
TD 131072
SOLVENT MeOD
NS 45
DS 0
SWH 39062.500 Hz
FIDRES 0.298023 Hz
AQ 1.6777216 sec
RG 3640
DW 12.800 usec
DE 21.00 usec
TE 298.0 K
D1 2.00000000 sec
D11 0.03000000 sec
TD0 1
======== CHANNEL f1 ========
NUC1 13C
P1 14.25 usec
PL1 4.40 dB
PL1W 31.74709702 W
SFO1 150.9251877 MHz
======== CHANNEL f2 ========
CPDPRG[2 waltz16
NUC2 1H
PCPD2 80.00 usec
PL2 -0.50 dB
PL12 14.16 dB
PL13 17.16 dB
PL2W 14.45830441 W
PL12W 0.49444431 W
PL13W 0.24780917 W
SFO2 600.1524006 MHz
F2 - Processing parameters
SI 65536
SF 150.9076111 MHz
WDW EM
SSB 0
LB 2.00 Hz
GB 0
PC 1.00
HO
OH
HO
HO
O
O
NH2
190
180
170
160
150
140
130
120
110
100
90
80
70
60
50
40
30
20
ppm

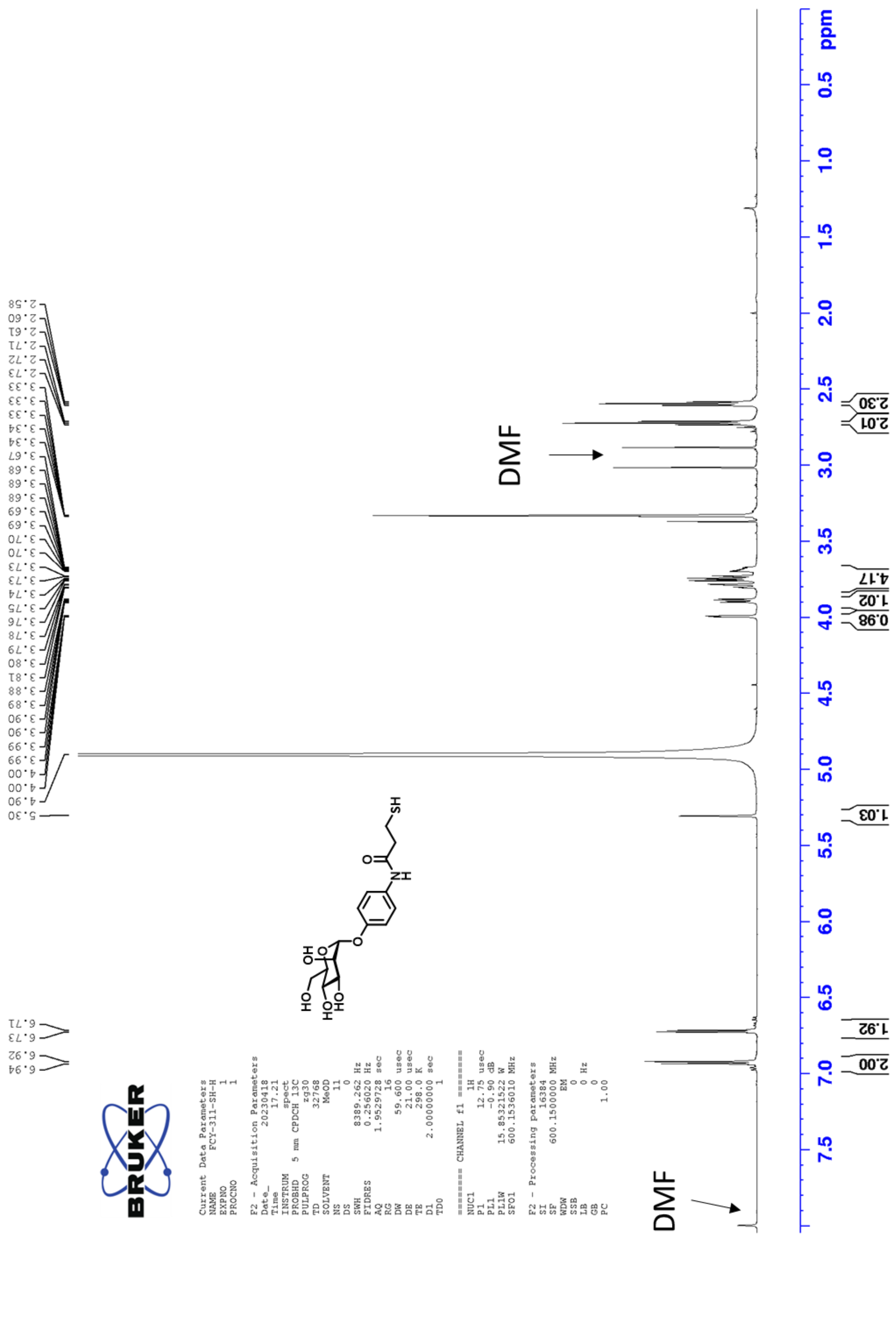
Current Data Parameters
NAME FCY-311-SH-H
EXPNO 1
PROCNO 1
F2 - Acquisition Parameters
Date_ 20230418
Time 17.21
INSTRUM spect
PROBHD 5 mm CPDCH 13C
PULPROG zg30
TD 32768
SOLVENT MeOD
NS 11
DS 0
SWH 8389.262 Hz
FIDRES 0.256020 Hz
AQ 1.9529728 sec
RG 16
DW 59.600 usec
DE 21.00 usec
TE 298.0 K
D1 2.00000000 sec
TD0 1
======== CHANNEL f1 ========
NUC1 1H
P1 12.75 usec
PL1 -0.90 dB
PL1W 15.85321522 W
SFO1 600.1536010 MHz
F2 - Processing parameters
SI 16384
SF 600.1500000 MHz
WDW EM
SSB 0
LB 0 Hz
GB 0
PC 1.00
DMF
DMF
ppm

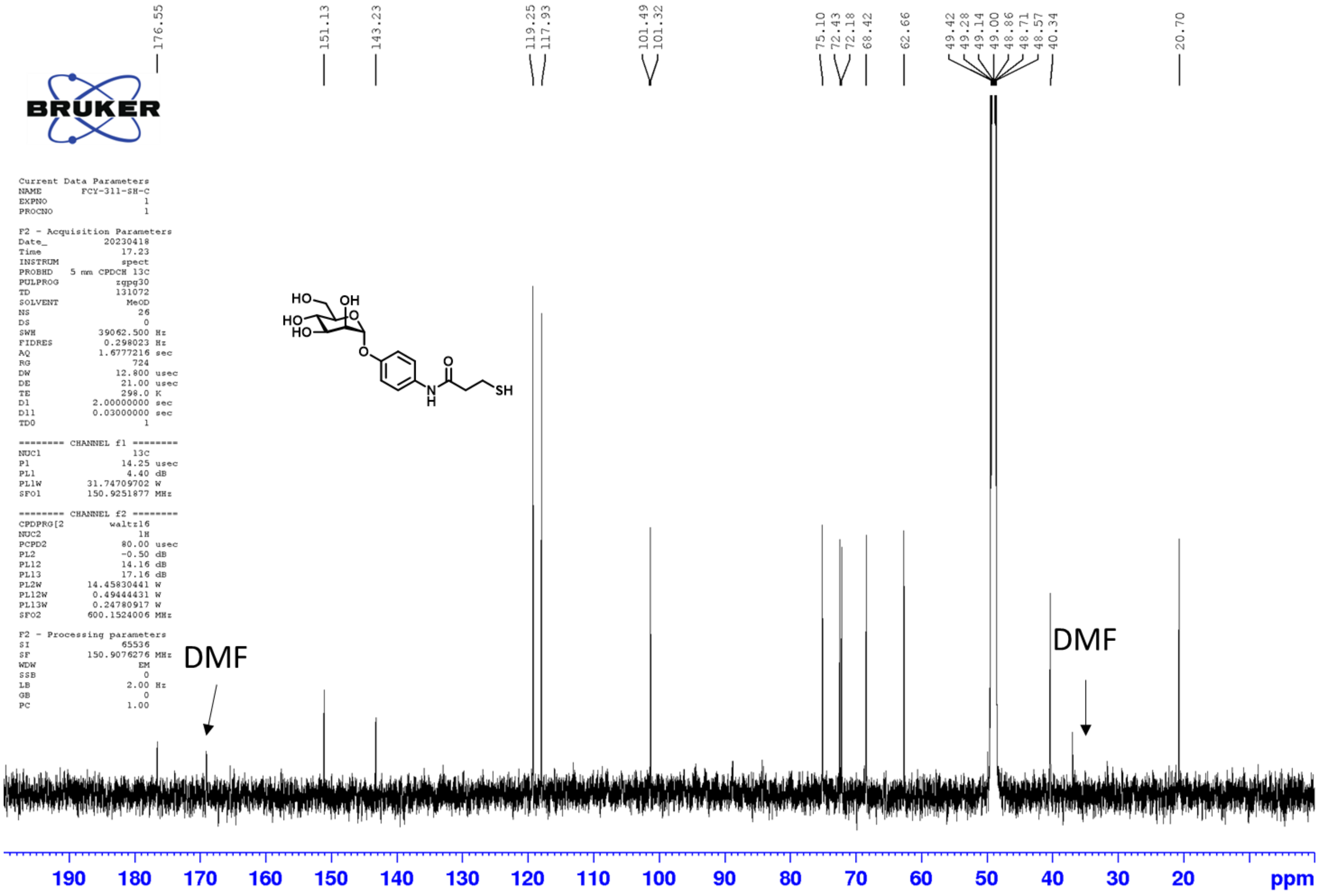
176.55
151.13
143.23
119.25
117.93
101.49
101.32
75.10
72.43
72.18
68.42
62.66
49.42
49.28
49.14
49.00
48.86
48.71
48.57
40.34
20.70
HO
OH
HO
HO
O
O
O
N
H
SH
Current Data Parameters
NAME FCY-311-SH-C
EXPNO 1
PROCNO 1
F2 - Acquisition Parameters
Date_ 20230418
Time 17.23
INSTRUM spect
PROBHD 5 mm CPDCH 13C
PULPROG zgpg30
TD 131072
SOLVENT MeOD
NS 26
DS 0
SWH 39062.500 Hz
FIDRES 0.298023 Hz
AQ 1.6777216 sec
RG 724
DW 12.800 usec
DE 21.00 usec
TE 298.0 K
D1 2.00000000 sec
D11 0.03000000 sec
TD0 1
======== CHANNEL f1 ========
NUC1 13C
P1 14.25 usec
PL1 4.40 dB
PL1W 31.74709702 W
SFO1 150.9251877 MHz
======== CHANNEL f2 ========
CPDPRG[2 waltz16
NUC2 1H
PCPD2 80.00 usec
PL2 -0.50 dB
PL12 14.16 dB
PL13 17.16 dB
PL2W 14.45830441 W
PL12W 0.49444431 W
PL13W 0.24780917 W
SFO2 600.1524006 MHz
F2 - Processing parameters
SI 65536
SF 150.9076276 MHz
WDW EM
SSB 0
LB 2.00 Hz
GB 0
PC 1.00
DMF
DMF
190
180
170
160
150
140
130
120
110
100
90
80
70
60
50
40
30
20
ppm

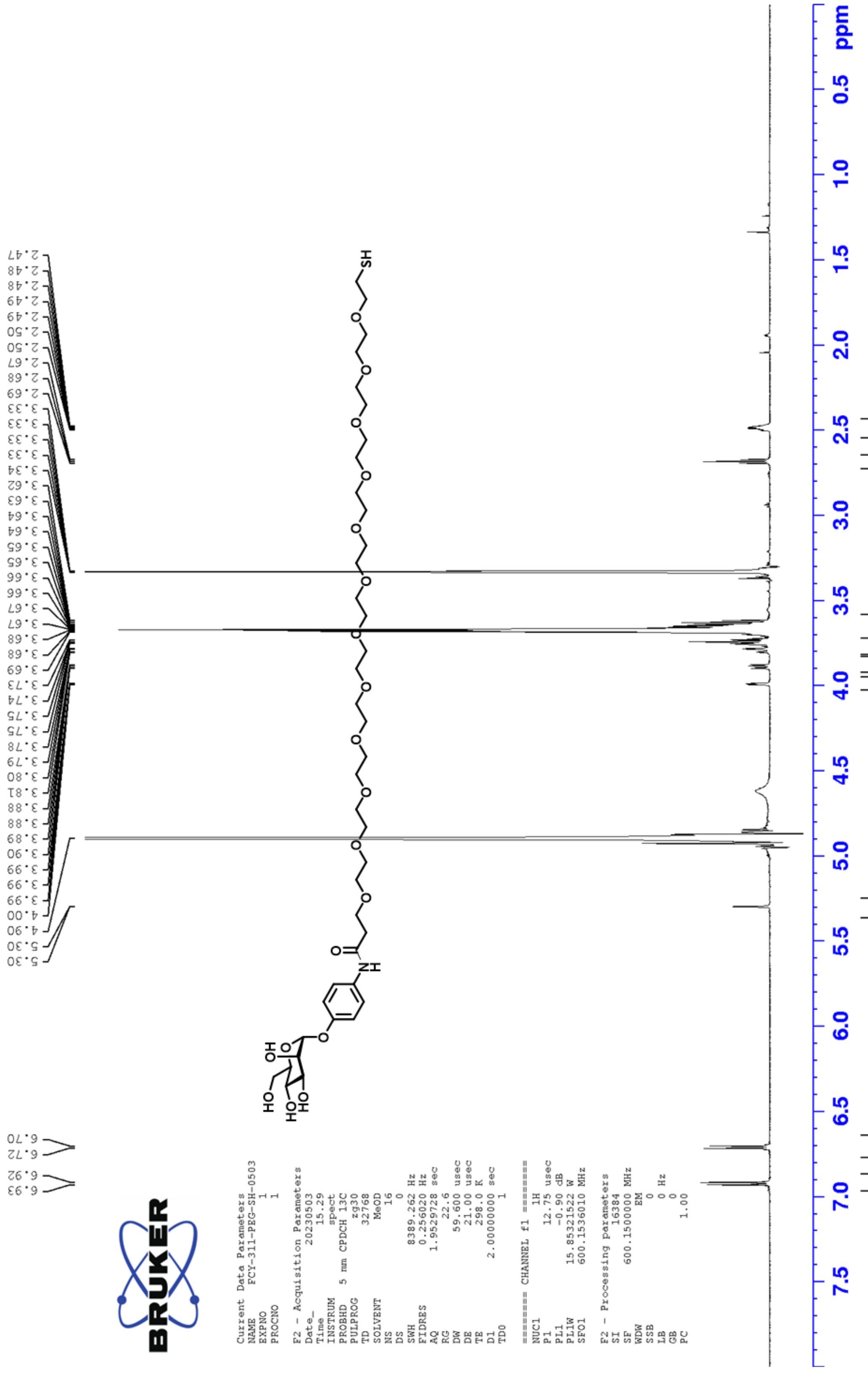

6.93
6.92
6.72
6.70
5.30
5.30
4.90
4.00
3.99
3.99
3.99
3.90
3.89
3.88
3.88
3.81
3.80
3.79
3.78
3.75
3.75
3.74
3.73
3.69
3.68
3.68
3.67
3.67
3.66
3.66
3.65
3.65
3.64
3.64
3.63
3.62
3.34
3.33
3.33
3.33
3.33
2.69
2.68
2.67
2.50
2.50
2.49
2.49
2.48
2.48
2.47
BRUKER
Current Data Parameters
NAME FCY-311-PEG-SH-0503
EXPNO 1
PROCNO 1
F2 - Acquisition Parameters
Date_ 20230503
Time 15.29
INSTRUM spect
PROBHD 5 mm CPDCH 13C
PULPROG zg30
TD 32768
SOLVENT MeOD
NS 16
DS 0
SWH 8389.262 Hz
FIDRES 0.256020 Hz
AQ 1.9529728 sec
RG 22.6
DW 59.600 usec
DE 21.00 usec
TE 298.0 K
D1 2.00000000 sec
TD0 1
======== CHANNEL f1 ========
NUC1 1H
P1 12.75 usec
PL1 -0.90 dB
PL1W 15.85321522 W
SFO1 600.1536010 MHz
F2 - Processing parameters
SI 16384
SF 600.1500000 MHz
WDW EM
SSB 0
LB 0 Hz
GB 0
PC 1.00
HO
HO
HO
OH
O
O
N
H
O
O
O
O
O
O
O
O
O
O
O
O
SH
2.00
1.99
1.02
1.01
1.09
5.54
48.64
1.74
2.02
7.5
7.0
6.5
6.0
5.5
5.0
4.5
4.0
3.5
3.0
2.5
2.0
1.5
1.0
0.5
ppm

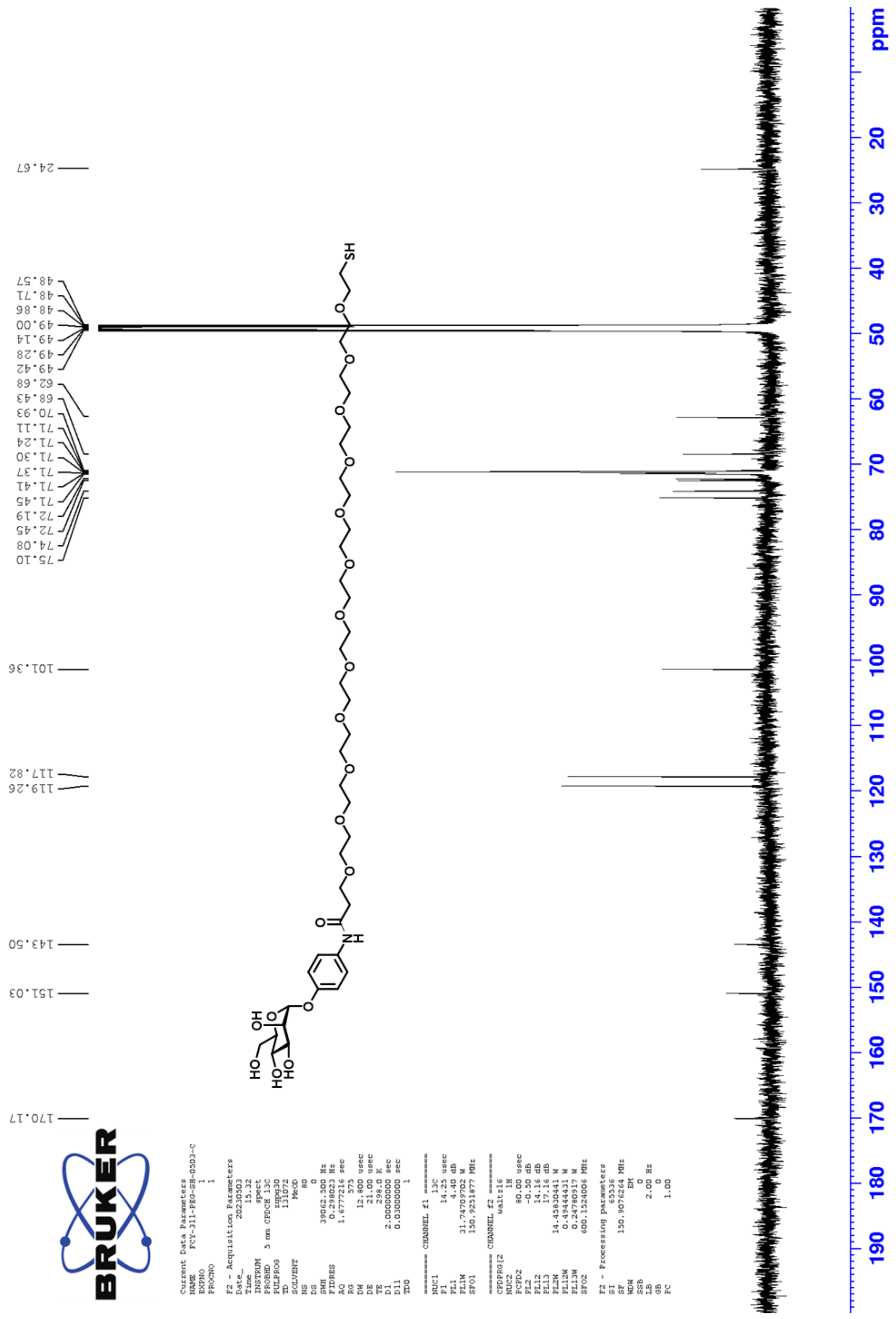
24.67
48.57
48.71
48.86
49.00
49.14
49.28
49.42
62.68
68.43
70.93
71.11
71.24
71.30
71.37
71.41
71.45
72.19
72.45
74.08
75.10
101.36
117.82
119.26
143.50
151.03
170.17
ppm

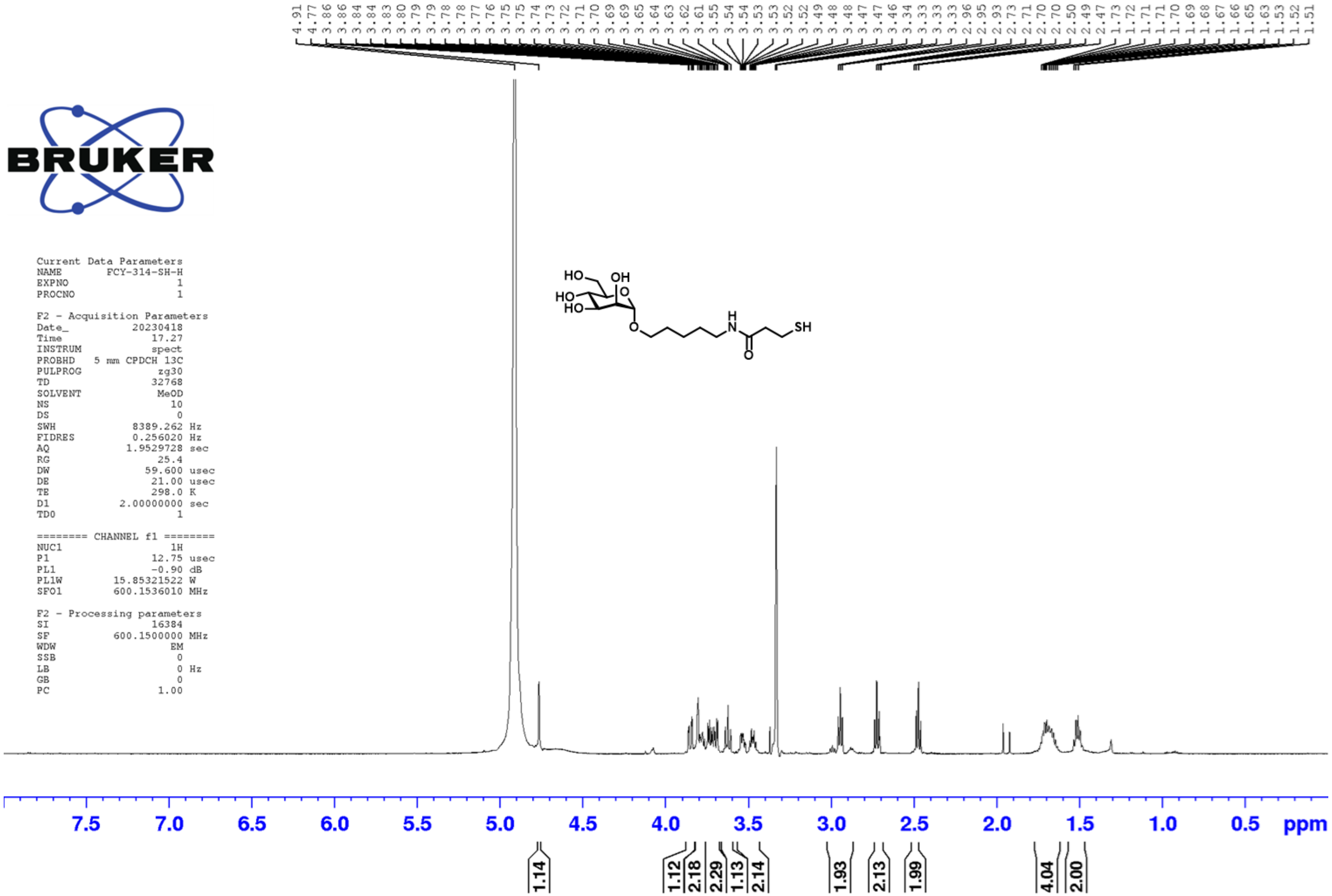
4.91 4.77 3.86 3.86 3.84 3.84 3.83 3.80 3.79 3.79 3.78 3.78 3.77 3.76 3.75 3.75 3.74 3.73 3.72 3.71 3.70 3.69 3.69 3.65 3.64 3.63 3.62 3.61 3.55 3.54 3.54 3.53 3.53 3.52 3.49 3.48 3.48 3.47 3.47 3.46 3.34 3.33 3.33 3.33 2.96 2.95 2.93 2.73 2.71 2.70 2.70 2.50 2.49 2.47 1.73 1.72 1.71 1.71 1.70 1.69 1.68 1.67 1.66 1.65 1.63 1.53 1.52 1.51
BRUKER
Current Data Parameters
NAME FCY-314-SH-H
EXPNO 1
PROCNO 1
F2 - Acquisition Parameters
Date_ 20230418
Time 17.27
INSTRUM spect
PROBHD 5 mm CPDCH 13C
PULPROG zg30
TD 32768
SOLVENT MeOD
NS 10
DS 0
SWH 8389.262 Hz
FIDRES 0.256020 Hz
AQ 1.9529728 sec
RG 25.4
DW 59.600 usec
DE 21.00 usec
TE 298.0 K
D1 2.00000000 sec
TD0 1
======== CHANNEL f1 ========
NUC1 1H
P1 12.75 usec
PL1 -0.90 dB
PL1W 15.85321522 W
SFO1 600.1536010 MHz
F2 - Processing parameters
SI 16384
SF 600.1500000 MHz
WDW EM
SSB 0
LB 0 Hz
GB 0
PC 1.00
HO
OH
O
HO
HO
O
H
N
O
SH
7.5 7.0 6.5 6.0 5.5 5.0 4.5 4.0 3.5 3.0 2.5 2.0 1.5 1.0 0.5 ppm
1.14
1.12 2.18 2.29 1.13 2.14
1.93 2.13 1.99
4.04 2.00

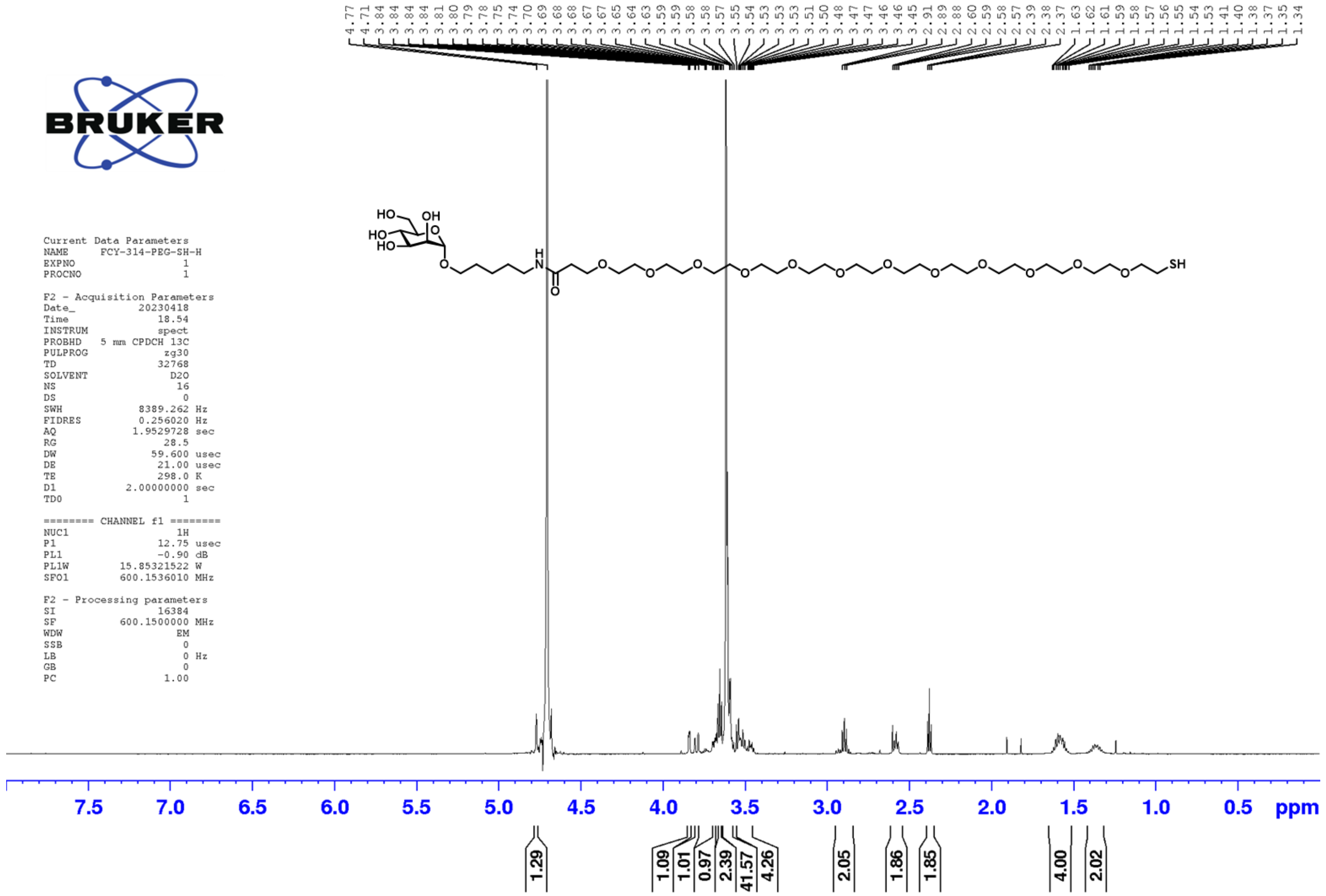
4.77
4.71
3.84
3.84
3.84
3.84
3.81
3.80
3.79
3.78
3.75
3.74
3.70
3.69
3.68
3.68
3.67
3.67
3.65
3.64
3.63
3.59
3.59
3.58
3.58
3.57
3.55
3.54
3.53
3.53
3.53
3.51
3.50
3.48
3.47
3.47
3.46
3.46
3.45
2.91
2.89
2.88
2.60
2.59
2.58
2.57
2.39
2.38
2.37
1.63
1.62
1.61
1.59
1.58
1.57
1.56
1.55
1.54
1.53
1.41
1.40
1.38
1.37
1.35
1.34
BRUKER
Current Data Parameters
NAME FCY-314-PEG-SH-H
EXPNO 1
PROCNO 1
F2 - Acquisition Parameters
Date_ 20230418
Time 18.54
INSTRUM spect
PROBHD 5 mm CPDCH 13C
PULPROG zg30
TD 32768
SOLVENT D2O
NS 16
DS 0
SWH 8389.262 Hz
FIDRES 0.256020 Hz
AQ 1.9529728 sec
RG 28.5
DW 59.600 usec
DE 21.00 usec
TE 298.0 K
D1 2.00000000 sec
TD0 1
======== CHANNEL f1 ========
NUC1 1H
P1 12.75 usec
PL1 -0.90 dB
PL1W 15.85321522 W
SFO1 600.1536010 MHz
F2 - Processing parameters
SI 16384
SF 600.1500000 MHz
WDW EM
SSB 0
LB 0 Hz
GB 0
PC 1.00
HO
OH
HO
HO
O
O
H
N
O
O
O
O
O
O
O
O
O
O
O
O
O
SH
7.5
7.0
6.5
6.0
5.5
5.0
4.5
4.0
3.5
3.0
2.5
2.0
1.5
1.0
0.5
ppm
1.29
1.09
1.01
0.97
2.39
41.57
4.26
2.05
1.86
1.85
4.00
2.02

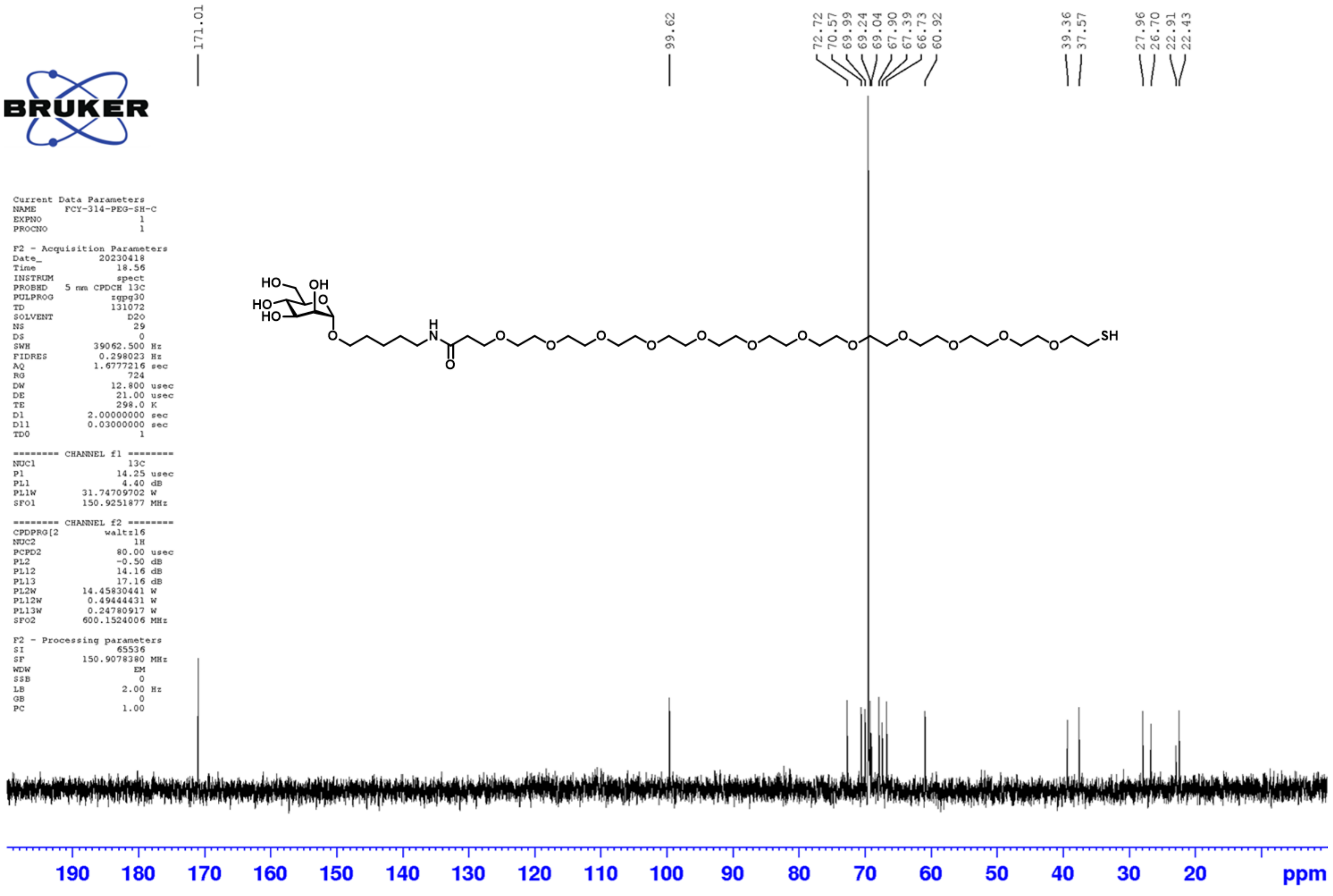

171.01
99.62
72.72
70.57
69.99
69.24
69.04
67.90
67.39
66.73
60.92
39.36
37.57
27.96
26.70
22.91
22.43
BRUKER
Current Data Parameters
NAME FCY-314-PEG-SH-C
EXPNO 1
PROCNO 1
F2 - Acquisition Parameters
Date_ 20230418
Time 18.56
INSTRUM spect
PROBHD 5 mm CPDCH 13C
PULPROG zgpg30
TD 131072
SOLVENT D2O
NS 29
DS 0
SWH 39062.500 Hz
FIDRES 0.298023 Hz
AQ 1.6777216 sec
RG 724
DW 12.800 usec
DE 21.00 usec
TE 298.0 K
D1 2.00000000 sec
D11 0.03000000 sec
TD0 1
======== CHANNEL f1 ========
NUC1 13C
P1 14.25 usec
PL1 4.40 dB
PL1W 31.74709702 W
SFO1 150.9251877 MHz
======== CHANNEL f2 ========
CPDPRG[2 waltz16
NUC2 1H
PCPD2 80.00 usec
PL2 -0.50 dB
PL12 14.16 dB
PL13 17.16 dB
PL2W 14.45830441 W
PL12W 0.49444431 W
PL13W 0.24780917 W
SFO2 600.1524006 MHz
F2 - Processing parameters
SI 65536
SF 150.9078380 MHz
WDW EM
SSB 0
LB 2.00 Hz
GB 0
PC 1.00
HO
OH
HO
HO
O
O
H
N
O
O
SH
190 180 170 160 150 140 130 120 110 100 90 80 70 60 50 40 30 20 ppm

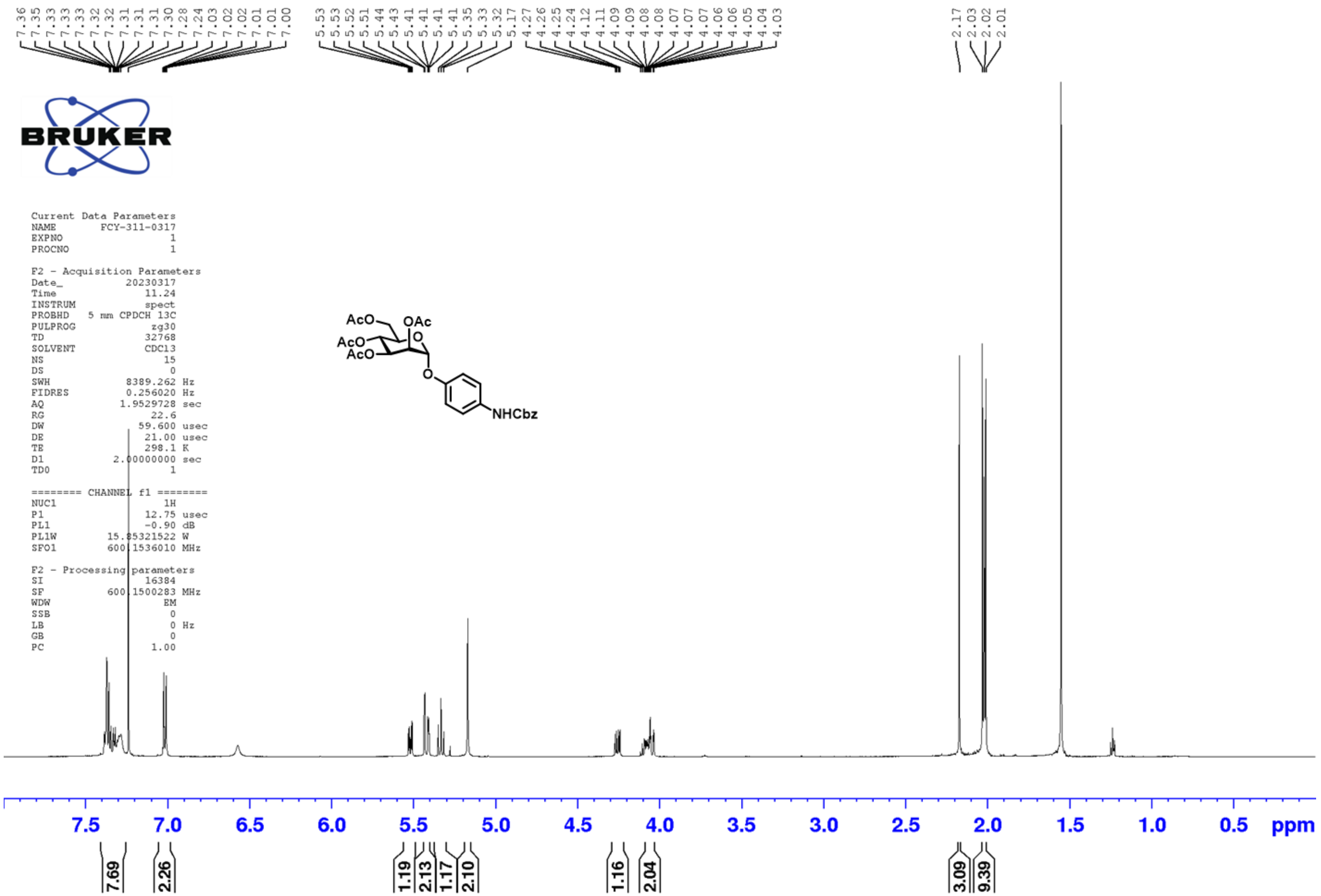

7.36 7.35 7.33 7.33 7.33 7.32 7.32 7.31 7.31 7.31 7.30 7.28 7.24 7.03 7.02 7.02 7.01 7.01 7.00
5.53 5.53 5.52 5.51 5.44 5.43 5.41 5.41 5.41 5.41 5.35 5.33 5.32 5.17
4.27 4.26 4.25 4.24 4.12 4.11 4.09 4.09 4.08 4.08 4.07 4.07 4.07 4.06 4.06 4.05 4.04 4.03
2.17 2.03 2.02 2.01
BRUKER
Current Data Parameters
NAME FCY-311-0317
EXPNO 1
PROCNO 1
F2 - Acquisition Parameters
Date_ 20230317
Time 11.24
INSTRUM spect
PROBHD 5 mm CPDCH 13C
PULPROG zg30
TD 32768
SOLVENT CDCl3
NS 15
DS 0
SWH 8389.262 Hz
FIDRES 0.256020 Hz
AQ 1.9529728 sec
RG 22.6
DW 59.600 usec
DE 21.00 usec
TE 298.1 K
D1 2.00000000 sec
TD0 1
======== CHANNEL f1 ========
NUC1 1H
P1 12.75 usec
PL1 -0.90 dB
PL1W 15.85321522 W
SFO1 600.1536010 MHz
F2 - Processing parameters
SI 16384
SF 600.1500283 MHz
WDW EM
SSB 0
LB 0 Hz
GB 0
PC 1.00
AcO
OAc
AcO
AcO
O
O
NHCbz
7.5 7.0 6.5 6.0 5.5 5.0 4.5 4.0 3.5 3.0 2.5 2.0 1.5 1.0 0.5 ppm
7.69 2.26 1.19 2.13 1.17 2.10 1.16 2.04 3.09 9.39

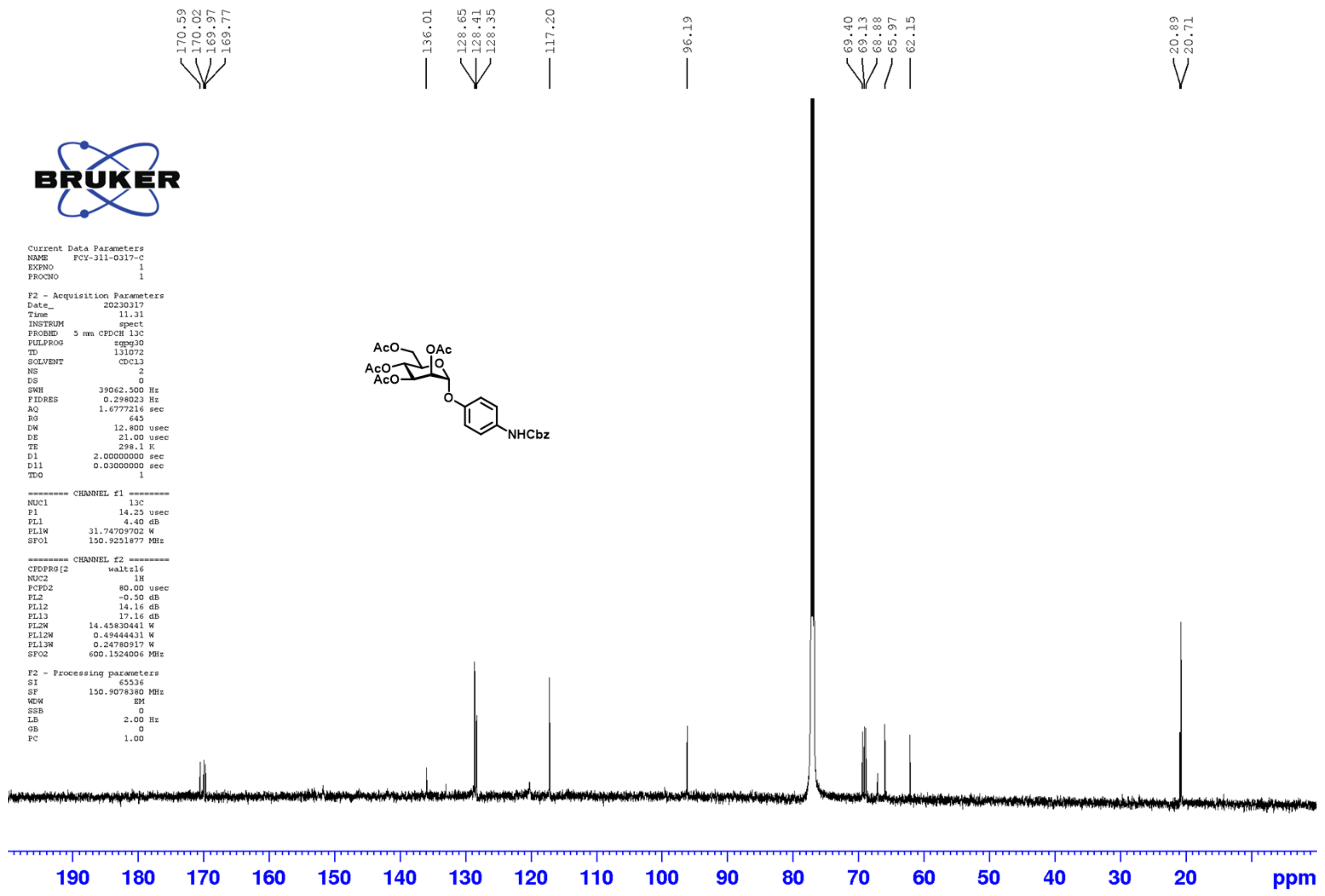
170.59
170.02
169.97
169.77
136.01
128.65
128.41
128.35
117.20
96.19
69.40
69.13
68.88
65.97
62.15
20.89
20.71
BRUKER
Current Data Parameters
NAME FCY-311-0317-C
EXPNO 1
PROCNO 1
F2 - Acquisition Parameters
Date_ 20230317
Time 11.31
INSTRUM spect
PROBHD 5 mm CPDCH 13C
PULPROG zgpg30
TD 131072
SOLVENT CDCl3
NS 2
DS 0
SWH 39062.500 Hz
FIDRES 0.298023 Hz
AQ 1.6777216 sec
RG 645
DW 12.800 usec
DE 21.00 usec
TE 298.1 K
D1 2.00000000 sec
D11 0.03000000 sec
TD0 1
======== CHANNEL f1 ========
NUC1 13C
P1 14.25 usec
PL1 4.40 dB
PL1W 31.74709702 W
SFO1 150.9251877 MHz
======== CHANNEL f2 ========
CPDPRG[2 waltz16
NUC2 1H
PCPD2 80.00 usec
PL2 -0.50 dB
PL12 14.16 dB
PL13 17.16 dB
PL2W 14.45830441 W
PL12W 0.49444431 W
PL13W 0.24780917 W
SFO2 600.1524006 MHz
F2 - Processing parameters
SI 65536
SF 150.9078380 MHz
WDW EM
SSB 0
LB 2.00 Hz
GB 0
PC 1.00
AcO
OAc
AcO
AcO
O
O
NHCbz
190
180
170
160
150
140
130
120
110
100
90
80
70
60
50
40
30
20
ppm

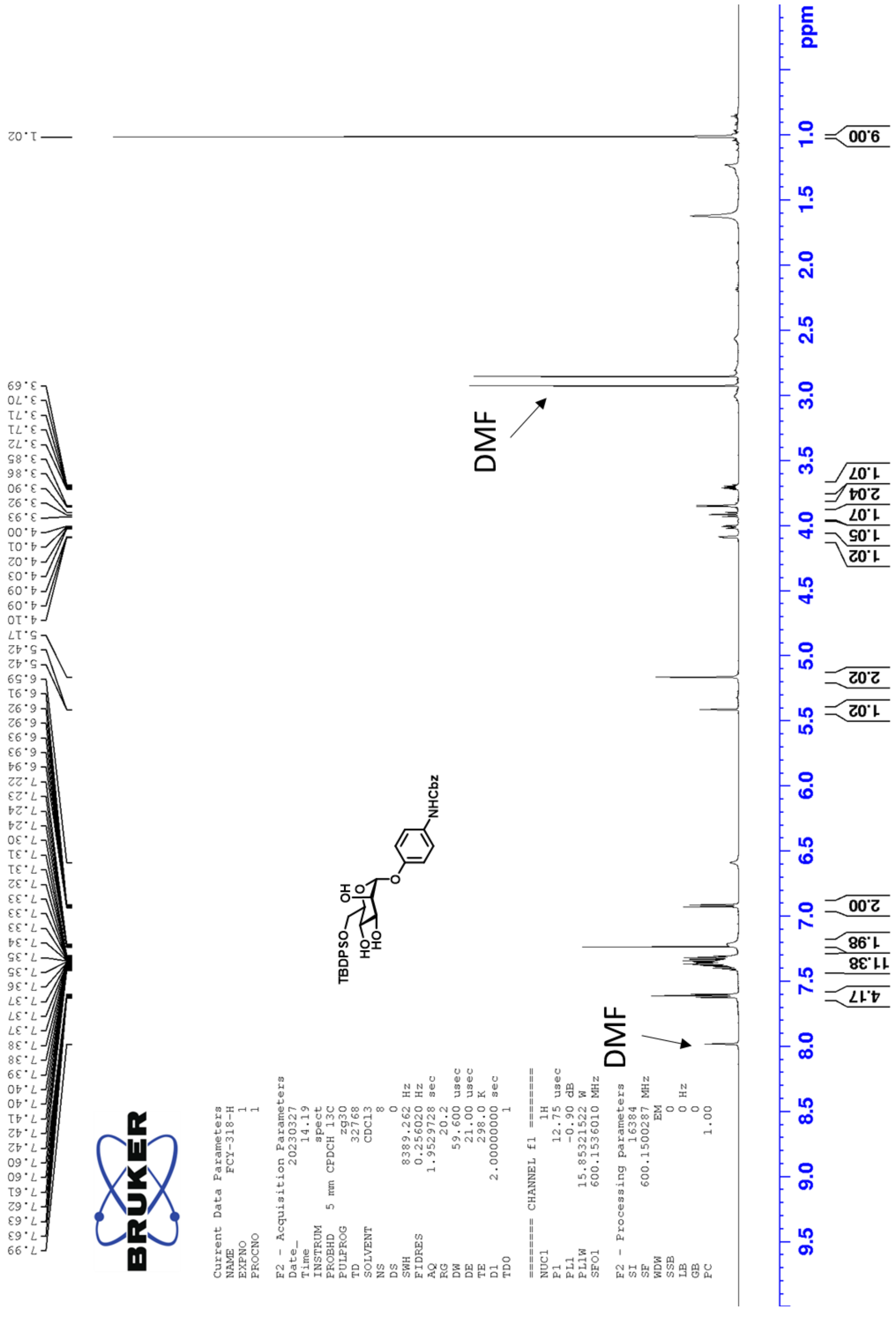
DMF
DMF
ppm
9.00
1.07
2.04
1.07
1.05
1.02
2.02
1.02
2.00
1.98
11.38
4.17
TBDPSO
HO
HO
OH
NHCbz
BRUKER
Current Data Parameters
NAME FCY-318-H
EXPNO 1
PROCNO 1
F2 - Acquisition Parameters
Date_ 20230327
Time 14.19
INSTRUM spect
PROBHD 5 mm CPDCH 13C
PULPROG zg30
TD 32768
SOLVENT CDCl3
NS 8
DS 0
SWH 8389.262 Hz
FIDRES 0.256020 Hz
AQ 1.9529728 sec
RG 20.2
DW 59.600 usec
DE 21.00 usec
TE 298.0 K
D1 2.00000000 sec
TD0 1
======= CHANNEL f1 =======
NUC1 1H
P1 12.75 usec
PL1 -0.90 dB
PL1W 15.85321522 W
SFO1 600.1536010 MHz
F2 - Processing parameters
SI 16384
SF 600.1500287 MHz
WDW EM
SSB 0
LB 0 Hz
GB 0
PC 1.00

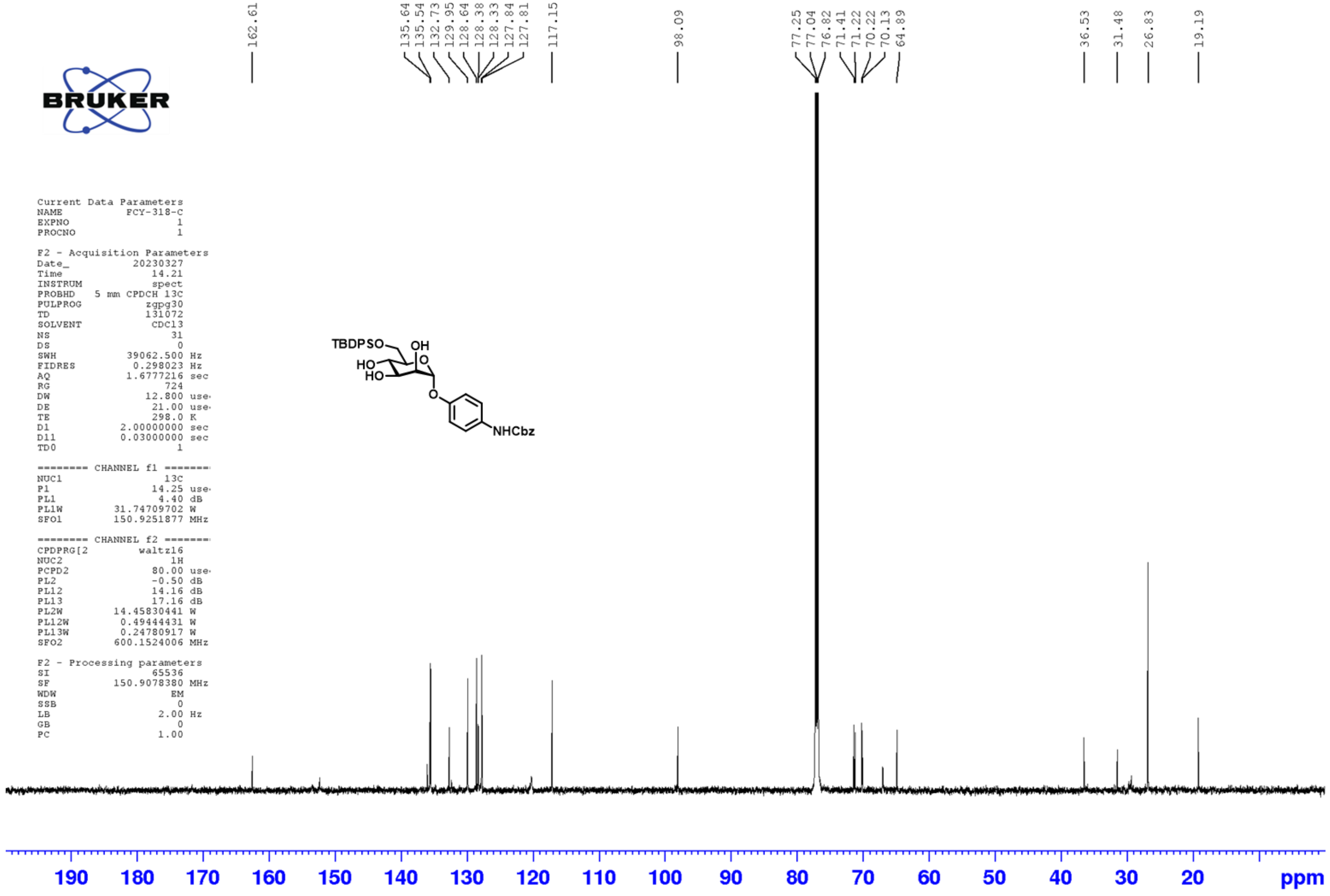
162.61
135.64
135.54
132.73
129.95
128.64
128.38
128.33
127.84
127.81
117.15
98.09
77.25
77.04
76.82
71.41
71.22
70.22
70.13
64.89
36.53
31.48
26.83
19.19
BRUKER
Current Data Parameters
NAME FCY-318-C
EXPNO 1
PROCNO 1
F2 - Acquisition Parameters
Date_ 20230327
Time 14.21
INSTRUM spect
PROBHD 5 mm CPDCH 13C
PULPROG zgpg30
TD 131072
SOLVENT CDCl3
NS 31
DS 0
SWH 39062.500 Hz
FIDRES 0.298023 Hz
AQ 1.6777216 sec
RG 724
DW 12.800 use
DE 21.00 use
TE 298.0 K
D1 2.00000000 sec
D11 0.03000000 sec
TD0 1
======== CHANNEL f1 ========
NUC1 13C
P1 14.25 use
PL1 4.40 dB
PL1W 31.74709702 W
SFO1 150.9251877 MHz
======== CHANNEL f2 ========
CPDPRG[2 waltz16
NUC2 1H
PCPD2 80.00 use
PL2 -0.50 dB
PL12 14.16 dB
PL13 17.16 dB
PL2W 14.45830441 W
PL12W 0.49444431 W
PL13W 0.24780917 W
SFO2 600.1524006 MHz
F2 - Processing parameters
SI 65536
SF 150.9078380 MHz
WDW EM
SSB 0
LB 2.00 Hz
GB 0
PC 1.00
TBDPSO
OH
HO
HO
O
O
NHCbz
190 180 170 160 150 140 130 120 110 100 90 80 70 60 50 40 30 20 ppm

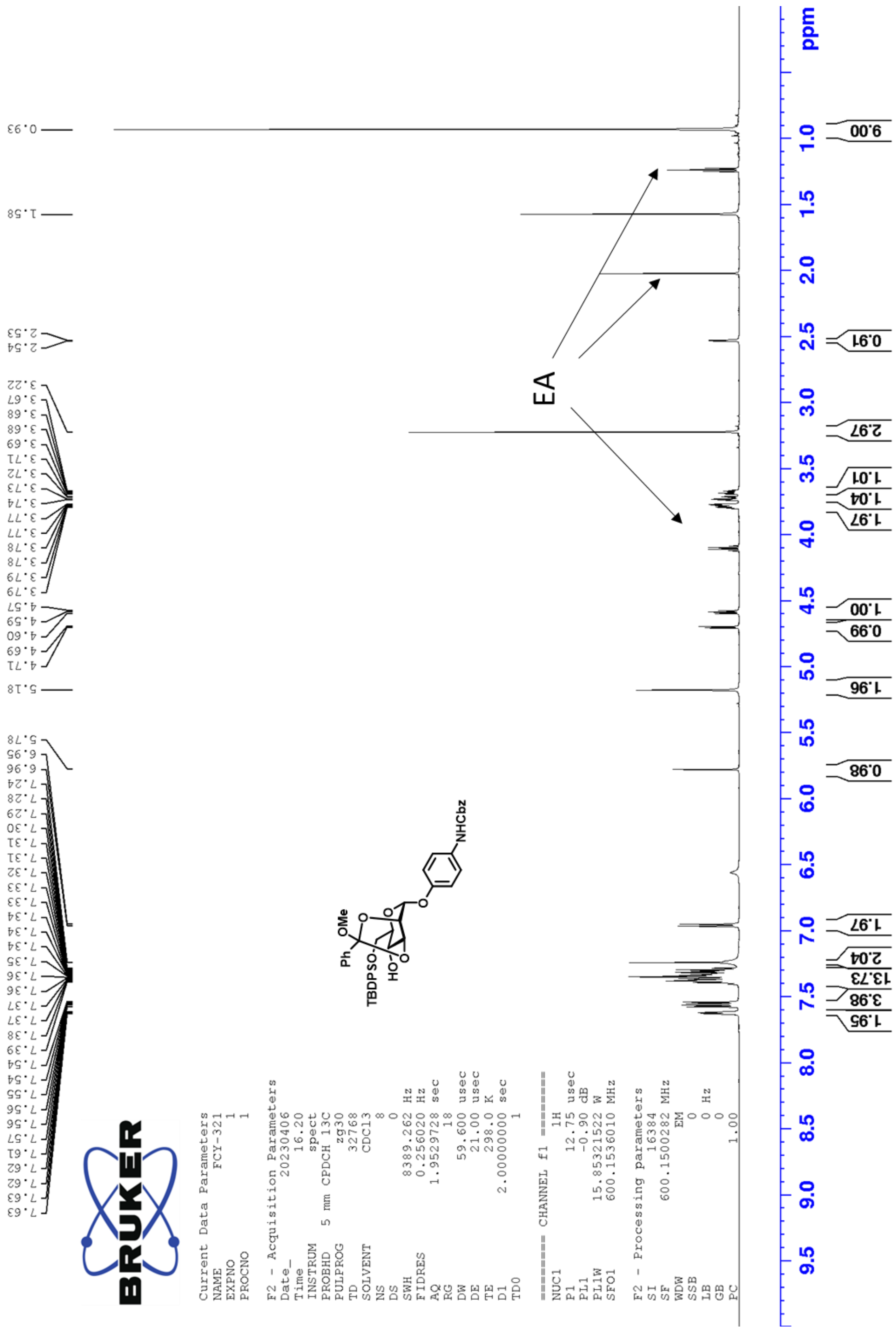
0.93
1.58
2.53
2.54
3.22
3.67
3.68
3.68
3.69
3.71
3.72
3.73
3.74
3.77
3.77
3.78
3.78
3.79
3.79
4.57
4.59
4.60
4.69
4.71
5.18
5.78
6.95
6.96
7.24
7.28
7.29
7.30
7.31
7.31
7.32
7.33
7.33
7.34
7.34
7.34
7.35
7.36
7.36
7.37
7.37
7.38
7.39
7.54
7.54
7.55
7.56
7.56
7.57
7.61
7.62
7.62
7.63
7.63
BRUKER
Current Data Parameters
NAME FCY-321
EXPNO 1
PROCNO 1
F2 - Acquisition Parameters
Date_ 20230406
Time 16.20
INSTRUM spect
PROBHD 5 mm CPDCH 13C
PULPROG zg30
TD 32768
SOLVENT CDCl3
NS 8
DS 0
SWH 8389.262 Hz
FIDRES 0.256020 Hz
AQ 1.9529728 sec
RG 18
DW 59.600 usec
DE 21.00 usec
TE 298.0 K
D1 2.00000000 sec
TD0 1
======== CHANNEL f1 ========
NUC1 1H
P1 12.75 usec
PL1 -0.90 dB
PL1W 15.85321522 W
SFO1 600.1536010 MHz
F2 - Processing parameters
SI 16384
SF 600.1500282 MHz
WDW EM
SSB 0
LB 0 Hz
GB 0
PC 1.00
NHCbz
Ph
OMe
TBDPSO
HO
EA
9.00
0.91
2.97
1.01
1.04
1.97
1.00
0.99
1.96
0.98
1.97
2.04
13.73
3.98
1.95
1.0
1.5
2.0
2.5
3.0
3.5
4.0
4.5
5.0
5.5
6.0
6.5
7.0
7.5
8.0
8.5
9.0
9.5
ppm

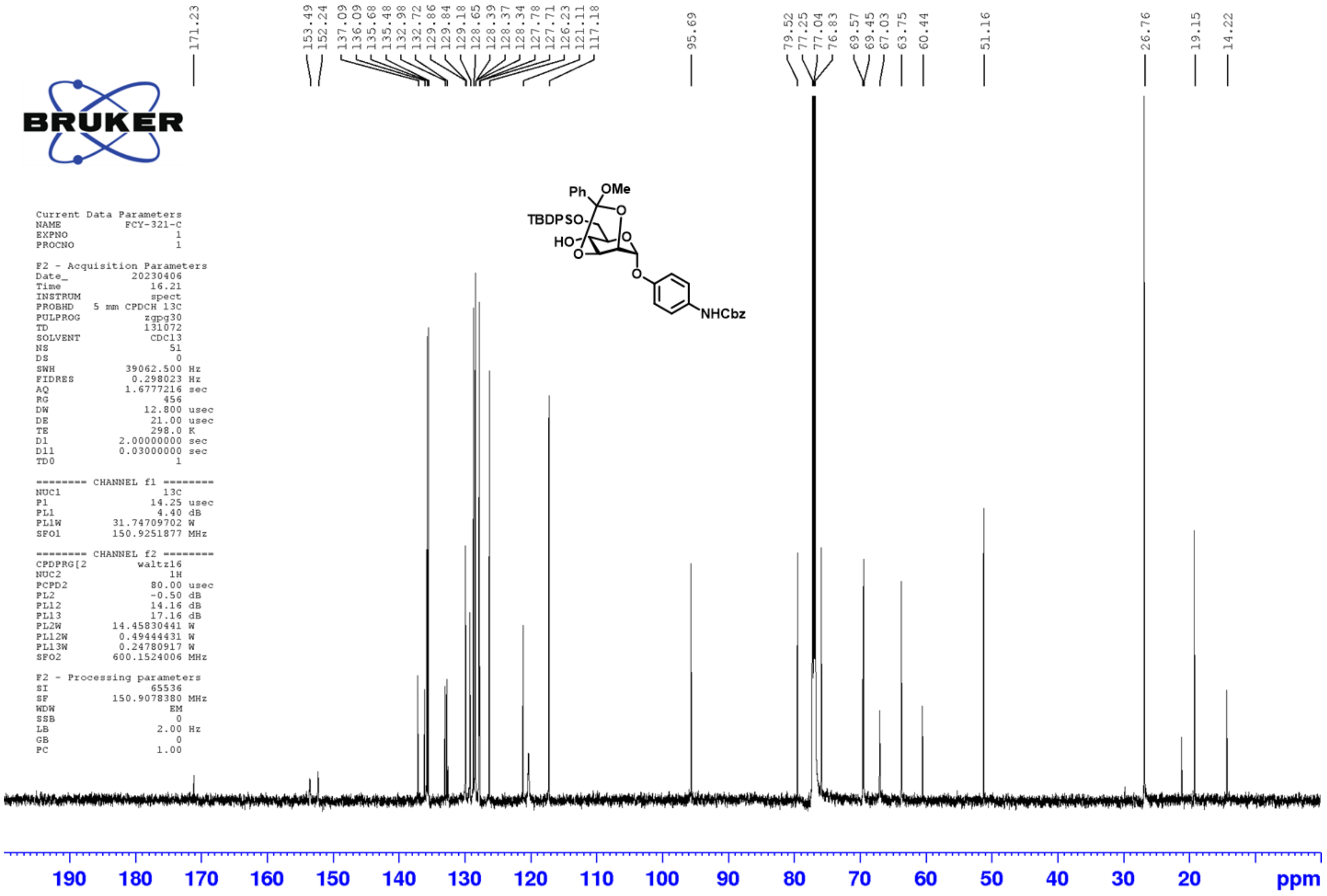


```
Current Data Parameters
NAME          FCY-321-C
EXPNO                 1
PROCNO                1

F2 - Acquisition Parameters
Date_          20230406
Time              16.21
INSTRUM           spect
PROBHD   5 mm CPDCH 13C
PULPROG          zgpg30
TD               131072
SOLVENT           CDCl3
NS                   51
DS                    0
SWH           39062.500 Hz
FIDRES         0.298023 Hz
AQ            1.6777216 sec
RG                  456
DW               12.800 usec
DE                21.00 usec
TE                298.0 K
D1           2.00000000 sec
D11          0.03000000 sec
TD0                   1

======== CHANNEL f1 ========
NUC1                13C
P1                14.25 usec
PL1                4.40 dB
PL1W        31.74709702 W
SFO1        150.9251877 MHz

======== CHANNEL f2 ========
CPDPRG[2        waltz16
NUC2                 1H
PCPD2             80.00 usec
PL2               -0.50 dB
PL12              14.16 dB
PL13              17.16 dB
PL2W        14.45830441 W
PL12W        0.49444431 W
PL13W        0.24780917 W
SFO2        600.1524006 MHz

F2 - Processing parameters
SI                65536
SF          150.9078380 MHz
WDW                  EM
SSB                   0
LB                 2.00 Hz
GB                    0
PC                 1.00
```

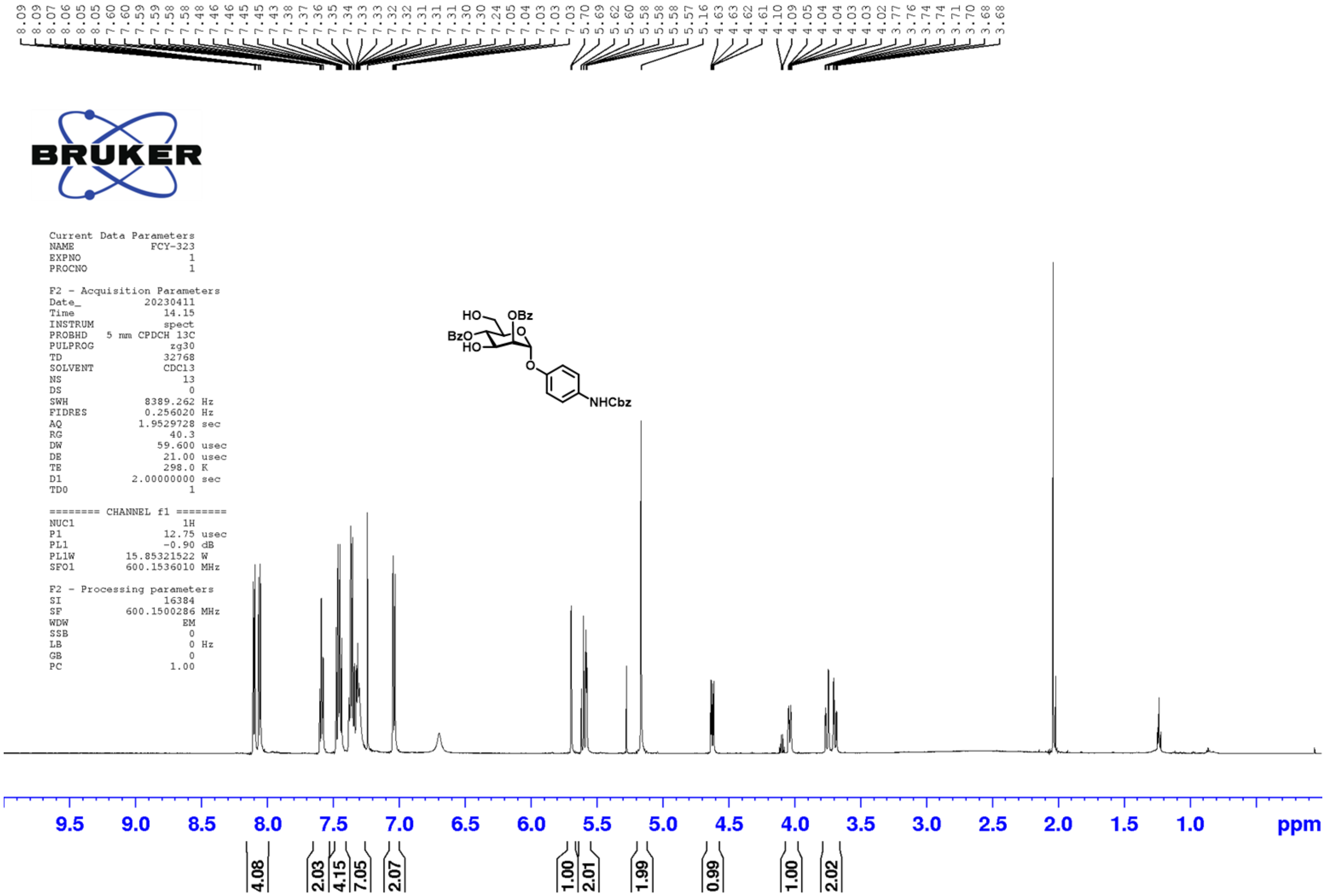
8.09 8.09 8.07 8.06 8.05 8.05 7.60 7.60 7.59 7.59 7.58 7.58 7.48 7.46 7.46 7.45 7.45 7.43 7.38 7.37 7.36 7.35 7.34 7.33 7.33 7.32 7.32 7.31 7.31 7.31 7.30 7.30 7.24 7.05 7.04 7.03 7.03 7.03 5.70 5.69 5.62 5.60 5.58 5.58 5.58 5.57 5.16 4.63 4.63 4.62 4.61 4.10 4.09 4.05 4.04 4.04 4.03 4.03 4.02 3.77 3.76 3.74 3.74 3.71 3.70 3.68 3.68
BRUKER
Current Data Parameters
NAME FCY-323
EXPNO 1
PROCNO 1
F2 - Acquisition Parameters
Date_ 20230411
Time 14.15
INSTRUM spect
PROBHD 5 mm CPDCH 13C
PULPROG zg30
TD 32768
SOLVENT CDCl3
NS 13
DS 0
SWH 8389.262 Hz
FIDRES 0.256020 Hz
AQ 1.9529728 sec
RG 40.3
DW 59.600 usec
DE 21.00 usec
TE 298.0 K
D1 2.00000000 sec
TD0 1
======== CHANNEL f1 ========
NUC1 1H
P1 12.75 usec
PL1 -0.90 dB
PL1W 15.85321522 W
SFO1 600.1536010 MHz
F2 - Processing parameters
SI 16384
SF 600.1500286 MHz
WDW EM
SSB 0
LB 0 Hz
GB 0
PC 1.00
HO
OBz
BzO
HO
O
O
NHCbz
9.5 9.0 8.5 8.0 7.5 7.0 6.5 6.0 5.5 5.0 4.5 4.0 3.5 3.0 2.5 2.0 1.5 1.0 ppm
4.08 2.03 4.15 7.05 2.07 1.00 2.01 1.99 0.99 1.00 2.02

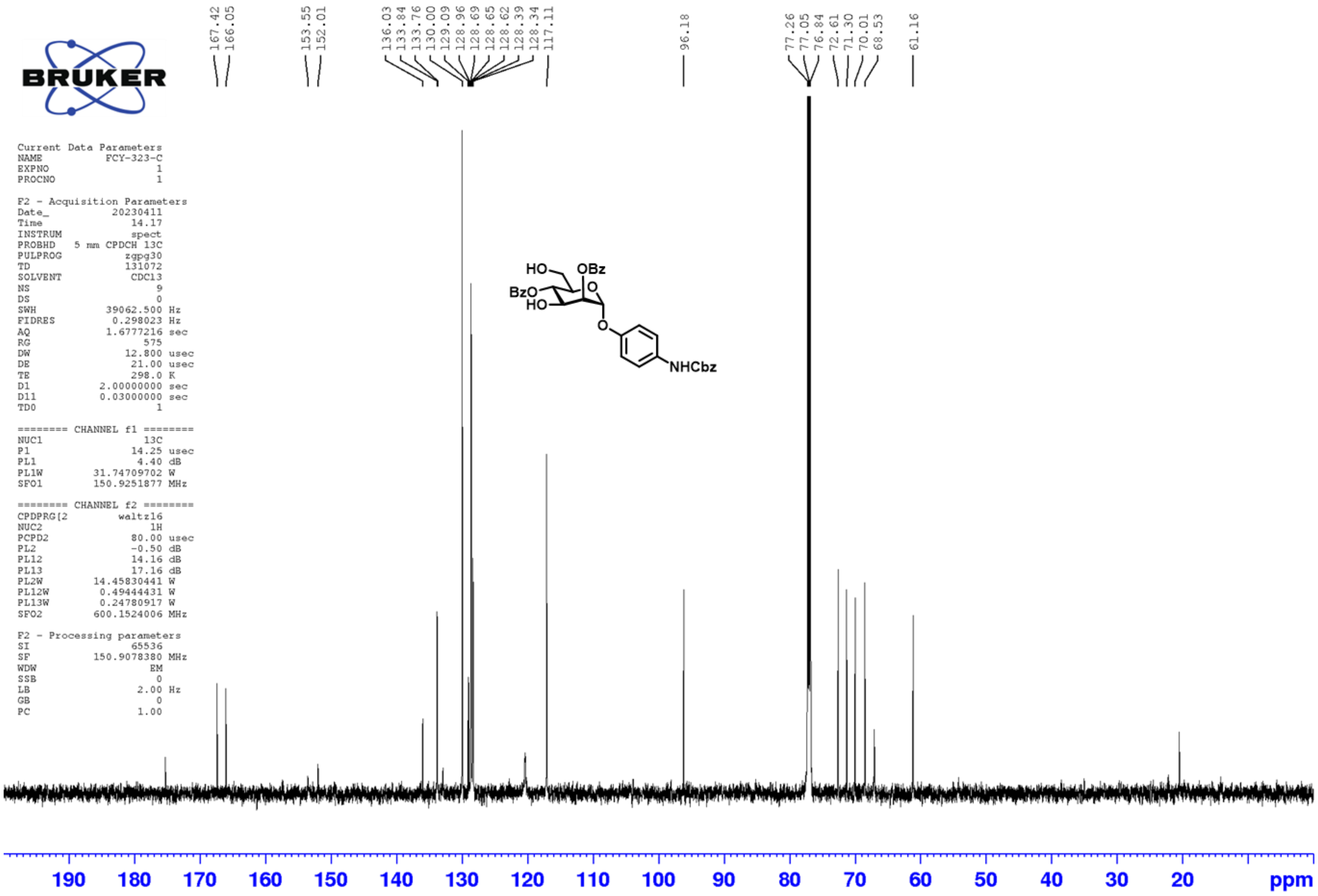
167.42
166.05
153.55
152.01
136.03
133.84
133.76
130.00
129.09
128.96
128.69
128.65
128.62
128.39
128.34
117.11
96.18
77.26
77.05
76.84
72.61
71.30
70.01
68.53
61.16
BRUKER
Current Data Parameters
NAME FCY-323-C
EXPNO 1
PROCNO 1
F2 - Acquisition Parameters
Date_ 20230411
Time 14.17
INSTRUM spect
PROBHD 5 mm CPDCH 13C
PULPROG zgpg30
TD 131072
SOLVENT CDCl3
NS 9
DS 0
SWH 39062.500 Hz
FIDRES 0.298023 Hz
AQ 1.6777216 sec
RG 575
DW 12.800 usec
DE 21.00 usec
TE 298.0 K
D1 2.00000000 sec
D11 0.03000000 sec
TD0 1
======== CHANNEL f1 ========
NUC1 13C
P1 14.25 usec
PL1 4.40 dB
PL1W 31.74709702 W
SFO1 150.9251877 MHz
======== CHANNEL f2 ========
CPDPRG[2 waltz16
NUC2 1H
PCPD2 80.00 usec
PL2 -0.50 dB
PL12 14.16 dB
PL13 17.16 dB
PL2W 14.45830441 W
PL12W 0.49444431 W
PL13W 0.24780917 W
SFO2 600.1524006 MHz
F2 - Processing parameters
SI 65536
SF 150.9078380 MHz
WDW EM
SSB 0
LB 2.00 Hz
GB 0
PC 1.00
HO
OBz
BzO
HO
O
O
NHCbz
190 180 170 160 150 140 130 120 110 100 90 80 70 60 50 40 30 20 ppm

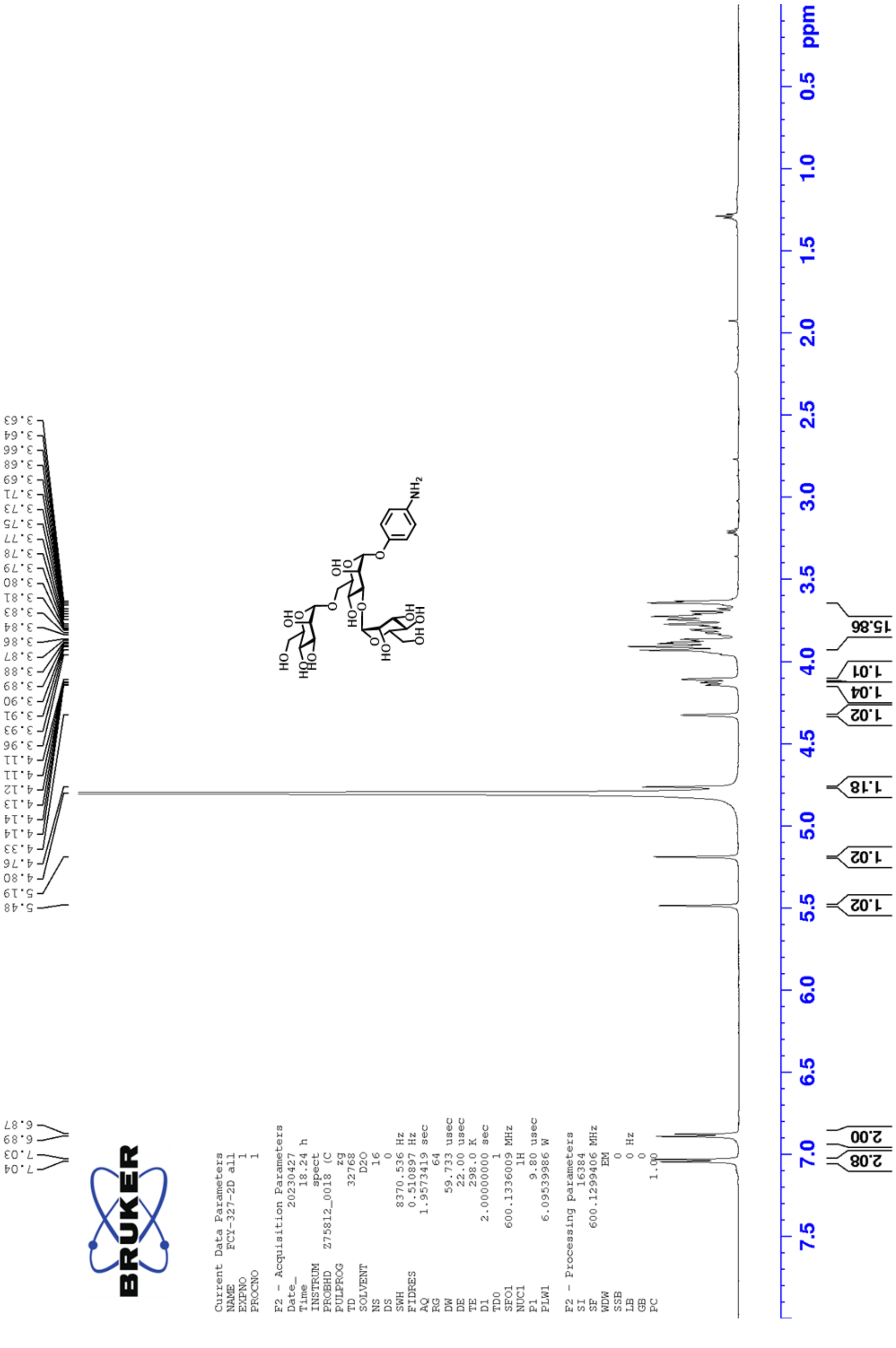
5.48
5.19
4.80
4.76
4.33
4.14
4.14
4.13
4.12
4.11
4.11
3.96
3.93
3.91
3.90
3.89
3.88
3.87
3.86
3.84
3.83
3.81
3.80
3.79
3.78
3.77
3.75
3.73
3.71
3.69
3.68
3.66
3.64
3.63
7.04
7.03
6.89
6.87
BRUKER
Current Data Parameters
NAME FCY-327-2D all
EXPNO 1
PROCNO 1
F2 - Acquisition Parameters
Date_ 20230427
Time 18.24 h
INSTRUM spect
PROBHD Z75812_0018 (C
PULPROG zg
TD 32768
SOLVENT D2O
NS 16
DS 0
SWH 8370.536 Hz
FIDRES 0.510897 Hz
AQ 1.9573419 sec
RG 64
DW 59.733 usec
DE 22.00 usec
TE 298.0 K
D1 2.00000000 sec
TD0 1
SFO1 600.1336009 MHz
NUC1 1H
P1 9.80 usec
PLW1 6.09539986 W
F2 - Processing parameters
SI 16384
SF 600.1299406 MHz
WDW EM
SSB 0
LB 0 Hz
GB 0
PC 1.00
HO
OH
NH2
15.86
1.01
1.04
1.02
1.18
1.02
1.02
2.00
2.08
ppm
0.5
1.0
1.5
2.0
2.5
3.0
3.5
4.0
4.5
5.0
5.5
6.0
6.5
7.0
7.5

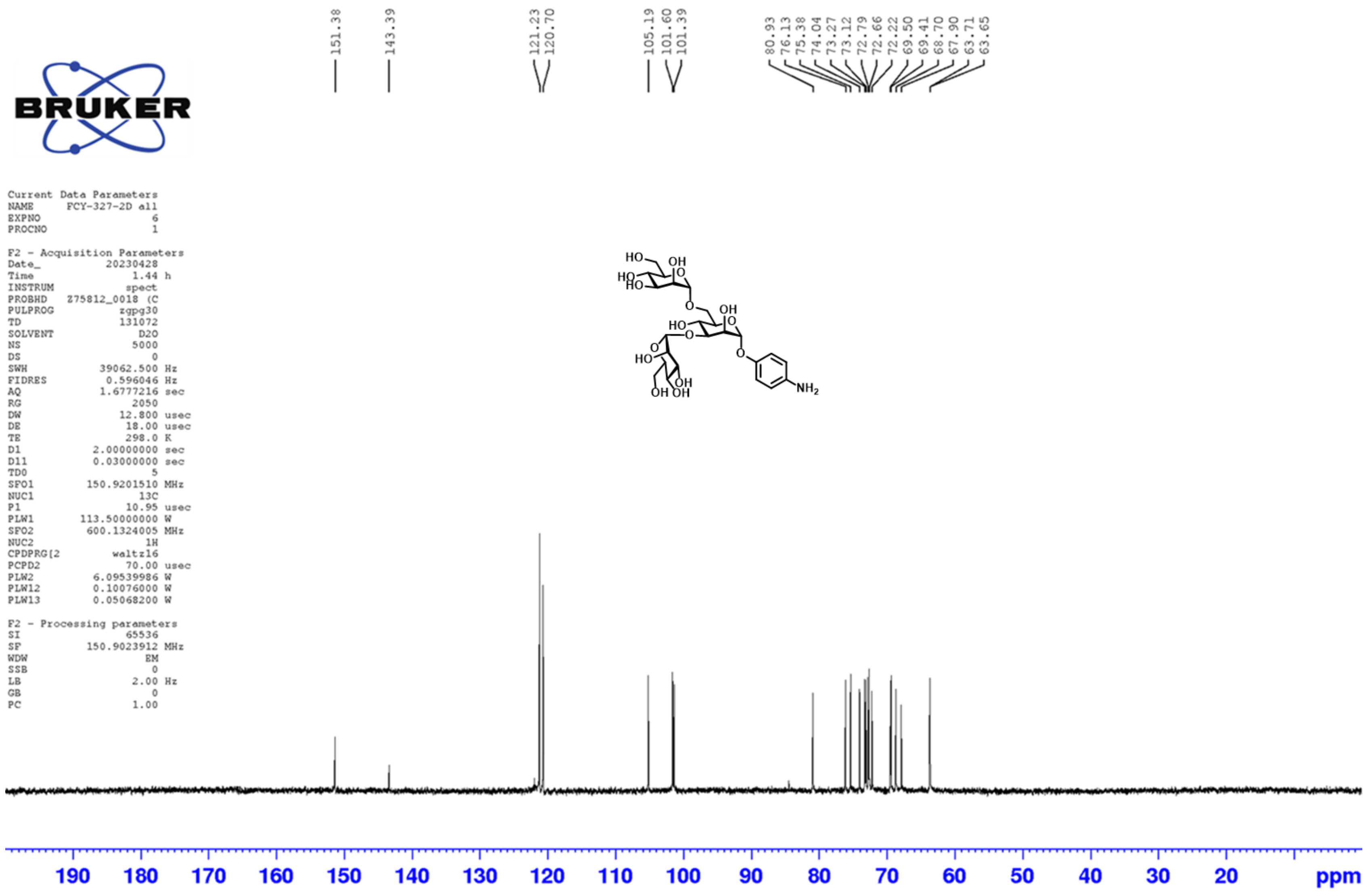
151.38
143.39
121.23
120.70
105.19
101.60
101.39
80.93
76.13
75.38
74.04
73.27
73.12
72.79
72.66
72.22
69.50
69.41
68.70
67.90
63.71
63.65
BRUKER
Current Data Parameters
NAME FCY-327-2D all
EXPNO 6
PROCNO 1
F2 - Acquisition Parameters
Date_ 20230428
Time 1.44 h
INSTRUM spect
PROBHD Z75812_0018 (C
PULPROG zgpg30
TD 131072
SOLVENT D2O
NS 5000
DS 0
SWH 39062.500 Hz
FIDRES 0.596046 Hz
AQ 1.6777216 sec
RG 2050
DW 12.800 usec
DE 18.00 usec
TE 298.0 K
D1 2.00000000 sec
D11 0.03000000 sec
TD0 5
SFO1 150.9201510 MHz
NUC1 13C
P1 10.95 usec
PLW1 113.50000000 W
SFO2 600.1324005 MHz
NUC2 1H
CPDPRG[2 waltz16
PCPD2 70.00 usec
PLW2 6.09539986 W
PLW12 0.10076000 W
PLW13 0.05068200 W
F2 - Processing parameters
SI 65536
SF 150.9023912 MHz
WDW EM
SSB 0
LB 2.00 Hz
GB 0
PC 1.00
190 180 170 160 150 140 130 120 110 100 90 80 70 60 50 40 30 20 ppm

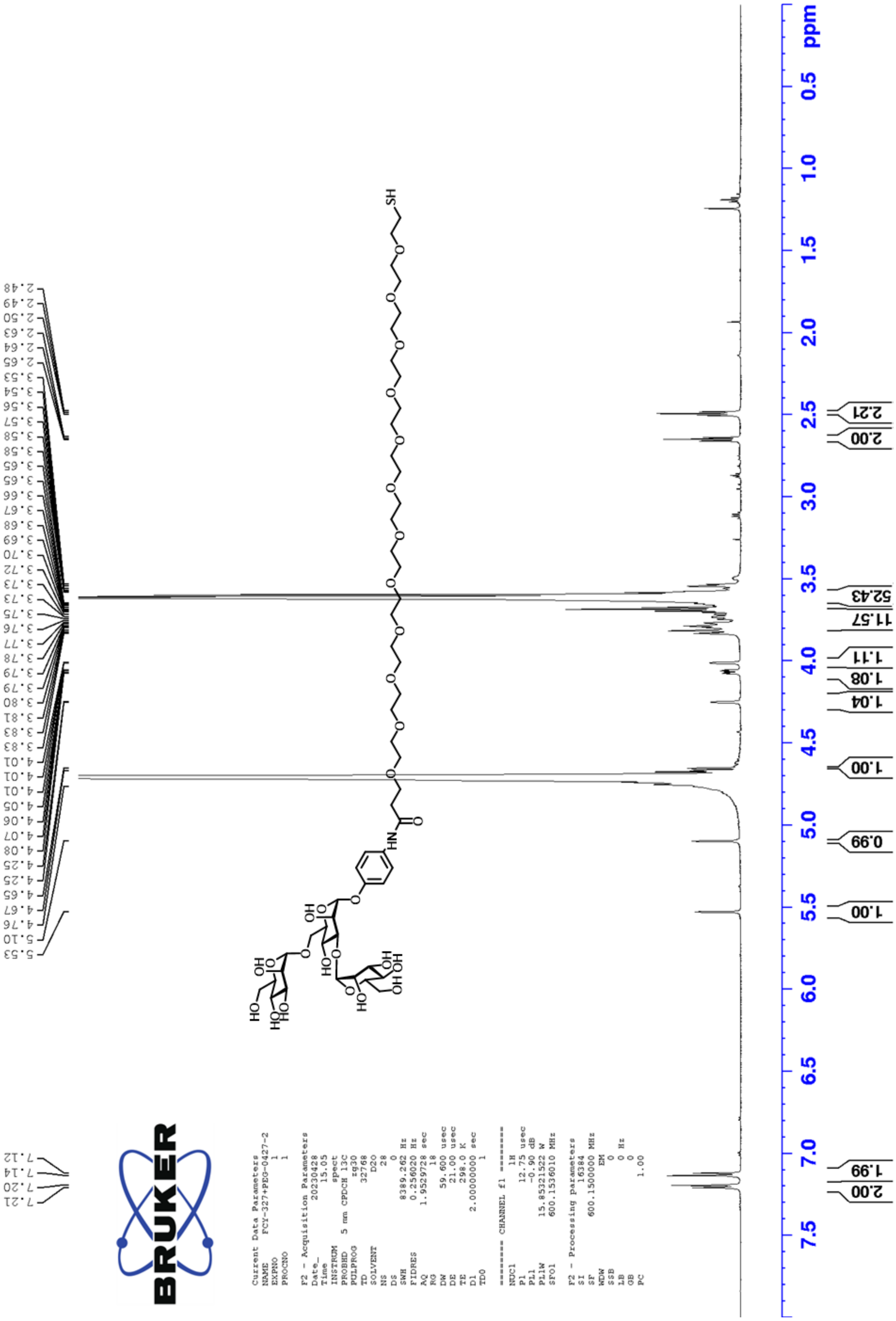
ppm
0.5
1.0
1.5
2.0
2.5
3.0
3.5
4.0
4.5
5.0
5.5
6.0
6.5
7.0
7.5
2.21
2.00
52.43
11.57
1.11
1.08
1.04
1.00
0.99
1.00
1.99
2.00
SH
HN
BRUKER

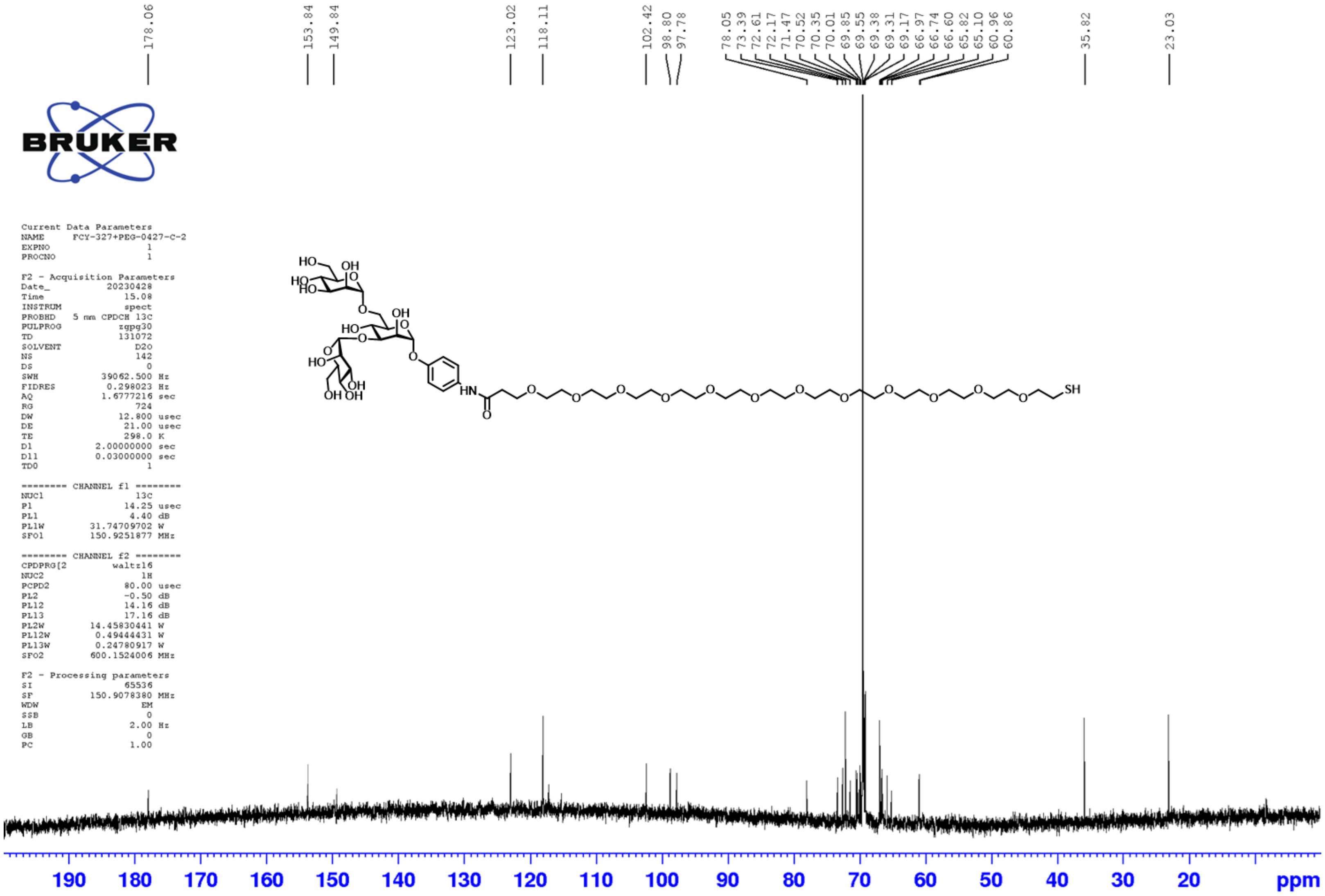
178.06
153.84
149.84
123.02
118.11
102.42
98.80
97.78
78.05
73.39
72.61
72.17
71.47
70.52
70.35
70.01
69.85
69.55
69.38
69.31
69.17
66.97
66.74
66.60
65.82
65.10
60.96
60.86
35.82
23.03
BRUKER
Current Data Parameters
NAME FCY-327+PEG-0427-C-2
EXPNO 1
PROCNO 1
F2 - Acquisition Parameters
Date_ 20230428
Time 15.08
INSTRUM spect
PROBHD 5 mm CPDCH 13C
PULPROG zgpg30
TD 131072
SOLVENT D2O
NS 142
DS 0
SWH 39062.500 Hz
FIDRES 0.298023 Hz
AQ 1.6777216 sec
RG 724
DW 12.800 usec
DE 21.00 usec
TE 298.0 K
D1 2.00000000 sec
D11 0.03000000 sec
TD0 1
======== CHANNEL f1 ========
NUC1 13C
P1 14.25 usec
PL1 4.40 dB
PL1W 31.74709702 W
SFO1 150.9251877 MHz
======== CHANNEL f2 ========
CPDPRG[2 waltz16
NUC2 1H
PCPD2 80.00 usec
PL2 -0.50 dB
PL12 14.16 dB
PL13 17.16 dB
PL2W 14.45830441 W
PL12W 0.49444431 W
PL13W 0.24780917 W
SFO2 600.1524006 MHz
F2 - Processing parameters
SI 65536
SF 150.9078380 MHz
WDW EM
SSB 0
LB 2.00 Hz
GB 0
PC 1.00
HO
OH
HO
O
OH
NH
SH
190
180
170
160
150
140
130
120
110
100
90
80
70
60
50
40
30
20
ppm

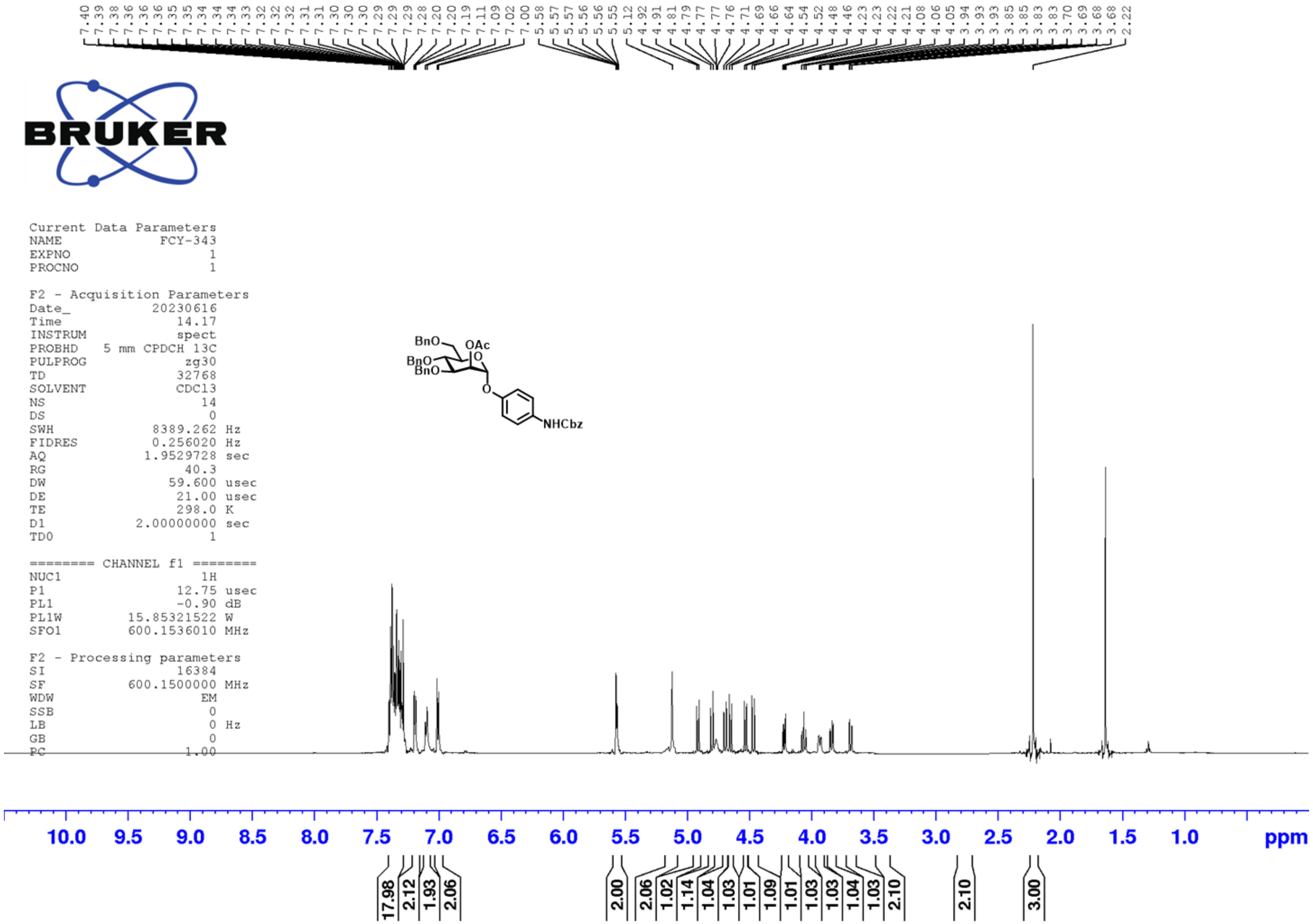
7.40 7.39 7.38 7.36 7.36 7.36 7.35 7.35 7.34 7.34 7.34 7.33 7.32 7.32 7.32 7.31 7.31 7.30 7.30 7.30 7.29 7.29 7.29 7.28 7.20 7.20 7.19 7.11 7.09 7.02 7.00 5.58 5.57 5.57 5.56 5.56 5.55 5.12 4.92 4.91 4.81 4.79 4.77 4.77 4.76 4.71 4.69 4.66 4.64 4.54 4.52 4.48 4.46 4.23 4.23 4.22 4.21 4.08 4.06 4.05 3.94 3.93 3.93 3.93 3.85 3.85 3.83 3.83 3.70 3.69 3.68 3.68 2.22
BRUKER
Current Data Parameters
NAME FCY-343
EXPNO 1
PROCNO 1
F2 - Acquisition Parameters
Date_ 20230616
Time_ 14.17
INSTRUM spect
PROBHD 5 mm CPDCH 13C
PULPROG zg30
TD 32768
SOLVENT CDCl3
NS 14
DS 0
SWH 8389.262 Hz
FIDRES 0.256020 Hz
AQ 1.9529728 sec
RG 40.3
DW 59.600 usec
DE 21.00 usec
TE 298.0 K
D1 2.00000000 sec
TD0 1
======== CHANNEL f1 ========
NUC1 1H
P1 12.75 usec
PL1 -0.90 dB
PL1W 15.85321522 W
SFO1 600.1536010 MHz
F2 - Processing parameters
SI 16384
SF 600.1500000 MHz
WDW EM
SSB 0
LB 0 Hz
GB 0
PC 1.00
BnO
OAc
BnO
BnO
O
O
NHCbz
10.0 9.5 9.0 8.5 8.0 7.5 7.0 6.5 6.0 5.5 5.0 4.5 4.0 3.5 3.0 2.5 2.0 1.5 1.0 ppm
17.98 2.12 1.93 2.06 2.00 2.06 1.02 1.14 1.04 1.03 1.01 1.09 1.01 1.03 1.03 1.04 1.03 2.10 2.10 3.00

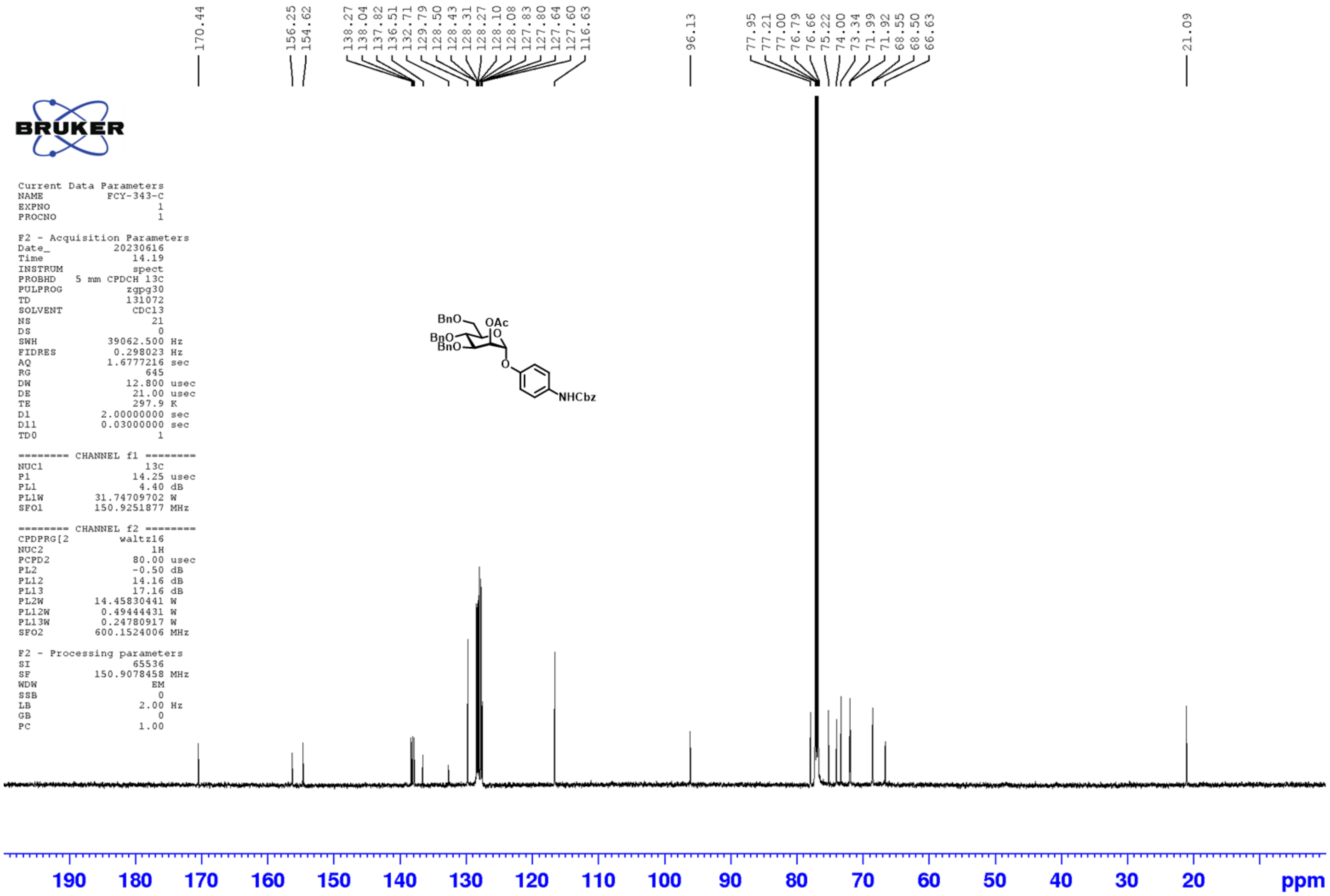
170.44
156.25
154.62
138.27
138.04
137.82
136.51
132.71
129.79
128.50
128.43
128.31
128.27
128.10
128.08
127.83
127.80
127.64
127.60
116.63
96.13
77.95
77.21
77.00
76.79
76.66
75.22
74.00
73.34
71.99
71.92
68.55
68.50
66.63
21.09
BRUKER
Current Data Parameters
NAME FCY-343-C
EXPNO 1
PROCNO 1
F2 - Acquisition Parameters
Date_ 20230616
Time 14.19
INSTRUM spect
PROBHD 5 mm CPDCH 13C
PULPROG zgpg30
TD 131072
SOLVENT CDCl3
NS 21
DS 0
SWH 39062.500 Hz
FIDRES 0.298023 Hz
AQ 1.6777216 sec
RG 645
DW 12.800 usec
DE 21.00 usec
TE 297.9 K
D1 2.00000000 sec
D11 0.03000000 sec
TD0 1
======== CHANNEL f1 ========
NUC1 13C
P1 14.25 usec
PL1 4.40 dB
PL1W 31.74709702 W
SFO1 150.9251877 MHz
======== CHANNEL f2 ========
CPDPRG[2 waltz16
NUC2 1H
PCPD2 80.00 usec
PL2 -0.50 dB
PL12 14.16 dB
PL13 17.16 dB
PL2W 14.45830441 W
PL12W 0.49444431 W
PL13W 0.24780917 W
SFO2 600.1524006 MHz
F2 - Processing parameters
SI 65536
SF 150.9078458 MHz
WDW EM
SSB 0
LB 2.00 Hz
GB 0
PC 1.00
BnO
OAc
BnO
BnO
O
O
NHCbz
190
180
170
160
150
140
130
120
110
100
90
80
70
60
50
40
30
20
ppm

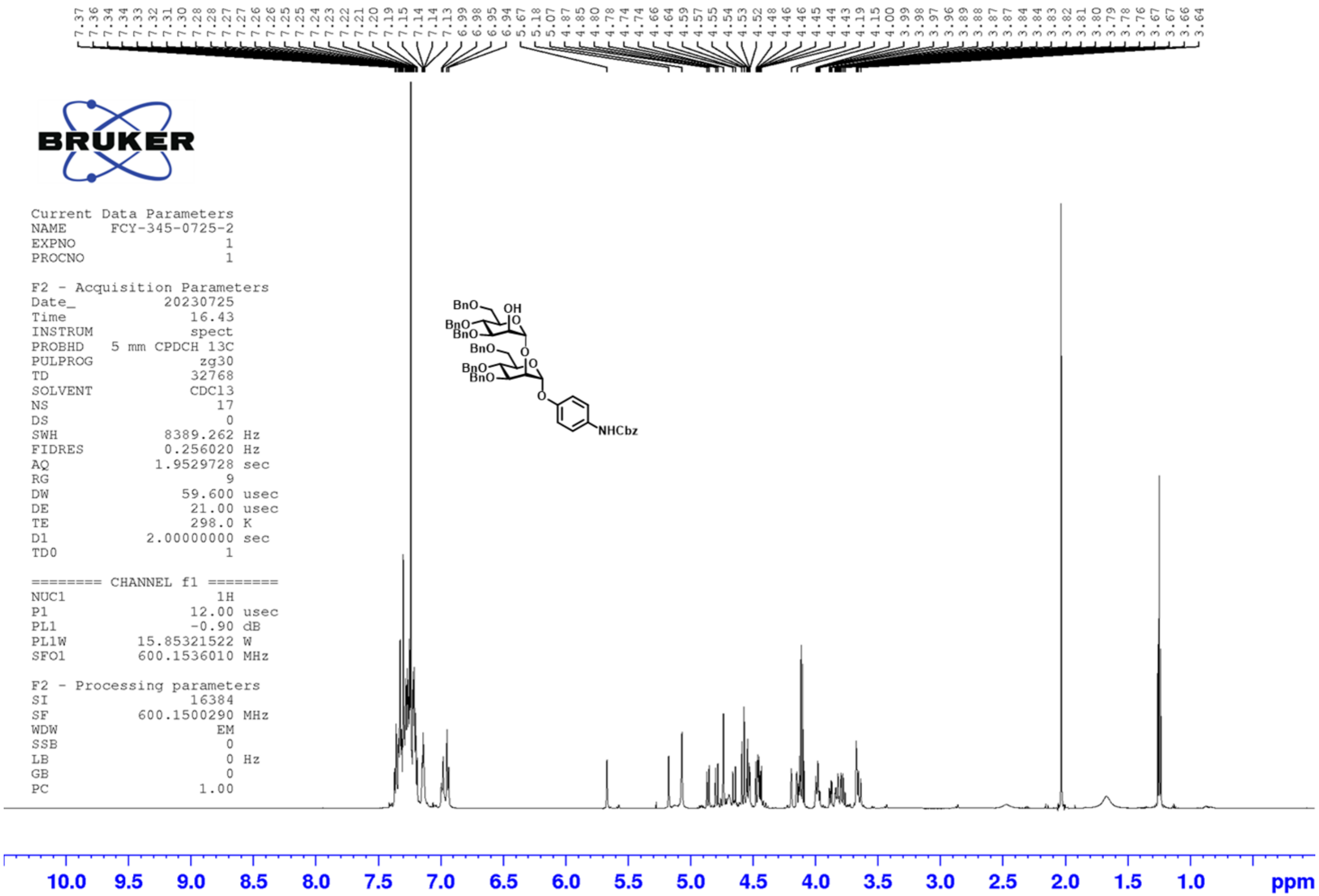
7.37 7.36 7.34 7.34 7.33 7.32 7.31 7.30 7.28 7.28 7.27 7.27 7.26 7.25 7.25 7.24 7.23 7.22 7.21 7.20 7.19 7.15 7.14 7.14 7.13 6.99 6.98 6.95 6.94 5.67 5.18 5.07 4.87 4.85 4.80 4.78 4.74 4.74 4.66 4.64 4.59 4.57 4.55 4.54 4.53 4.52 4.48 4.46 4.46 4.45 4.44 4.43 4.19 4.15 4.00 3.99 3.98 3.97 3.96 3.89 3.88 3.87 3.87 3.84 3.84 3.83 3.82 3.81 3.80 3.79 3.78 3.76 3.67 3.67 3.66 3.64
BRUKER
Current Data Parameters
NAME FCY-345-0725-2
EXPNO 1
PROCNO 1
F2 - Acquisition Parameters
Date_ 20230725
Time 16.43
INSTRUM spect
PROBHD 5 mm CPDCH 13C
PULPROG zg30
TD 32768
SOLVENT CDCl3
NS 17
DS 0
SWH 8389.262 Hz
FIDRES 0.256020 Hz
AQ 1.9529728 sec
RG 9
DW 59.600 usec
DE 21.00 usec
TE 298.0 K
D1 2.00000000 sec
TD0 1
======== CHANNEL f1 ========
NUC1 1H
P1 12.00 usec
PL1 -0.90 dB
PL1W 15.85321522 W
SFO1 600.1536010 MHz
F2 - Processing parameters
SI 16384
SF 600.1500290 MHz
WDW EM
SSB 0
LB 0 Hz
GB 0
PC 1.00
BnO
OH
BnO
BnO
BnO
BnO
BnO
NHCbz
10.0 9.5 9.0 8.5 8.0 7.5 7.0 6.5 6.0 5.5 5.0 4.5 4.0 3.5 3.0 2.5 2.0 1.5 1.0 ppm

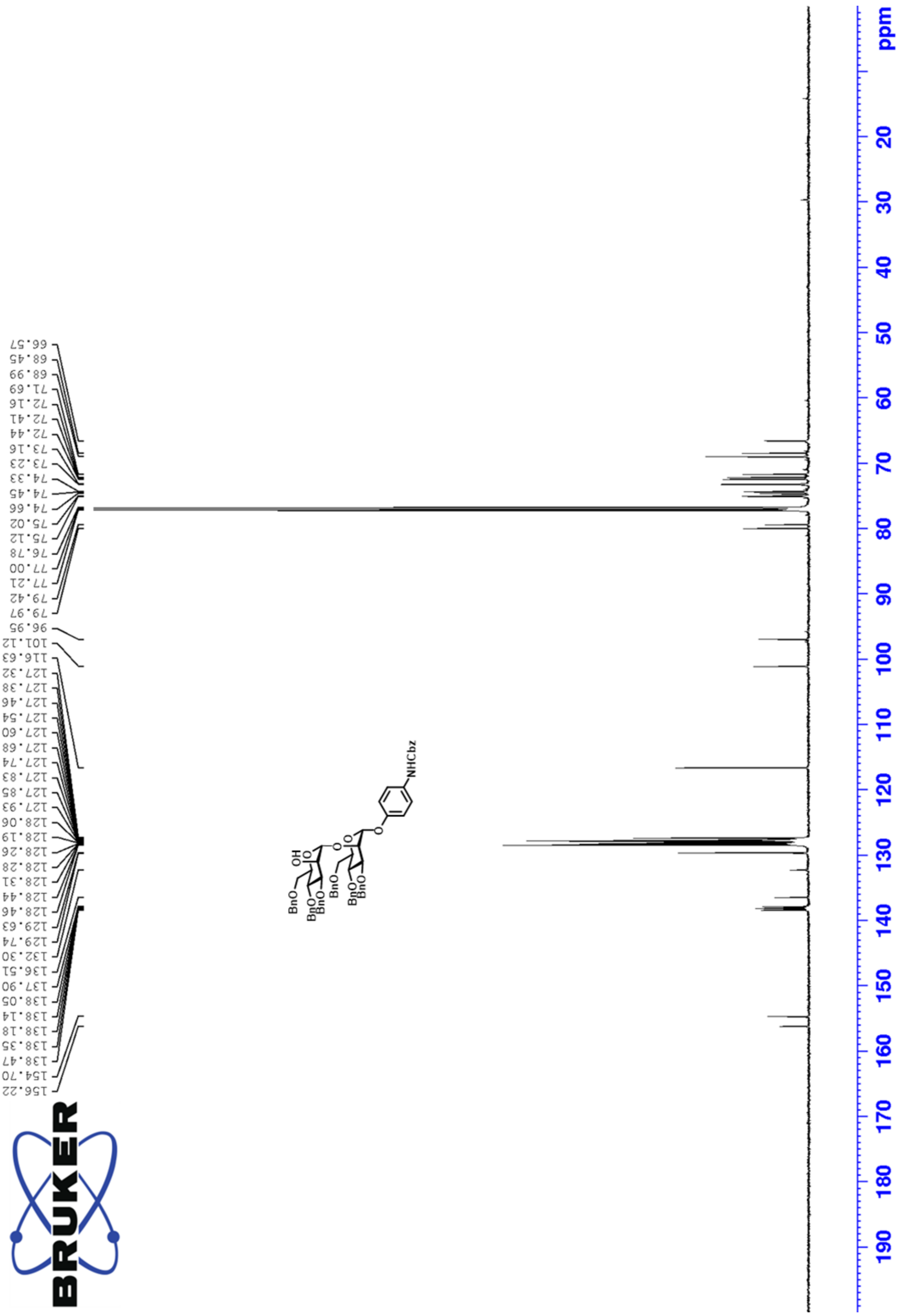
66.57
68.45
68.99
71.69
72.16
72.41
72.44
73.16
73.23
74.33
74.45
74.66
75.02
75.12
76.78
77.00
77.21
79.42
79.97
96.95
101.12
116.63
127.32
127.38
127.46
127.54
127.60
127.68
127.74
127.83
127.85
127.93
128.06
128.19
128.26
128.28
128.31
128.44
128.46
129.63
129.74
132.30
136.51
137.90
138.05
138.14
138.18
138.35
138.47
154.70
156.22
NHCbz
BnO
OH
BnO
BnO
BnO
BnO
BnO
ppm
20
30
40
50
60
70
80
90
100
110
120
130
140
150
160
170
180
190

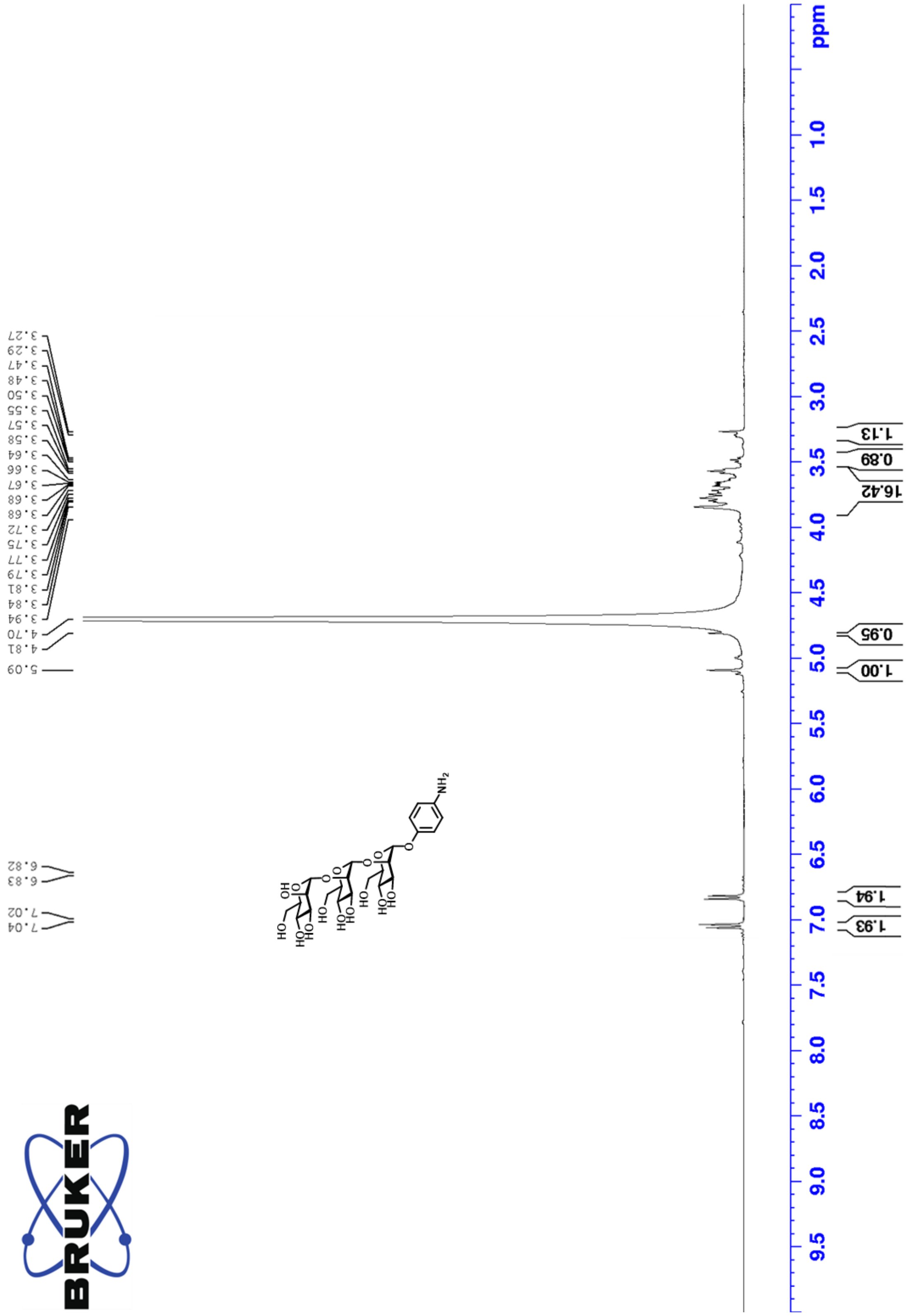
3.27
3.29
3.47
3.48
3.50
3.55
3.57
3.58
3.64
3.66
3.67
3.68
3.68
3.72
3.75
3.77
3.79
3.81
3.84
3.94
4.70
4.81
5.09
6.82
6.83
7.02
7.04
1.13
0.89
16.42
0.95
1.00
1.94
1.93
ppm
1.0
1.5
2.0
2.5
3.0
3.5
4.0
4.5
5.0
5.5
6.0
6.5
7.0
7.5
8.0
8.5
9.0
9.5
NH2

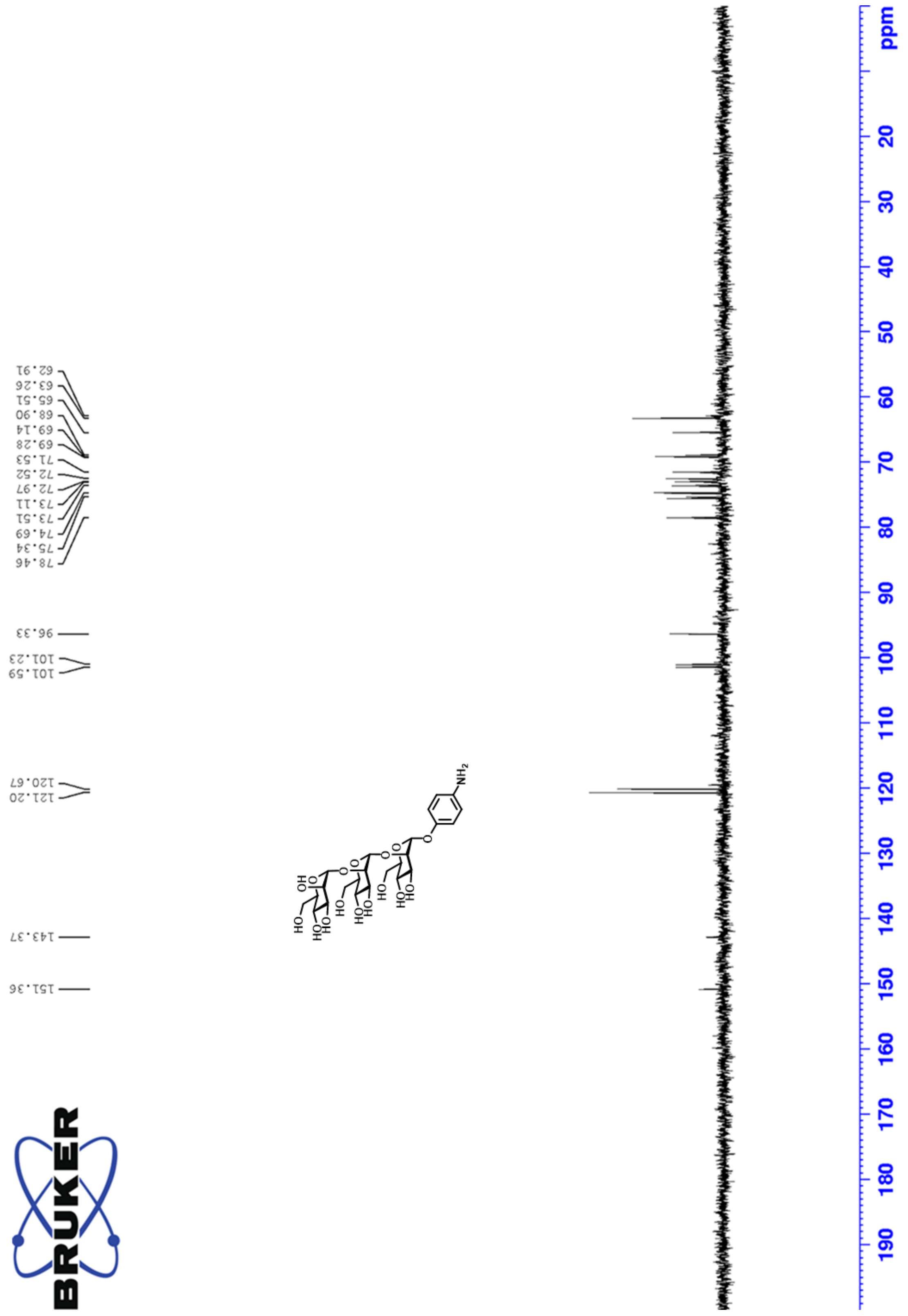
62.91
63.26
65.51
68.90
69.14
69.28
71.53
72.52
72.97
73.11
73.51
74.69
75.34
78.46
96.33
101.23
101.59
120.67
121.20
143.37
151.36
NH2
HO
OH
O
ppm
20
30
40
50
60
70
80
90
100
110
120
130
140
150
160
170
180
190

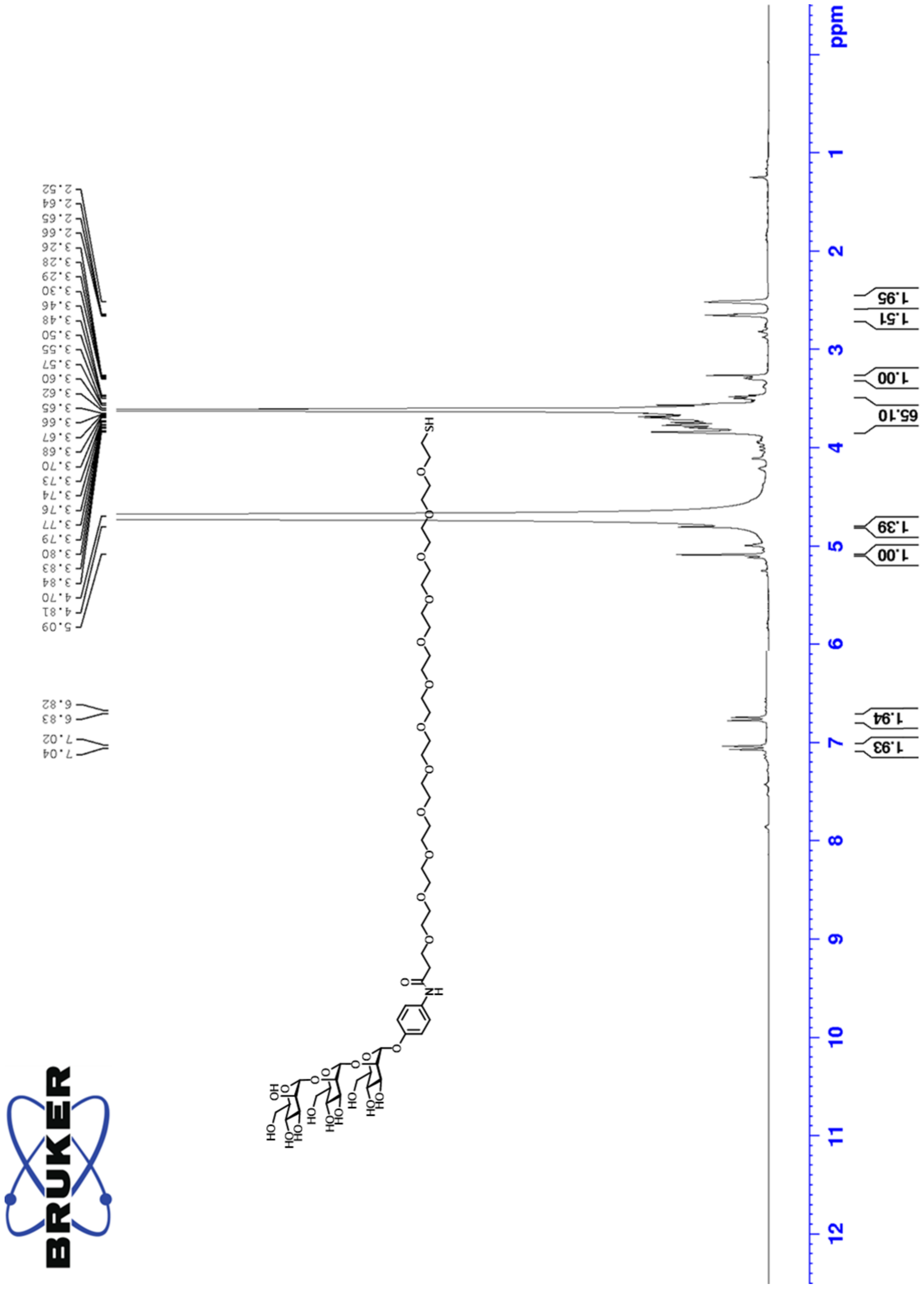
2.52
2.64
2.65
2.66
3.26
3.28
3.29
3.30
3.46
3.48
3.50
3.55
3.57
3.60
3.62
3.65
3.66
3.67
3.68
3.70
3.73
3.74
3.76
3.77
3.79
3.80
3.83
3.84
4.70
4.81
5.09
6.82
6.83
7.02
7.04
1.95
1.51
1.00
65.10
1.39
1.00
1.94
1.93
ppm
1
2
3
4
5
6
7
8
9
10
11
12

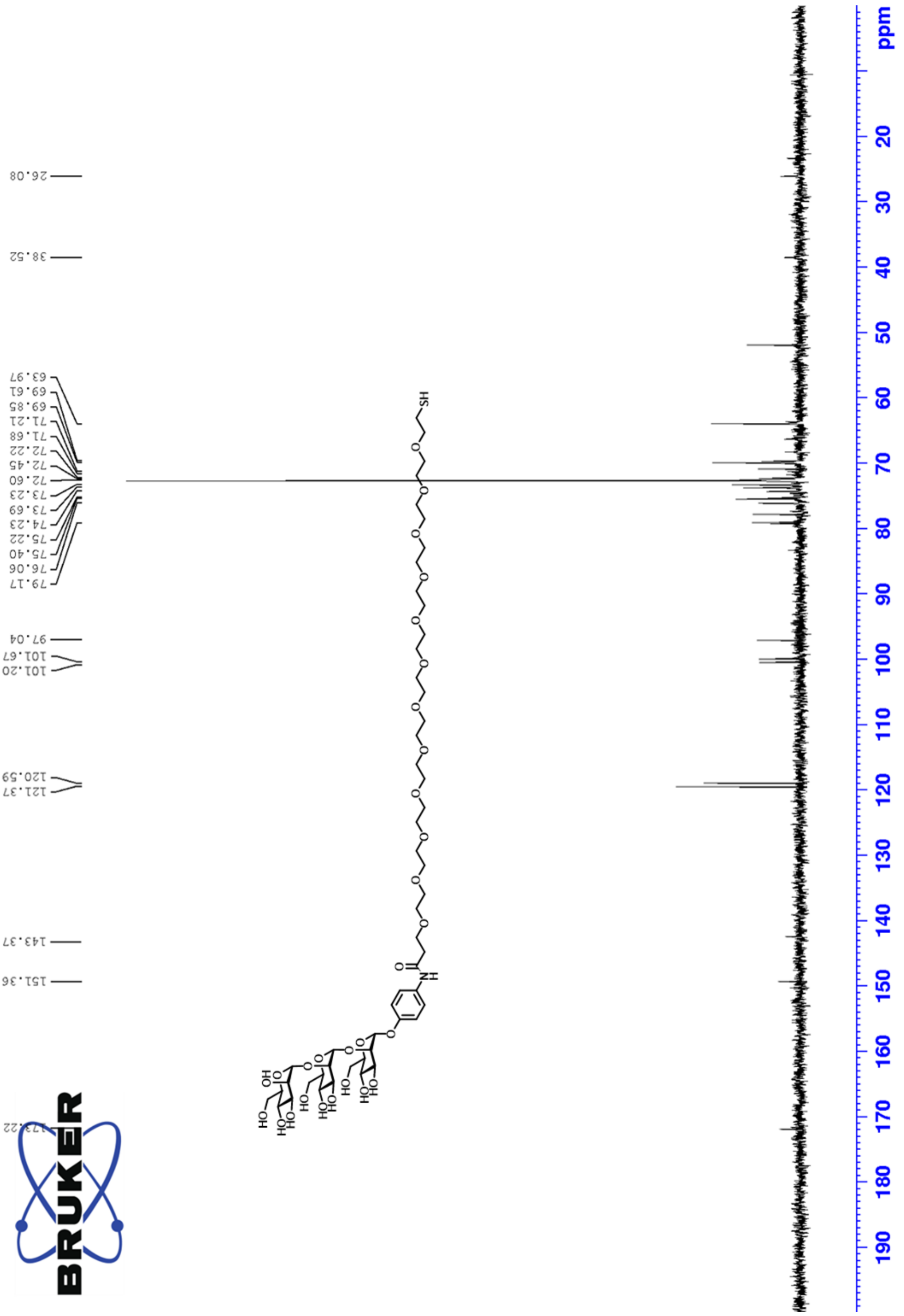
26.08
38.52
63.97
69.61
69.85
71.21
71.68
72.22
72.45
72.60
73.23
73.69
74.23
75.22
75.40
76.06
79.17
97.04
101.67
101.20
120.59
121.37
143.37
151.36
173.22
BRUKER
ppm
20
30
40
50
60
70
80
90
100
110
120
130
140
150
160
170
180
190
